# Dynamics of the HD regulatory subdomain of PARP-1; substrate access and allostery in PARP activation and inhibition

**DOI:** 10.1101/2020.09.04.283309

**Authors:** Tom E.H. Ogden, Ji-Chun Yang, Marianne Schimpl, Laura E. Easton, Elizabeth Underwood, Philip B. Rawlins, Michael M. McCauley, Marie-France Langelier, John M. Pascal, Kevin J. Embrey, David Neuhaus

## Abstract

PARP-1 is a key early responder to DNA damage in eukaryotic cells. An allosteric mechanism links initial sensing of DNA single-strand breaks by PARP-1’s F1 and F2 domains *via* a process of further domain assembly to activation of the catalytic domain (CAT); synthesis and attachment of poly(ADP-ribose) (PAR) chains to protein sidechains then signals for assembly of DNA repair components. A key component in transmission of the allosteric signal is the HD subdomain of CAT, which alone bridges between the assembled DNA-binding domains and the active site in the ART subdomain of CAT. Here we present a study of isolated CAT domain from human PARP-1, using NMR-based dynamics experiments to analyse WT apo-protein as well as a set of inhibitor complexes (with veliparib, olaparib, talazoparib and EB-47) and point mutants (L713F, L765A and L765F), together with new crystal structures of the free CAT domain and inhibitor complexes. Variations in both dynamics and structures amongst these species point to a model for full-length PARP-1 activation where first DNA binding and then substrate interaction successively destabilise the folded structure of the HD subdomain to the point where its steric blockade of the active site is released and PAR synthesis can proceed.

## INTRODUCTION

Poly(ADP-ribose) polymerase 1 (PARP-1) is a highly abundant, chromatin-associated protein that is a key early responder to genomic stress in eukaryotes (1). Upon sensing DNA damage, particularly single-strand breaks (SSBs) that are the commonest form of lesion (2), it becomes strongly activated, catalysing addition of poly(ADP-ribose) (PAR) to nearby proteins, including itself, and thereby signalling for assembly of downstream DNA repair factors (3,4). Inhibition of PARP enzymes has emerged as an important route to cancer therapy, since the combined effects of PARP inhibition and defective homologous recombination (HR) selectively kill BRCA-deficient tumour cells, whereas healthy cells remain largely unaffected (5,6). This is an example of a “synthetic lethality”, so-called because it results from the cumulative effects of losing two complementary repair pathways simultaneously; similar effects involving PARP inhibition in conjunction with other tumour-associated repair defects have also been found (7,8). There is an emerging consensus that the toxic effects of PARP inhibition in tumour cells result from inhibitor-bound PARP being retained, or trapped, on DNA lesions, thereby blocking replication and repair, but the underlying molecular mechanisms responsible for such trapping have so far been elusive (9).

Previous studies have shown that initial recognition of DNA SSBs by PARP-1 is achieved by the two N-terminal zinc finger domains F1 and F2 (Figure 1A) (10-13), which upon binding at the break co-operate to bend and twist the DNA into a conformation that is inaccessible to intact double-stranded DNA (14). This DNA binding initiates an assembly cascade that forms a network of domain-domain interactions, first seen in the context of double-strand break binding without F2 (15), in which the initially independent F1, F2, F3 and WGR domains gather on the damage site, and in so doing juxtapose the WGR and F3 domains to create a composite interface that interacts with the regulatory HD subdomain of CAT (Figure 1D) (14,15). Remarkably, it is the interaction of the HD subdomain with this composite interface contributed by WGR and F3, rather than any direct contact between the CAT domain and the DNA, that constitutes a key part of the DNA-dependent activity switch in PARP-1. It is clear from the architecture of the complex that the HD subdomain forms a structural bridge between, on the one side, the assembled F1, F2, F3 and WGR domains on the DNA, and on the other, the ART subdomain, implying that the HD transmits the activation signal to the active site (Figure 1B,C). Initially it was suggested that this might be achieved by DNA-dependent distortions of the HD domain structure (15). However, later work highlighted the importance of dynamics; an elegant HXMS study showed that DNA-binding by full-length PARP-1 leads to a significant increase in solvent exposure for parts of the HD subdomain (Figure 1D) (16). It was proposed in the same study that this correlates with increased local dynamics within the HD, and that this in turn is responsible for opening access to the enzyme active site, which is auto-inhibited by the HD in un-activated PARP-1. Overall, the role of DNA binding in PARP-1 activation is thus to cause the WGR and F3 domains to be held together in the correct arrangement for them jointly to create the appropriate “landing pad” for the HD, and it is the resulting interaction of HD with WGR and F3 that supplies the free energy required to modulate the internal dynamics of the HD (14,15). In the free protein, where the mutual relationship of WGR and F3 is spatially unconstrained (17), any transient interactions that may occur between HD and either WGR or F3 are insufficient to cause activation.

**Figure 1:**
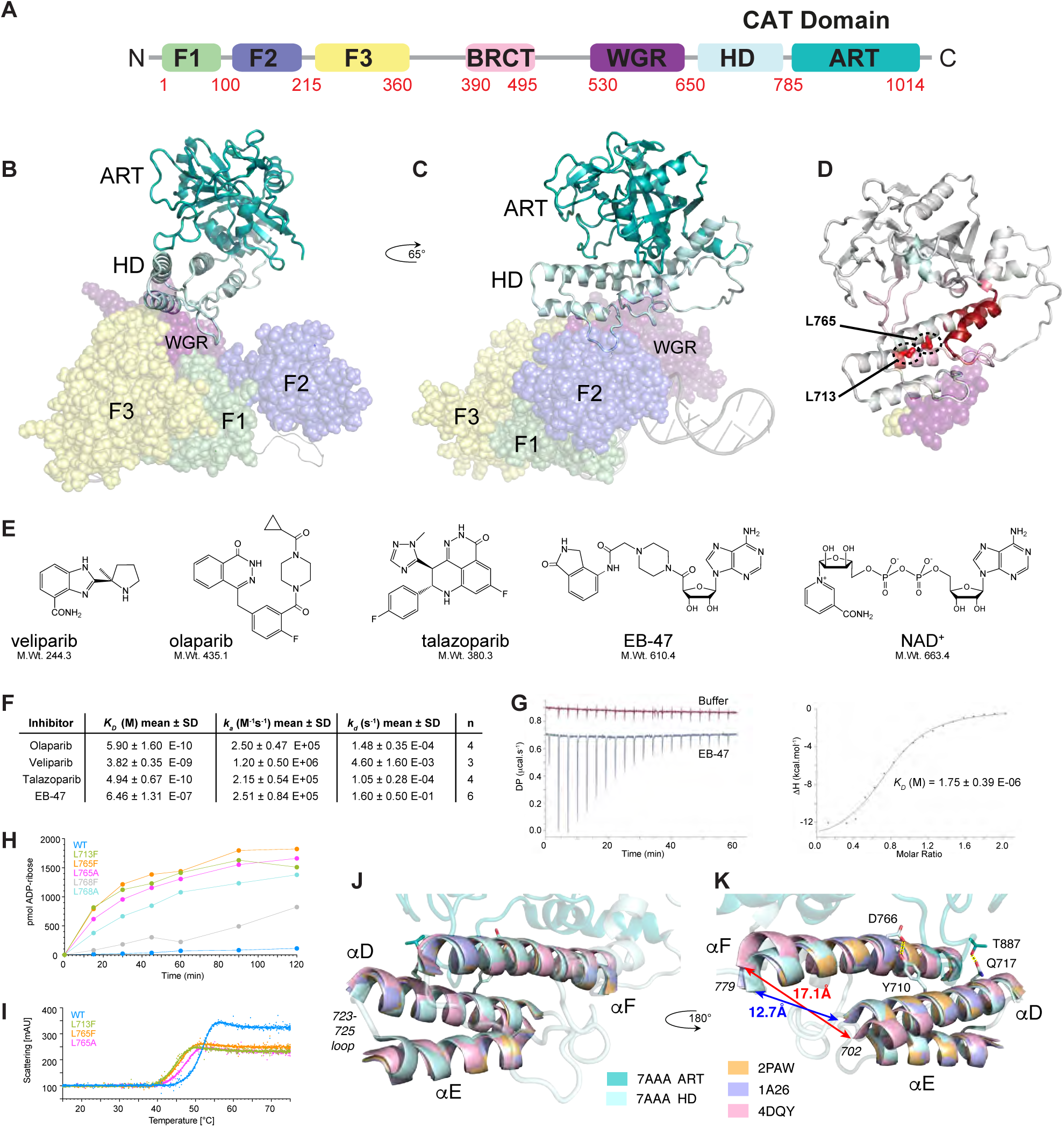
A) PARP-1 domain structure. B, C) Model of PARP-1 bound to a DNA single-strand break, showing how the HD subdomain acts as a bridge between the DNA-interacting domains (F1, F2, F3 and WGR, shown as semi-transparent spheres) and the ART subdomain (shown as cartoon; the BRCT domain and interdomain linkers are not represented). The model was built by combining co-ordinates from PDB 4DQY (F1, F3 and WGR-CAT bound to a DNA blunt end) and PDB 2N8A (F1 and F2 bound to a 45-nucleotide DNA dumbbell that mimics a single-strand break) as described previously (14). D) PARP-1 CAT domain showing locations of the HD subdomain mutations studied here (L713 and L765), the parts of the surfaces of the WGR and F3 domains with which the HD subdomain interacts (shown as semi-transparent spheres), and a summary of the previously published HXMS data (16) showing NH exchange rate changes upon DNA binding for the CAT domain in the context of full-length PARP-1; progressively darker shades of red indicate progressively greater increases in NH exchange upon PARP-1 binding to the 45nt DNA dumbbell (see Dawicki-McKenna *et al*. (16) for a complete description). E) Covalent structures of the four inhibitors studied here (veliparib, olaparib, talazoparib and EB-47) and PARP-1’s natural substrate, NAD^+^. F) Binding affinities, association and dissociation rates for interaction of veliparib, olaparib, talazoparib and EB-47 with full-length PARP-1, measured using surface plasmon resonance. G) Isothermal calorimetry determination of *K*_*D*_ for the binding of PARP-1 CAT domain with EB-47. H) Catalytic activity of isolated PARP-1 CAT domain and mutants, tested using a colorimetric assay that measures the incorporation of ADP-ribose into PAR using biotinylated NAD^+^ (22). I) Thermal melt data for WT PARP-1 CAT domain and the L713F, L765F and L764A mutants, measured using nanoDSF (differential scanning fluorimetry). J, K) Superpositions of PDB 1A26, PDB 2PAW and PDB 4DQY onto the structure of PDB 7AAA, using the backbone N, C*α*, C’ atoms of helices D, E and F; for all four molecules, helices D, E and F are shown as solid, while the remainder of the structure is shown for 7AAA only and is semi-transparent. Changes caused by DNA binding (in 4DQY) include particularly a realignment of helix D and straightening of helix F. The H-bonds linking Tyr710 to Asp766 (at the site of the kink in helix F) and Thr887 to Gln717 are indicated (shown for 7AAA only), as is the position of the 723-725 loop.

While dynamics were clearly implicated by these findings, at the start of this study the nature of the internal dynamics of the HD subdomain, the ways in which these changed during activation and their relevance to trapping, remained largely unclear. Earlier work had characterised a series of mutants, including several in the HD subdomain that have an elevated rate of PAR synthesis relative to wild-type PARP-1 in the absence of binding to DNA damage (15). We hypothesised that the higher basal catalysis rates in these mutants likely reflect changes in their HD subdomains that might partially mimic those caused by DNA-damage-dependent activation, and that given these effects are seen in the absence of DNA, where interactions with F3 and WGR are not significant, they could be studied in the isolated CAT domain. Similarly, we reasoned that studying the effects of adding inhibitors to isolated CAT domain could lead to insights into the relevance of HD dynamics to trapping. PAR is a highly negatively charged, branched polymer built from units of ADP-ribose derived from NAD^+^, and it has long been argued that dissociation of PARP from DNA damage sites is strongly facilitated by electrostatic repulsion between the DNA and the elongating PAR chains attached to PARP itself (3). Prevention of polymer production must therefore be a key contribution to trapping of inhibitor-bound PARP complexes, but more recently it has been suggested that binding of certain inhibitors may superpose further effects relevant to trapping by altering the chain of interactions linking the enzyme active site to the DNA-binding domains, thereby changing the affinity of the complex for the DNA damage site (7,18). Such “reverse allostery” would necessarily involve changes to the properties of the HD subdomain, and again these changes would most likely involve dynamics. Such inhibitor-induced changes would also be expected to be relevant to understanding activation. The great majority of PARP inhibitors mimic only the nicotinamide moiety of PARP’s natural substrate NAD^+^, and it has previously been shown that a non-hydrolysable analogue of NAD^+^ (benzamide adenine dinucleotide, BAD), which includes also the adenosine moiety, is incapable of binding to isolated PARP-1 CAT domain unless the HD subdomain is deleted (19); in addition, a series of inhibitors designed to reach into the adenosine binding pocket were all similarly unable to bind to the isolated, intact CAT domain (20). In light of such observations, the autoinhibitory blockade of catalysis caused by the HD subdomain, and which is relieved during activation, is hypothesised to be primarily a steric barrier to binding of the adenosine moiety of NAD^+^. It is therefore highly relevant to investigate effects seen for larger inhibitors that go some way towards mimicking the adenosine moiety of NAD^+^, while still being compatible with binding, albeit more weakly, to the intact, isolated CAT domain. An example of exactly such a case is the inhibitor EB-47 (21), which is included in our study.

In this study we have sought to uncover the structural nature of dynamic changes that underlie the regulatory role of the HD subdomain of PARP-1. To this end, we have investigated a variety of mainly NMR-based comparisons across a set of inhibitor complexes and point mutants of the isolated PARP-1 CAT domain, in particular using amide group NH chemical shifts, ^15^N relaxation parameters and solvent exchange characteristics, and in addition we have determined X-ray crystallographic structures of the apo-form of human PARP-1 CAT domain, as well as its complexes with inhibitors used in this study where crystal structures were unavailable at the start of the work.

## MATERIALS AND METHODS

### Plasmids and mutagenesis

The DNA construct used for human PARP-1 (656-1014 with a V762A substitution, hereafter referred to as PARP-1 CAT domain) was codon-optimised for expression in *E. coli* and incorporated into a pET28 vector also containing a sequence for N-terminally His_6_-tagged *Geobacillus stearothermophilus* di-hydrolipoamide acetyltransferase (UniProt P11961) lipoyl-binding domain; the resultant protein contained a TEV cleavage site between the lipoyl-binding and PARP-1CAT domains. Constitutively partially active mutants (L713F, L765F, L765A) of this fragment were cloned from this vector using a QuikChange II site-directed mutagenesis kit (Agilent). The DNA construct used for proteins for crystallography was similar except it coded for PARP-1 662-1011 (again with V762A) and was cloned into pET24a vector containing N-terminal Avi- and hexahistidine tags followed by a TEV cleavage site.

### PARP-1 activity assay

Poly(ADP-ribose) catalysis was measured using a previously described assay (22) in which hexahistidine-tagged PARP-1 was immobilized on Ni(II)-NTA-coated plates (Qiagen) in the presence of NAD^+^/biotinylated-NAD^+^ (Trevigen). The amount of ADP-ribose produced over time was quantified based on streptavidin-horseradish peroxidase (Pierce) interaction with biotinylated poly(ADP-ribose) and utilization of the Ultra-TMB chromogenic substrate (Pierce). PARP-1 catalytic domains (wild-type and mutants, see Figure 1H) were purified as described (23).

### Protein expression and purification

All protein fragments for NMR and biophysics experiments were expressed using *E. coli* BL21 (DE3) cells. Unlabelled PARP-1 CAT domain used for biophysical experiments was expressed in 2x TY supplemented with 50 µg ml^-1^ kanamycin; protein synthesis was induced with 2 mM IPTG once OD_600_ reached 0.8 and continued overnight at 30°C. Samples of PARP-1 CAT domain uniformly labelled with ^15^N or with [^2^H,^15^N,^13^C], or specifically labelled with ^15^N at single residue types (lysine, arginine, leucine or isoleucine) were expressed in M9 minimal media as described previously (24). For samples of the CAT domain point-mutants, expression media were supplemented with 10 mM benzamide (Sigma-Aldrich) and expression performed overnight at 20°C. Purification was carried out as described previously for NMR samples of PARP-1 CAT domain (24).

Protein for crystallography was expressed in *E. coli* Gold BL21 (DE3) and purified by immobilised metal affinity chromatography with Ni^2+^-NTA resin, proteolytic cleavage of the affinity tag and size exclusion chromatography. PARP1 was concentrated to 24-37 mg/ml in a final buffer containing 50 mM Tris pH 8.0, 150 mM NaCl and 1 mM TCEP. For SPR measurements, human PARP-1 (NCBI accession AAH37545, residues 2-1014) was cloned into a pFastBac expression vector to include an N-terminal 6His-6Lys tag followed by a TEV cleavage site. P1 virus was generated and protein expressed following standard Invitrogen protocols. PARP-1 was purified by an IMAC affinity purification step followed by a Size Exclusion Chromotography step using an AKTA system. The final protein was stored at -80^°^C in 50 mM KH_2_PO_4_ pH7.8, 150 mM KCl, 5 mM 2-mercaptoethanol and 10% glycerol.

### NMR spectroscopy

All NMR measurements employed in-house Bruker Avance III 600 MHz or Avance III HD 800 MHz spectrometers or the Avance III HD 950 MHz spectrometer at the MRC Biomedical NMR Centre, all equipped with 5 mm [^1^H,^13^C,^15^N]-cryogenic probes. All NMR samples (except those used in the real-time ^2^H_2_O exchange series) were prepared using 50 mM [^2^H_11_] Tris, pH 7.0, 50 mM NaCl, 2 mM [^2^H_10_] DTT and 0.02% (w/v) NaN_3_ in 95:5 H_2_O/^2^H_2_O (NMR Buffer). Samples of PARP-1 CAT domain in complex with PARP inhibitors for NMR were made by adding inhibitors (50mM in [^2^H_6_] DMSO, Sigma-Aldrich) to a final ratio of 1:1.5 (veliparib, olaparib and talazoparib) or 1:1.6 (EB-47); these relatively large nominal excesses of ligand were used to ensure that the protein would be fully saturated even if there were moderately large errors in quantification of the components. NMR data were processed using the programmes MddNMR (25), NMRPIPE (26), and TopSpin (Bruker BioSpin GmbH), and analysed using the programmes CcpNMR Analysis 2.4.2 (27) and Sparky version 3.115 (28). The assignment process for the backbone amides of wild-type PARP-1 (656-1014) has been described previously (24). Starting from those assignments, backbone amide NH signals for PARP-1 CAT domain mutants and inhibitor complexes were assigned by careful comparison of 3D [^15^N,^1^H]-HSQC-NOESY spectra (29) (*τ*_*m*_=70 ms) recorded at 25 °C; ^15^N-labelled protein concentrations were 0.3 mM (veliparib complex), 0.4 mM (olaparib and EB-47 complexes, WT apo-protein and point mutants) or 0.5 mM (talazoparib complex). Chemical shift perturbations (CSPs) were calculated from ^15^N and ^1^H chemical shifts of backbone amides from [^15^N-^1^H]-TROSY spectra using the formula (30)

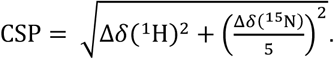

CSP values were mapped onto structures of wild-type catalytic domain (7AAA) and veliparib (7AAC), olaparib (7AAD), talazoparib (4UND) and EB-47 (7AAB) complexes in Pymol by globally normalising values to CSP_max_ the largest CSP value across all of the samples; scaled CSP values were converted into a grey-to-yellow colour ramp using a home-written script. In order to avoid the mapped structural plots being dominated by the relatively uninformative largest CSP values, the colour scale was set to run from 0 to 0.2(CSP_max_), with CSP values above 0.2(CSP_max_) being uniformly represented as yellow.

### ^15^N relaxation experiments

Backbone ^15^N longitudinal relaxation times (*T*_*1*_), spin-locked relaxation times (*T*_*1ρ*_) and ^15^N{^1^H} steady-state NOE values were determined for 0.4 mM ^15^N-labelled samples of WT PARP-1 CAT domain, CAT domain point-mutants and inhibitor complexes at 25 °C using pulse sequences described by Lakomek et al. (31). For the *T*_*1*_ measurements the delays used for all samples were 0, 120, 120, 280, 480, 800, 1200, 1200, 2000, 3200, 4800 and 6800 ms while for the *T*_*1ρ*_ measurements they were 0.15, 2, 4, 4, 7, 12, 20, 34, 34, 50, 70 and 100 ms (the duplicate values were used to check reproducibility). Relaxation times and errors were calculated using the non-linear least squares fitting routine in Sparky. Steady-state {^1^H}^15^N NOE values were measured using a saturation time of 7 s, and errors were derived as previously described (32). Values of *τ*_*c*_ for backbone amides in all samples were calculated from *T*_*1*_ and *T*_*1ρ*_ values using the formula

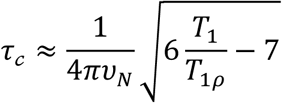

where *v*_*N*_ is the ^15^N resonance frequency in Hz (33).

### NH solvent exchange experiments

CLEANEX-PM experiments (34) were recorded for 0.4 mM ^15^N-labelled samples of WT PARP-1 CAT domain, CAT domain point-mutants and inhibitor complexes at 25 °C using a mixing time of 150 ms. In order to facilitate comparisons between intensities in CLEANEX-PM spectra from different samples, they were each approximately normalized using a factor calculated for each sample as

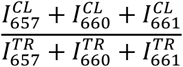

where 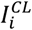 represents the intensity of residue *i* in the CLEANEX-PM spectrum and 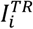 represents the intensity of the corresponding peak in a 400 μM ^15^N TROSY reference spectrum (residues Val657, Gly660 and Thr661 were chosen to be used since they give well-resolved signals and, being in the unstructured extreme N-terminus of the protein, give high intensity in CLEANEX-PM while being unlikely to be affected by inhibitor binding or the HD subdomain mutations).

Slowly exchanging backbone amide NH protons were detected using a real-time series of 16 [^15^N-^1^H]-TROSY experiments starting from lyophilised PARP-1 CAT domain. Samples were prepared by initially snap-freezing in dry ice 0.5 mL, 0.1 mM aliquots of ^15^N-labelled WT PARP-1 CAT domain in (H_2_O-based) NMR buffer and lyophilising overnight (>16 hours). Pellets were then re-suspended in 0.5 mL ^2^H_2_O (Sigma-Aldrich) and the samples immediately transferred to 5 mm NMR tubes (Norell). For the inhibitor complexes, 1.5 uL of 50 mM PARP inhibitor stock solubilised in 100% ^2^H_6_-DMSO was added immediately after re-suspension to give a final inhibitor concentration of 0.15 mM (a 1:1.5 ratio). Sixteen TROSY spectra were then recorded continuously over a 48-hour period, and peak heights from each measured in CCPN analysis 2.4 (Vranken et al., 2005). For further analysis, peak intensities at the 3-hour, 12-hour and 39-hour time points were corrected and normalised between the spectra to account for differences in sample concentration and number of scans. In a control experiment, WT PARP-1 CAT domain was subjected to exactly the same lyophilisation procedure except that the protein was resuspended in H_2_O rather than ^2^H_2_O; the TROSY spectrum of this sample was essentially unchanged relative to that of the protein prior to lyphilisation (Figure S1). Data from the CLEANEX and ^2^H_2_O real-time exchange experiments were mapped onto the relevant structures for display in Pymol version 1.8.6.2 (35). Fast exchanging NH signals detected in CLEANEX experiments are coloured red, while slowly exchanging NH signals delected in the real-time TROSY series are coloured according to approximate exchange rates estimated from the intensities of the 3, 12 and 39 hour time points as follows: light blue: [I(3h) – I(12h) > 0.5 x I(3h)]; medium blue: [I(3h) – I(12h) < 0.5 x I(3h) & I(12h) – I(39h) > 0.25 x (I12h)]; dark blue: [I(3h) – I(12h) < 0.5 x I(3h) and I(12h) – I(39h) < 0.25 x (I12h)]. Residues for which no signal was observed in CLEANEX or in the real-time TROSY series are coloured white.

### Biophysical measurements

Thermal stabilities of 0.5 mM samples of WT PARP-1 CAT domain and the L713F, L765F and L765A point mutants in NMR buffer were assessed by nanoDSF using a Prometheus NT.48 instrument (Nanotemper). Samples were heated from 15 °C to 75 °C on a gradient increasing by 2 °C per minute at 30% excitation.

For surface plasmon resonance (SPR) measurements with full-length PARP-1, a Biacore T200 or 8K instrument was used to monitor binding interactions utilising a Biacore NTA Series S Sensor Chip (Cytiva). 20 mM HEPES, 150 mM NaCl, 1 mM TCEP and 0.05% Tween-20, 1% (v/v) buffer was used for both immobilisation and binding at 25 °C. A flow rate of 30 μL/min. was used for binding analysis. 2 μg/mL protein was immobilized using Nickel-NTA capture-coupling (36) resulting in approximately 2000 RU of immobilised PARP-1 protein. Prior to kinetic analysis, solvent calibration and double referencing subtractions were made to eliminate bulk refractive index changes, injection noise and data drift. Affinity and binding kinetic parameters were determined by global fitting to a 1:1 binding model within the Biacore Evaluation or Insight Software (Cytiva).

Isothermal titration calorimetry (ITC) experiments were performed on a MicroCal iTC200 calorimeter (Malvern) with 20 µM PARP-1 CAT domain in the cell and 200 µM EB-47 in the pipette, both in ITC buffer (50 mM sodium phosphate, 50 mM NaCl, 0.5 mM TCEP). EB-47 powder was initially solubilised in [^2^H_6_] DMSO, and the [^2^H_6_] DMSO concentration in both cell and pipette was matched at a final concentration of 2% (v/v). Experiments were conducted at 25°C, with an initial injection of 0.5 µL followed by 19 injections of 2 µL with 180 second intervals between each injection. A series of control injections of 200µM EB-47 into ITC buffer only was also performed to determine the heat change from dilution. Data was fitted to a single class binding site model using MicroCal PEAQ-ITC Analysis Software to determine *K*_*D*_.

Circular dichroism (CD) spectra were measured using a Jasco J-815 spectrometer, scanning in the range of 260 to 190 nm at 20°C. Samples of wild-type PARP-1 CAT domain and CAT domain mutants were dialysed overnight into CD buffer (50 mM sodium phosphate pH 7.4, 50 mM NaCl, 0.1 mM TCEP). For complexes of wild-type PARP-1 CAT domain with veliparib, olaparib and talazoparib, the inhibitor was dissolved in [^2^H_6_] DMSO and added to the protein to reach a protein:inhibitor ratio of 1:1.5 so as to ensure saturation, then dialysed into fresh buffer 3 times to remove [^2^H_6_] DMSO. For the complex of wild-type PARP-1 CAT domain with EB-47, the inhibitor was solubilised in CD buffer and added directly to protein, again to reach a protein:inhibitor ratio of 1:1.5.

### X-ray crystallographic structure determination

PARP-1 (662-1011) was crystallised in hanging drop vapour diffusion in 2.45 M ammonium sulphate and 0.1 M Tris pH 8.5. Protein pre-incubated for 30 minutes with 1 mM olaparib was crystallised under the same conditions. The veliparib complex was obtained by soaking crystals for 24 hours with 2mM compound in reservoir solution containing 2 % DMSO. Cryoprotection comprised a brief immersion in a solution of 2.6 M ammonium sulphate, 5 % DMSO, 5 % ethylene glycol and 0.1 M Tris pH 8.5. The structure in complex with EB-47 was obtained by co-crystallisation; 0.85 mM PARP-1 (662-1011) was mixed with 1 mM EB-47 (from a 100 mM stock in DMSO) and crystallised at 20 °C in of 0.4 M ammonium sulphate, 25 % PEG 3350, 0.1 M PCTP buffer (sodium propionate, sodium cacodylate, and Bis-Tris propane) pH 5.5. X-ray diffraction data were collected at Diamond Light Source beamlines I03 and I04. Data processing was carried out with autoPROC (37) using XDS (38), POINTLESS (39) and AIMLESS (40) and the CCP4 suite (41). Anisotropy correction was performed with STARANISO (42). Structures were solved by molecular replacement with AMoRe (43) or Phaser (44); and refined using Buster (45) and Coot (46). Initial ligand geometry restraints were generated using Grade (47). Data collection and refinement statistics are summarised in Table 1.

**Table 1.** Crystallographic data collection and refinement.

|  | **PARP1**<br>***apo*** | **PARP1**<br>**veliparib** | **PARP1**<br>**olaparib** | **PARP1**<br>**EB-47** |
| --- | --- | --- | --- | --- |
| **PDB ID** | 7AAA | 7AAC | 7AAD | 7AAB |
| **Data collection** |  |  |  |  |
| DLS beamline | I03 | I03 | I04 | I03 |
| Wavelength (Å) | 0.9762 | 0.97625 | 0.9795 | 0.97623 |
| Space group | P2<sub>1</sub>2<sub>1</sub>2<sub>1</sub> | P2<sub>1</sub>2<sub>1</sub>2<sub>1</sub> | P2<sub>1</sub>2<sub>1</sub>2<sub>1</sub> | P6<sub>1</sub> |
| Cell dimensions |  |  |  |  |
| *a*, *b*, *c* (Å) | 48.6 | 48.2 | 48.0 | 134.9 |
|  | 92.5 | 91.6 | 91.6 | 134.9 |
|  | 163.5 | 162.0 | 162.1 | 111.3 |
| $\alpha$, $\beta$, $\gamma$ (°) | 90 | 90 | 90 | 90 |
|  | 90 | 90 | 90 | 90 |
|  | 120 | 90 | 90 | 90 |
| Resolution (Å) | 48.59-1.74 | 81.02-1.59 | 81.07-2.21 | 116.83-2.80 |
|  | (1.79-1.74) | (1.73-1.59) | (2.33-2.21) | (3.05-2.80) |
| *I* / σ*I* | 10.1 (1.2) | 10.8 (1.6) | 9.1 (1.5) | 7.3 (1.5) |
| CC 1/2 | 0.997(0.527) | 0.998 (0.657) | 0.996 (0.809) | 0.992 (0.501) |
| Completeness (%) | 99.9 (99.9) | 92.5 (73.8) | 89.8 (79.0) | 93.2 (51.2) |
| Redundancy | 6.4 (6.1) | 6.3 (4.3) | 9.9 (9.6) | 10.3 (10.4) |
| **Refinement** |  |  |  |  |
| Resolution (Å) | 48.59-1.74 | 65.00-2.31 | 81.07-2.21 | 116.83-2.80 |
| No. reflections | 489209 | 422964 | 259439 | 202538 |
| *R*<sub>work</sub> / *R*<sub>free</sub> | 0.204/0.227 | 0.190 / 0.222 | 0.208/0.254 | 0.181 / 0.215 |
| No. atoms |  |  |  |  |
| Protein | 5456 | 5485 | 5464 | 5393 |
| Inhibitor |  | 36 | 64 | 78 |
| Buffer | 27 | 10 | 15 | 20 |
| Water | 358 | 482 | 105 | 21 |
| *B*-factors |  |  |  |  |
| Protein | 32.3 | 23.9 | 39.2 | 66.2 |
| Ligand |  | 11.6 | 27.2 | 47.9 |
| Buffer | 35.1 | 23.8 | 53.8 | 91.5 |
| Water | 33.3 | 28.3 | 26.1 | 27.4 |
| R.m.s. deviations |  |  |  |  |
| Bond lengths (Å) | 0.01 | 0.01 | 0.01 | 0.01 |
| Bond angles (°) | 1.04 | 1.04 | 1.14 | 1.22 |

Values in parentheses are for the highest resolution shell.

## RESULTS

### X-ray Crystal Structure of Human PARP-1 CAT Domain Apo Protein

In order to establish an appropriate reference for the comparisons in this study, we determined an X-ray crystal structure of the isolated CAT domain (residues 662-1011) of human PARP-1 (PDB 7AAA); data collection and refinement statistics appear in Table 1. Although there are many published crystal structures of this domain bound to various inhibitors, and a structure of un-liganded PARP-1 CAT domain from chicken (PDB 2PAW) was published in 1996 (48,49), to our knowledge there has been no previous report of a crystal structure of the human apo-protein. In accordance with multiple reports of crystal soaking with inhibitors for the generation of small molecule complex structures, we found that the CAT domain crystallises readily in the absence of any ligands. Upon solving the apo structure, we observed weakly defined electron density in the nicotinamide pocket, which was attributed to a molecule of DMSO from the cryoprotection process during crystal freezing.

Comparison of this new structure of the unliganded human CAT domain to the previously determined corresponding chicken protein structure 2PAW reveals that the backbone conformations of the ART subdomain in the two structures are near-identical, whereas those of the HD subdomain (and residues 779-788 in the HD-ART linker) show somewhat greater differences (Figure 1J,K). Helix F shows a very mild bend in the new structure, whereas in 2PAW the distortion in this helix is slightly stronger and takes the form of a kink near residue Asp766. In addition, several of the helices of the HD are slightly differently arranged, resulting in small differences between their relative orientations and positions between the two structures; this is particularly marked for helix E, the helix most distant from the ART domain.

It is also of interest to compare this new un-liganded structure of the isolated CAT domain from human PARP-1 to the corresponding domain as seen in the complex of the F1, F3, WGR and CAT domains of human PARP-1 bound to a model DNA double-strand break (PDB 4DQY) (15). In that complex, DNA-dependent contacts to the WGR and F3 domains cause distortions of the HD subdomain, which were previously analysed by comparing the conformation in the DNA-bound complex with that seen in the isolated CAT domain of the chicken protein. Although the chicken protein structure originally used for this comparison was that of a complex with carbaNAD (PDB 1A26) (50), the protein backbone conformation in that complex is essentially identical to that in 2PAW, presumably because the 2-carba-NAD molecule binds relatively superficially and does not penetrate substantially into the CAT domain active site. Comparison of the new human CAT domain apo-protein structure to 4DQY shows essentially the same differences involving leucines 698 and 701 near the very short helix C that were previously reported. These changes go together with shifts in the mutual arrangement of the helices, particularly for D and F (see also Tables S1-S3 for tables of superposition statistics for individual helices and combinations of helices); helix F is mildly bent in the isolated CAT domain apo-protein structures but becomes straight upon DNA binding, while the axis of helix D shifts causing the N-terminus of D and the C-terminus of F to move further apart (the distance Ser702 C*α* - Arg779 C*α* increases from 12.7Å to 17.1Å). Differences within the ART subdomain between the DNA-bound and apo-structures are much smaller than those in the HD, but there is a very localised difference involving residues Pro885 - Tyr889 that come close to helix D of the HD subdomain. This could be linked to possible DNA-modulation of an interaction between the sidechains of Thr887 and Gln717 that might be relevant to the movement of helix D; the OH group of Thr887 in the new apo-protein structure (though probably not in the chicken protein structures 2PAW or 1A26) is well-placed to form a hydrogen bond to the sidechain carbonyl of Gln717, whereas in the DNA-bound structure this hydrogen bond is absent.

The overall extent of these differences amongst the various structures is captured in the rmsd statistics collected in Tables 2, S4 and S5, which underline the contrast between the close similarity amongst the ART domain conformations in all cases and the greater divergence amongst the conformations and relative dispositions of the HD subdomains. To demonstrate the latter the most clearly we have included rmsds calculated in an unconventional way, using two different sets of atoms; set 1 is used to carry out the fitting of the molecules to one another, following which the rmsd is calculated using the co-ordinates of set 2 without moving the molecules away from the positions defined by the fitting already carried out using set 1. Whereas conventional rmsd values (i.e. calculated with set 1 = set 2) for the HD or ART subdomains capture only internal differences within each subdomain, these unconventional rmsds (i.e. calculated with set 1 ≠ set 2) are mainly sensitive to changes in their relative disposition.

**Table 2.** Superposition statistics for the HD and ART subdomains in structures of PARP-1 CAT domain.

| **Fitted atoms (set 1)**<br>**Measured atoms (set 2)**<br>(all fits are to 7AAA chain A) | **Resolution**<br>(Å) | **Space group** | **CAT**<br>**CAT**<br>(Å) | **ART**<br>**ART**<br>(Å) | **HD**<br>**HD**<br>(Å) | **ART**<br>**HD**<br>(Å) | **HD**<br>**ART**<br>(Å) |
| --- | --- | --- | --- | --- | --- | --- | --- |
| 7AAA chain B (apo-protein) | 1.74 | P 2<sub>1</sub>2<sub>1</sub>2<sub>1</sub> | 0.508 | 0.191 | 0.710 | 1.015 | 1.113 |
| 2PAW chain A (apo-protein) | 2.30 | P 2<sub>1</sub>2<sub>1</sub>2<sub>1</sub> | 0.679 | 0.568 | 0.630 | 1.041 | 0.976 |
| 1A26 chain A (2-carba-NAD complex) | 2.25 | P 2<sub>1</sub>2<sub>1</sub>2<sub>1</sub> | 0.681 | 0.496 | 0.646 | 1.189 | 1.033 |
| 4DQY chain C (F1,F3,WGR-CAT:DNA) | 3.25 | P 2<sub>1</sub>2<sub>1</sub>2<sub>1</sub> | 1.293 | 0.804 | 1.566 | 2.233 | 2.328 |
| 7AAC chain A (veliparib complex) | 1.59 | P 2<sub>1</sub>2<sub>1</sub>2<sub>1</sub> | 0.145 | 0.133 | 0.120 | 0.181 | 0.389 |
| 2RD6 chain A (veliparib complex) | 2.30 | P 3 2 1 | 0.980 | 0.855 | 0.541 | 1.720 | 1.550 |
| 7AAD chain A (olaparib complex) | 2.21 | P 2<sub>1</sub>2<sub>1</sub>2<sub>1</sub> | 0.241 | 0.231 | 0.220 | 0.272 | 0.456 |
| 4UND chain A (talazoparib complex) | 2.20 | P 2<sub>1</sub>2<sub>1</sub>2<sub>1</sub> | 1.174 | 0.760 | 0.850 | 2.429 | 2.855 |
| 4PJT chain A (talazoparib complex) | 2.35 | P 2<sub>1</sub>2<sub>1</sub>2<sub>1</sub> | 1.185 | 0.690 | 0.847 | 2.574 | 3.042 |
| 7AAB chain A (EB-47 complex) | 2.8 | P 6<sub>1</sub> | 1.789 | 0.506 | 0.969 | 4.175 | 4.362 |
| 6VKQ chain A (EB-47 complex) | 2.9 | P 4<sub>1</sub>2<sub>1</sub>2 | 1.421 | 0.771 | 0.997 | 3.166 | 2.763 |

The atoms used for superposition were as follows:
CAT = 666-721, 730-743, 750-779, 790-936, 939-1009 (N,C<sup>α</sup>,C');;
HD = 666-721, 730-743, 750-779 (N,C<sup>α</sup>,C');;
ART = 790-936, 939-1009 (N,C<sup>α</sup>,C');;
Set 1 atoms are used to calculate transformed co-ordinates resulting from fitting;
Set 2 atoms are used to calculate an rms difference without further changing the co-ordinates.

### Inhibitor Complexes and Mutants

We next used NMR spectroscopy to characterise the isolated CAT domain of wild-type PARP-1 (residues 662-1014) both free and in complex with the inhibitors veliparib, olaparib, talazoparib and EB-47 (see Figure 1E), and in addition three CAT domain mutants (L713F, L765F and L765A) that, based on previous studies, are known to be constitutively partially active in the full-length context (15). These inhibitors were chosen since they largely span the range of binding affinities and cytotoxicities of PARP inhibitors currently in the clinic or clinical trial, while EB-47 was selected because, unlike the others, it includes a moiety that partially mimics the adenosine moiety of NAD^+^, the natural substrate of PARP-1 (Figure 1E). Binding data for all four inhibitors were obtained using SPR (Figure 1F), and in the case of the EB-47 complex also using ITC (Figure 1G; ITC measurements for the more tightly-binding inhibitors were considered unreliable due to probable complications from non-equilibrium binding). The mutants of HD subdomain residues were selected from amongst those previously characterised as having the largest enhancements of basal PAR synthesis compatible with the yield and solution stability required for NMR experiments. Figure 1H shows the results of a colourimetric automodification assay for WT PARP-1 and a series of HD subdomain mutants (L713F, L765F, L765A, L768F and L768A), demonstrating that the mutations used in the present study cause constitutive enhancements of basal PARylation activity by factors of approximately 20-fold. The two mutation sites are located near the centres of helices D and F, facing one another but offset by one helical turn (Figure 1D). Thermal melting curves for the isolated CAT domain mutants (Figure 1I) showed that their unfolding transitions occurred at lower temperatures than for WT CAT domain (the fitted value of T_onset_ for WT PARP-1 CAT domain was 42.5 °C, whereas those for the corresponding L713F, L765F and L765A CAT domain mutants were respectively 34.8 °C, 35.5 °C and 36.7 °C). Test experiments revealed that in order to limit precipitation during experiments NMR data acquisition for the CAT domain mutants needed to be carried out at 25 °C, so in order to facilitate comparisons amongst the various spectra this temperature was adopted for all NMR experiments (even though the WT CAT domain could tolerate somewhat higher temperatures for data acquisition).

NMR signal assignments were obtained for the backbone amide resonances of WT CAT domain predominantly by using globally [^15^N, ^13^C, ^2^H] labelled protein in conjunction with a standard suite of triple resonance NMR experiments. However, given the high degree of overlap in the spectra of this 360-residue species, and the fact that many C^β^ and also some C^α^ signals were missing from these data (presumably due to high transverse relaxation rates, particularly for the carbon resonances), it was necessary to augment these results using NOE-based experiments as well as spectra from samples labelled with ^15^N in specific amino-acid types only (Lys, Arg, Leu, Ile, Gly+Ser). Overall, this approach yielded backbone amide group assignments for 331 (96.5%) of the 343 non-proline residues (Figures S1B and S1C). Notably, we could not assign any signals corresponding to residues Ala823 - Asn827 in this or indeed any of the other species in this study, suggesting the presence of intermediate-rate dynamics (see below). We found that although deuteration of the CAT domain was necessary to allow the triple resonance experiments used for signal assignment to succeed, it was not needed for observation of amide signals using TROSY experiments. NMR experiments with the mutants and the complexes with inhibitors therefore employed [^15^N] singly-labelled protein, and signal assignments for these species were made by careful comparison against TROSY spectra of free WT CAT domain in conjunction with NOE-based experiments. The inhibitor complexes with veliparib, olaparib and talazoparib all exhibited slow exchange on the NMR chemical shift timescale, as would be expected given their binding affinities, and there was no evidence for any weak secondary binding sites (spectra obtained at a 1:1 inhibitor:protein ratio were indistinguishable from those obtained in presence of a 50% excess of inhibitor). In the case of the complex with EB-47 the spectra were of markedly lower quality, which may have been caused in part by internal dynamics within the complex (see below). This restricted somewhat the number of assignments that could be made for this complex, particularly in the ART subdomain, but nonetheless backbone amide group assignments were made for 302 (88.0%) non-proline residues. For the other inhibitor complexes, the corresponding totals of assigned non-proline backbone amide signals were: veliparib 328 (95.6%), olaparib 327 (95.3%), talazoparib 324 (92.4%), while for the mutants they were: L713F 323 (94.2%), L765F 326 (95.0%) and L765A 328 (95.6%).

### Chemical Shift Comparisons

Analysis of chemical shift differences between amide group NH signals of free WT CAT domain and those of the inhibitor complexes showed, as expected, that chemical shift perturbations (CSPs) were dominated by direct effects of inhibitor binding (Figure 2A-D). The largest contributions to these CSPs would be expected to come from magnetically induced ring-currents within the aromatic ring systems of the inhibitors. These cause large chemical shift changes for nearby nuclei, but can be non-trivial to quantify accurately, particularly for ring current contributions caused by relatively uncommon or multiply-fused aromatic ring systems. As discussed in later sections, crystal structures were determined or previously available for all of the inhibitor complexes used in this study, which facilitated interpretation since these structures allowed spatial relationships of individual protein amide NH groups to aromatic ring systems of bound inhibitors to be visualised. With this in mind, it is clear that some moderately-sized CSPs are detected at protein residues that are sufficiently distant from the bound inhibitor that direct effects are very unlikely, implying that these effects most likely result from conformational adjustments in the protein following inhibitor binding; in particular, this seems clearly to be the case for some residues within helices of the HD subdomain. The sizes of such effects range from those in the veliparib complex, which shows only negligible CSPs in parts of the HD distant from the inhibitor, through those for the olaparib complex, which shows appreciable effects only in helix F (that may in part be due to ring current effects), to those for the talazoparib and EB-47 complexes that show more widely distributed CSPs in the HD subdomain, particularly in helices C, D and F.

**Figure 2:**
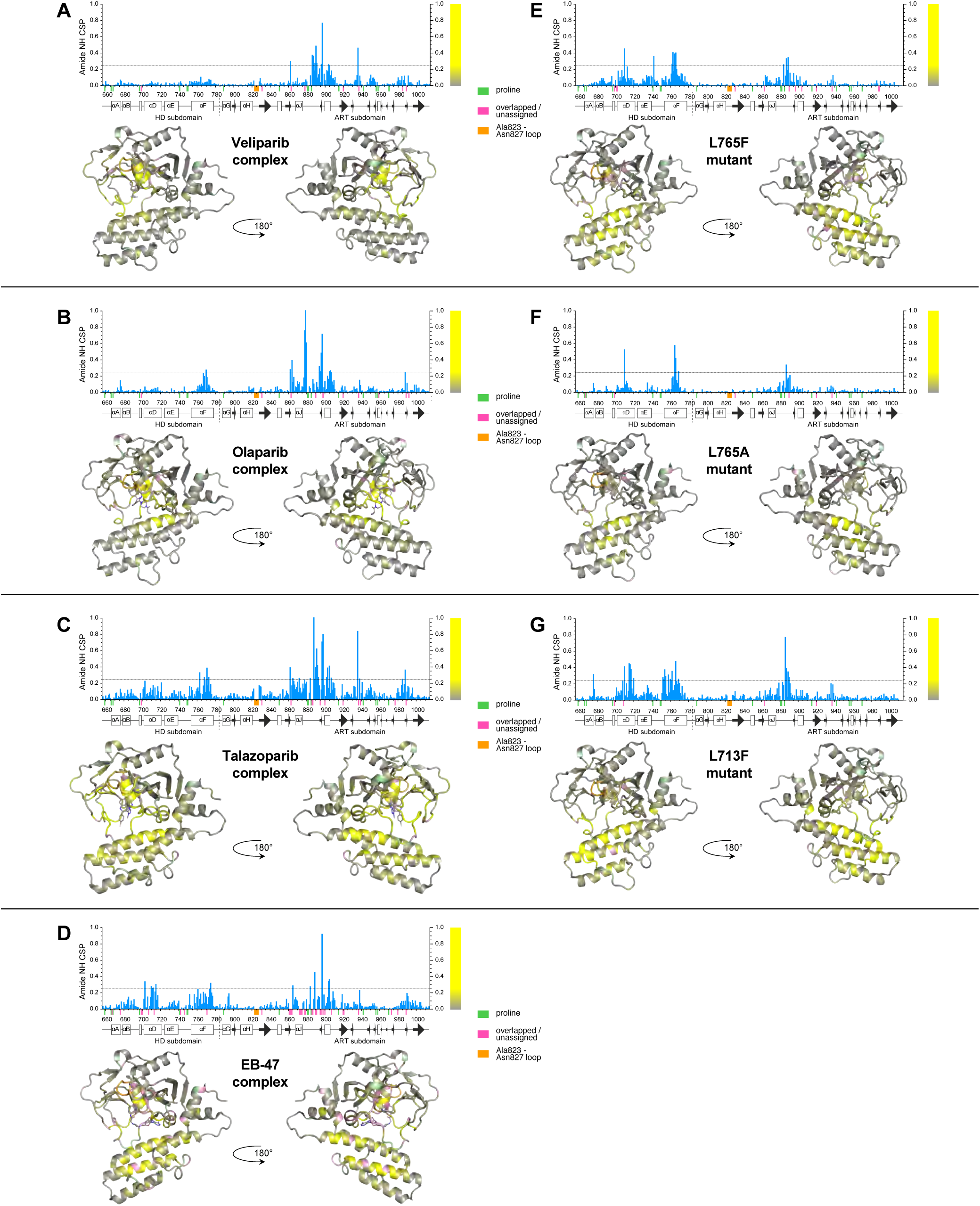
Backbone amide group chemical shift perturbations for complexes of PARP-1 CAT domain with A) veliparib, B) olaparib, C) talazoparib, D) EB-47, and backbone amide group chemical shift differences between WT PARP-1 CAT domain and the mutants E) L765F, F) L765A and G) L713F, in each case shown both as histograms and mapped to the relevant CAT domain complex crystal structure or (for the mutants) the WT protein crystal structure. The largest difference in any of the datasets is 1.247ppm (for G888 in the talazoparib complex, shown truncated in the figure). The bar to the right of each histogram shows the colour code used to map these CSP values to the structures: values between 0 and 0.249 (0.2 times the largest CSP value, shown with a horizontal line in the histograms) are shown using a colour ramp running from grey to yellow, while values >0.249 are uniformly shown as yellow; this approach was followed to prevent the mapping being excessively dominated by a small number of the largest differences. Small coloured bars beneath the sequence scale and matching colours on the structures are used to indicate the positions of prolines (pale green), overlapped or unassigned signals (pink) and the Ala823-Asn827 loop for which no signals were seen in any spectrum (orange). Secondary structure elements are shown beneath the histograms.

As commented above, for the EB-47 complex (but not the others in this study), many of the signals were significantly broadened upon complexation, and this was particularly the case amongst signals from the ART subdomain. One likely reason for this is that the adenosine moiety in EB-47 may not be fixed in a unique orientation relative to the protein, but rather may adopt a variety of conformations e.g. with different *χ* angles, such that the adenine aromatic system would exert markedly different ring current effects on signals from neighbouring nuclei in the protein in the different conformations. The two available crystal structures of the EB-47 complex provide clear evidence that such conformational differences can indeed occur (see below). This would result in substantial broadening of protein signals from nuclei in the vicinity of the adenine ring if such conformations interconvert at an intermediate rate on the chemical shift timescale (that is, *k* ≈ Δ*f* where *k* is the interconversion rate in sec^-1^ and Δ*f* is the frequency difference expressed in Hz between corresponding signals due to a given nucleus in two interconverting conformers). The broadening is unlikely to arise from exchange between the free and bound states, as the protein binding site should be approx. 99% saturated under our experimental conditions, given the measured *K*_*D*_ value.

In the case of the mutants (Figure 2E-G), CSPs were observed mainly in the spatial vicinity of the mutations, i.e. for parts of helices D, E and F and the nearby ART domain loop. As would be expected, larger and more extensive changes were seen close to the mutation sites for the L765F and L713F mutants, where new aromatic ring current effects must result from introduction of Phe residues, than were seen for L765A; this makes the comparison of shifts between WT and mutant more straightforward to interpret in the case of L765A. Interestingly, one of the largest CSPs in all three mutants occurs for the NH group of Tyr710. The apo-protein crystal structure, as well as 2PAW, 1A26 and 4DQY and the crystal structures of all the inhibitor complexes in this study (see below), show that in all cases the sidechain OH of Tyr710 probably forms a hydrogen bond to the sidechain carboxylate of Asp766 (see Figure 1K), and they further show that Asp766 is located at precisely the point where helix F is prone to form a kink in several of the structures (2PAW, 1A26 and the two EB-47 complex structures; see below). While a shift difference at the NH of Tyr710 in the L765F and L713F mutants could at least in part be caused by ring current effects from the introduced Phe residues, this is not the case for the L765A mutant. It seems plausible, therefore, that a large chemical shift difference relative to WT for the NH of Tyr710 may correlate with kinking of helix F centred on Asp766, implying that such kinking may be present in at least the L713F mutant, and possibly also the L765F and L765A mutants; it is difficult to see how other forms of helical re-arrangement or adjustment would easily result in a significant chemical shift change that was so localised to a very small number of residues. For the ART subdomain, NH chemical shift changes in the mutants relative to WT are largely restricted to the region Ala880 - Met890. The fact that these changes are clearly seen in the L765A mutant shows that they cannot be wholly due to ring currents caused by addition of a Phe residue, suggesting that this loop in the ART subdomain is sensitive to relative movements of helices D and F in the HD subdomain, with which it forms contacts.

Overall, these chemical shift differences relative to the WT apo-protein spectrum, both in the inhibitor complexes and in the mutants, are consistent with structural adjustments occurring within the folded structure of the CAT domain. In the case of the inhibitor complexes, the CSP data alone are not well-suited to assessing whether significant structural changes occur in the ART subdomain, since the CSPs are dominated by direct effects caused by proximity to the inhibitors; however, analysis of the crystal structures of these complexes (see below) shows that in fact the backbone conformations of the ART domains are substantially unperturbed upon inhibitor binding. For the mutants, the CSPs indicate that there are no appreciable changes in the ART domain outside of the few residues in loops that contact the HD directly. In contrast, the chemical shift data for residues in the HD subdomain strongly suggest that, both for the inhibitor complexes and for the mutants, there are appreciable differences in protein conformation relative to the WT apo-protein. The data indicate that in general these are not large-scale changes, such as full or partial melting of entire regions that would be expected to lead to new signals at “random coil” chemical shifts, and/or gross changes in lineshapes. Rather, the chemical shift changes are consistent with more limited structural differences, such as relatively subtle alterations to the mutual packing of helices, movements of the HD subdomain relative to the ART, changes in loop conformations or changes in helical bending.

### ^15^N Relaxation analysis

We next carried out ^15^N relaxation experiments for the WT human PARP-1 CAT domain, the mutants and the inhibitor complexes, in each case acquiring *T*_*1*_, *T*_*1ρ*_ and heteronuclear ^15^N{^1^H} steady-state NOE data. As a first step in the analysis, we used the *T*_*1*_ and *T*_*1ρ*_ data to calculate approximate rotational correlation times, *τ*_*c*_, for each residue (33). The rotational correlation time is a global property, but by calculating it on a per-residue basis and comparing averaged values for core secondary structural elements in the HD subdomain with similarly obtained values for the ART subdomain we sought to establish whether these two regions of the CAT domain showed evidence for independent motion. The results were consistent across the WT apo-protein, the mutants and the inhibitor complexes (Figures S2 and S3); in each case they showed that, both for the HD subdomain and for the ART subdomain, the approximate rotational correlation time was in the region of 25 ns, as would be expected for a single globular protein domain of approximately 40kDa (33). This demonstrates that the two subdomains tumble in solution as a single unit, showing that if there is any independent motion of the two subdomains, it occurs on a timescale slower than overall tumbling.

Local variations in the ^15^N relaxation parameters generally reflect regions having flexibility on a timescale comparable to or faster than overall molecular tumbling (itself characterised by the 25ns value of *τ*_*c*_ described above). Of the parameters measured in this study, the steady-state ^15^N{^1^H} heteronuclear NOE data reveal the sites of local mobility within the CAT domain structure the most clearly (Figure 3A); values of approximately 0.8 are expected for regions lacking flexibility on this fast timescale, while lower values correlate with progressively more extensive local motions of the corresponding amide NH groups relative to the rigid core of the protein domain and appear as dips in the value of the heteronuclear NOE in the histograms in Figure 3 (31,51,52). Mapping of these data for the WT apo-protein onto the crystal structure shows that the regions of highest local mobility correspond to surface loops, but notably such loops are not uniformly mobile. In the HD subdomain, the residues that show the clearest indication of local flexibility are Asp678 (between helices A and B), Gly723 - Ser725 (between helices D and E), Phe744 - Lys748 (which form part of the linker between helices E and F) and Gly780 - Asp788 (which form the linker to the ART subdomain). Of these, only the 744-748 loop is known to form significant interactions with another domain (WGR) in the DNA-bound complex 4DQY (15), suggesting that the flexibility of that particular loop might be quenched in the full-length context upon DNA damage recognition; the others are likely to remain flexible in the context of full-length PARP-1 bound to DNA. In the ART domain, a number of relatively flexible sequence regions lie spatially close together in locations at or near contacts that would be present in the recently described complex of PARP CAT domain with the accessory protein HPF1 (24); these include particularly residues Met890 - Phe891, His934 - Leu941 and Asn979 - Thr981. In addition, the loop Ala823 - Asn827, which consistently fails to show NH signals in any of the spectra of any of the species studied here, also lies within the HPF1 binding region, as does the extremely flexible C-terminal region, Phe1009 - Trp1014. The loss of signals from the Ala823 - Asn827 loop is most likely due to conformational exchange at an intermediate rate on the chemical shift timescale (i.e. *k*_*ex*_ ≈ Δ*δ*, likely approx. 10-1000 s^-1^), suggesting that these are likely to be the slowest of the motions considered in this section; the observation of only a single set of signals in the other regions discussed implies that they are probably in the fast exchange regime.

**Figure 3:**
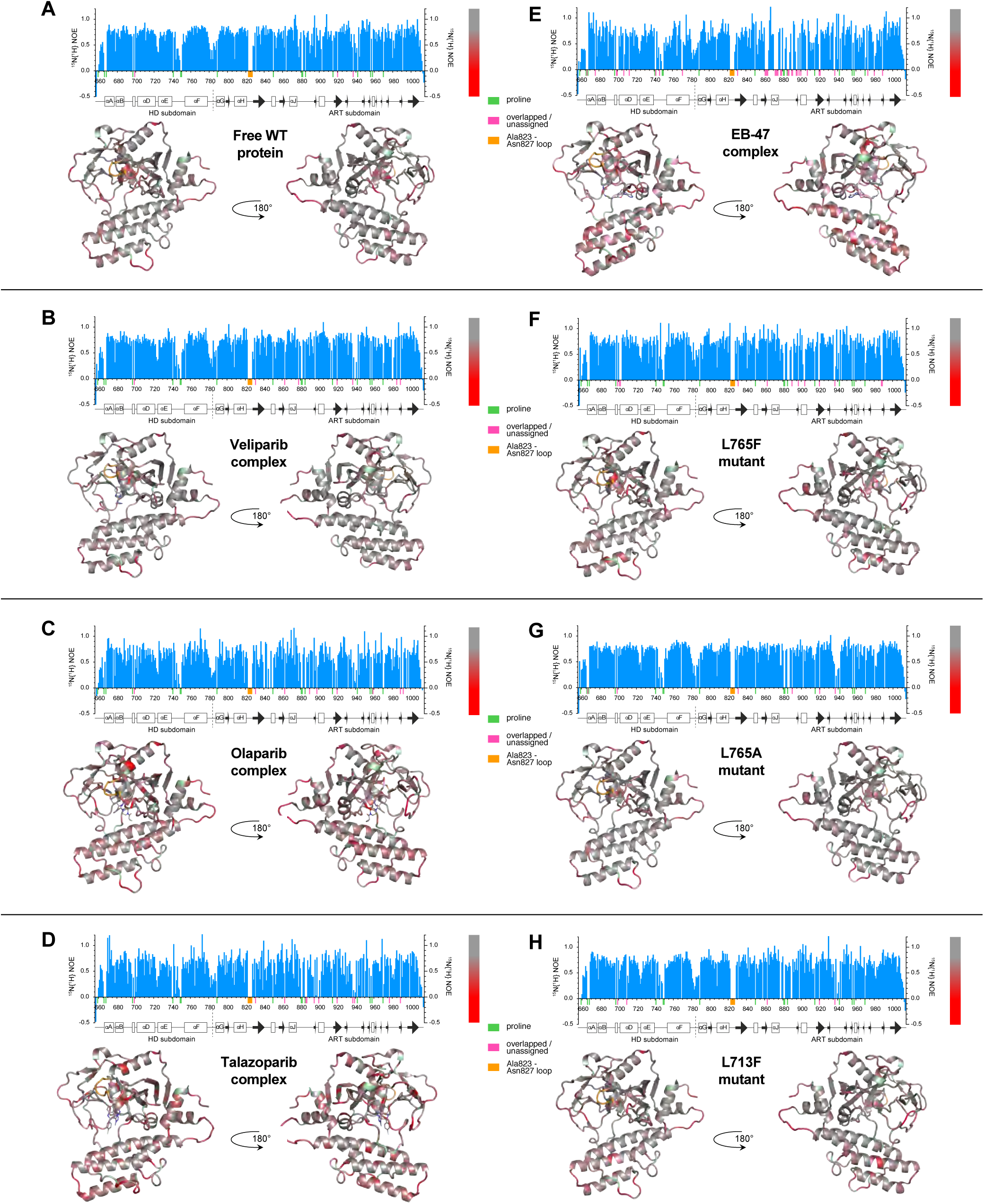
Steady-state {^1^H} ^15^N NOE data for A) PARP-1 CAT domain, its complexes with B) veliparib, C) olaparib, D) talazoparib and E) EB-47, as well as the point mutants F) L765F, G) L765A and H) L713F. In each case, the data are shown both as histograms and mapped to the relevant CAT domain crystal structure or (for the mutants) the WT protein crystal structure. The bar to the right of each histogram shows the colour code used to map these data to the structures: the colour ramp runs from grey (0.8, the approximate value expected for NH groups that show no internal motions faster than overall tumbling of the protein) to red (0.0), with values above 0.8 uniformly grey and values below 0.0 uniformly red. Small coloured bars beneath the sequence scale and matching colours on the structures are used to indicate the positions of prolines (pale green), overlapped or unassigned signals (pink) and the Ala823-Asn827 loop for which no signals were seen in any spectrum (orange). Further histograms of steady-state {^1^H}^15^N NOE, *T*_*1*_ and *T*_*1ρ*_ data appear in Figures S4-S15.

The pattern seen for the WT apo protein is very largely repeated for each of the inhibitor complexes and mutants (Figure 3B-H), although in a few cases, particularly for residues of the ART domain in the inhibitor complexes, the fact that data could not be obtained for as many residues as in the WT apo-protein limited the interpretation somewhat. In the case of the L713F mutant, some additional flexibility relative to WT was observed for residues Tyr710 and Ser711, whereas for the two mutants of Leu765 no changes were seen in the immediate vicinity of the mutation. Corresponding plots of the *T*_*1*_ and *T*_*1ρ*_ data appear in Figures S4 – S15, and they each show similar trends.

Collectively, these results from analysis of ^15^N relaxation data are consistent with those obtained from the chemical shift analysis, namely that structural differences between the WT apo CAT domain and either the inhibitor complexes or the mutants do not involve large-scale changes in substantial portions of the structure or in fast-timescale dynamics processes. Further, they suggest that whatever differences in dynamics may or may not exist on slower timescales within the CAT domains in these various species, they are generally not reflected in the fast processes sampled by these ^15^N relaxation experiments.

### Amide proton exchange

We therefore turned next to NMR measurements of amide NH proton solvent exchange (Figures 4 and 5), which report primarily on the strength of hydrogen bonding. Since an amide NH proton that participates in an intramolecular hydrogen bond is protected against exchange with solvent while that hydrogen bond remains intact, NH solvent exchange rates are strongly influenced by the frequency of transient “opening” events during which hydrogen bonds are broken, which in turn reflects local flexibility (53-55). Some relevant background and theory appears in Figure S16.

**Figure 4:**
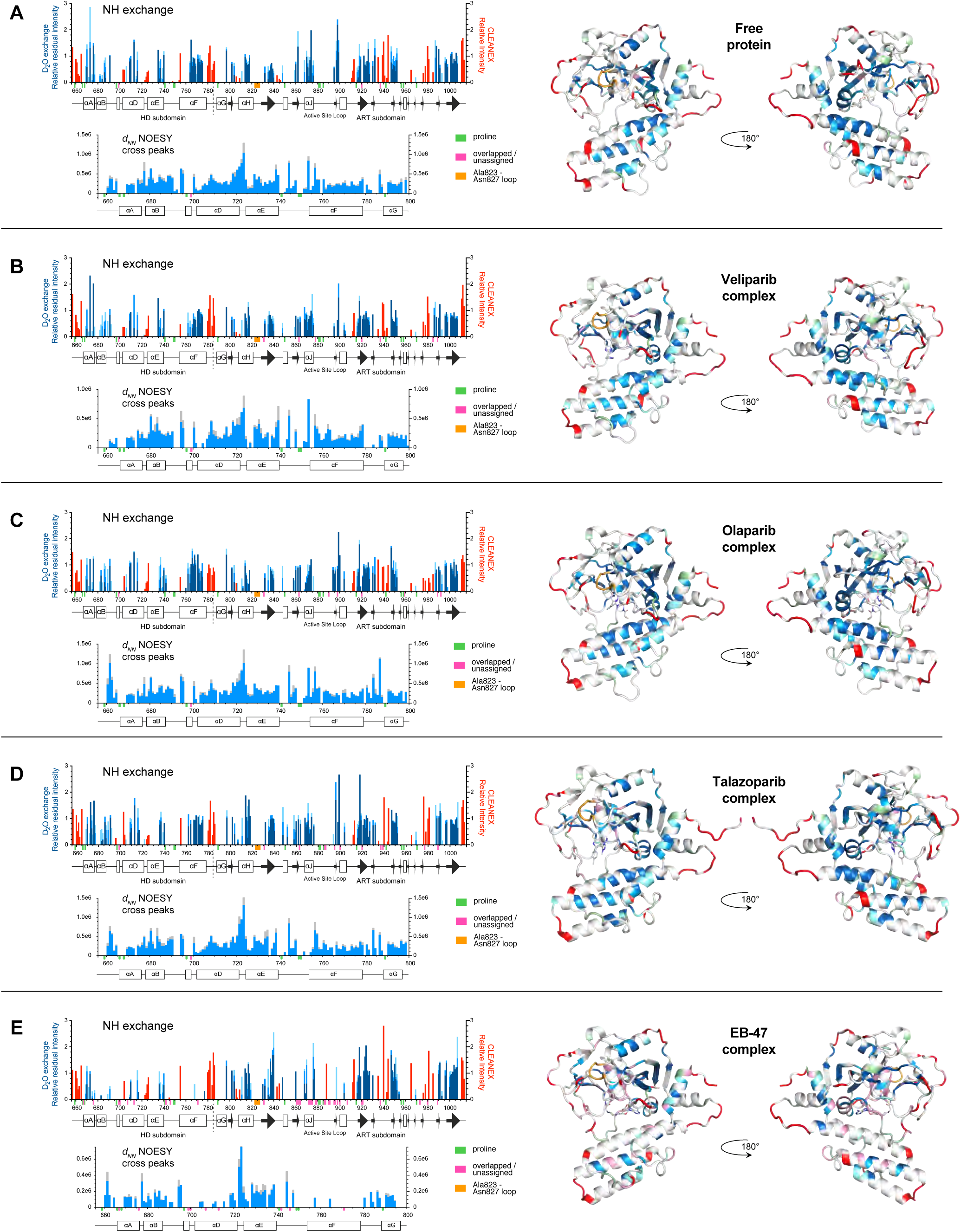
Backbone amide solvent NH exchange data and sequential [NH(*i*), NH(*i+1*)] NOESY cross-peak data (denoted *d*_*NN*_) for the HD subdomain, each shown for A) PARP-1 CAT domain, and its complexes with B) veliparib, C) olaparib, D) talazoparib and E) EB-47. In each case, the data are shown both as histograms and mapped to the relevant CAT domain complex crystal structure. For the NH exchange data, red bars indicate normalised peak intensities measured in CLEANEX-PM experiments, which detect the fastest exchange rates, while the blue bars represent intensities measured in real-time ^2^H_2_O exchange series, which detect slowly exchanging NHs; dark blue represents intensity after 3h, mid-blue after 12h and light blue after 39h, respectively; for details of how these colours were mapped to the structures, see materials and methods section. For the NOESY cross-peak data, runs of continuous (NH, NH) sequential cross peaks indicate stretches of helical conformation (in each case the spectra contain two symmetry-related cross peaks that would ideally have identical intensity; the blue bar represents the lower of these intensities, the grey bar represents the average intensity over both). Small coloured bars beneath the sequence scale and matching colours on the structures are used to indicate the positions of prolines (pale green), overlapped or unassigned signals (pink) and the Ala823-Asn827 loop for which no signals were seen in any spectrum (orange).

**Figure 5:**
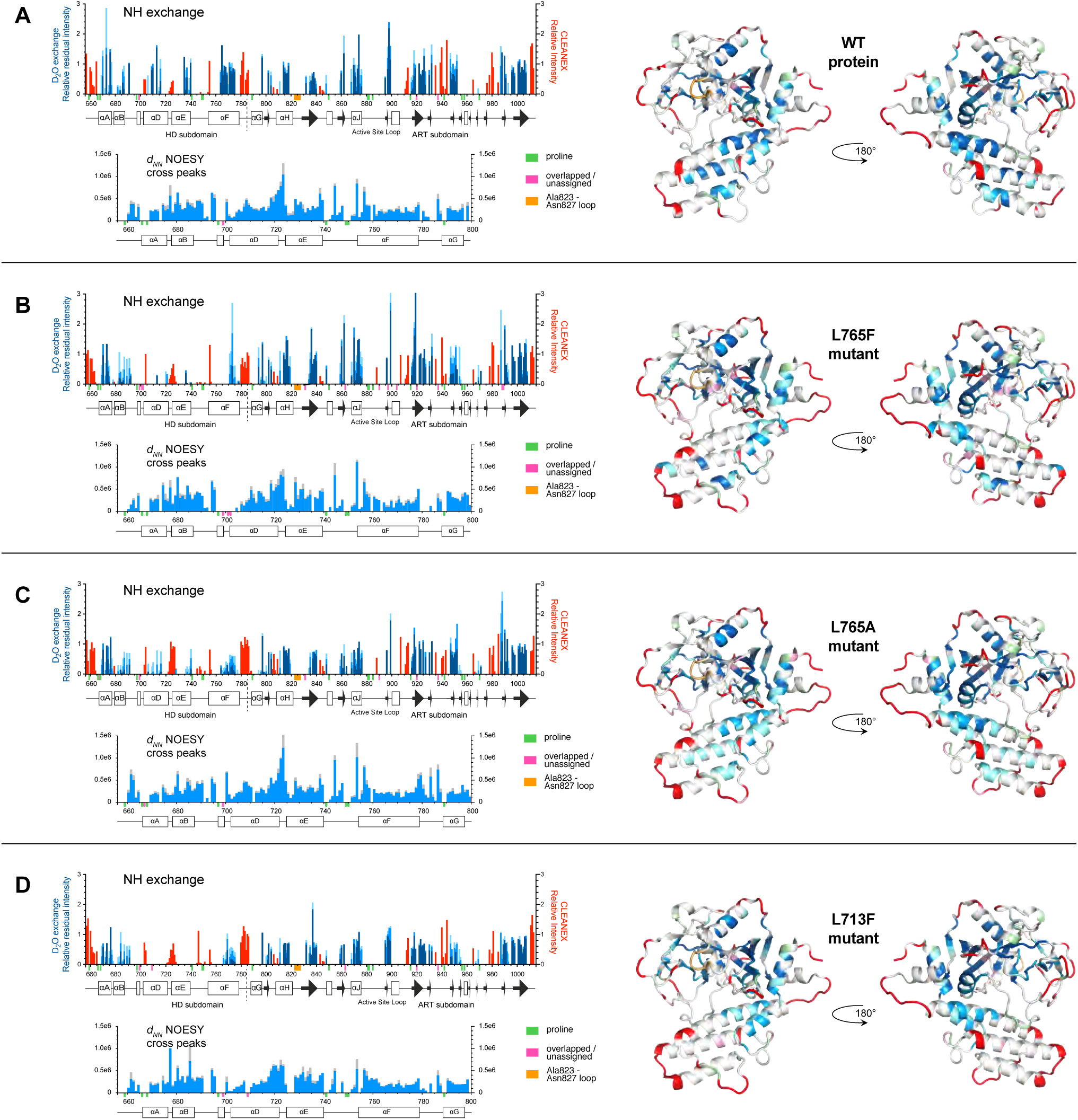
Backbone amide solvent NH exchange data and sequential [NH(*i*), NH(*i+1*)] NOESY cross-peak data (denoted *d*_*NN*_) for the HD subdomain, each shown for for A) PARP-1 CAT domain, B) L765F mutant, C) L765A mutant and D) L713F mutant. All other details as for Figure 4.

The range of possible NH solvent exchange rates in a protein is extremely wide. The fastest involve NH protons in random coil regions that experience no protection through intramolecular hydrogen bonding; model compound studies (56) allow this rate to be estimated for a given set of conditions, and for a protein at pH7 and 25°C it is roughly 10 s^-1^, with variations of up to about a factor of ten in either direction depending on sequence context (see also Figure S16). In contrast, those NHs that are protected by participation in the strongest hydrogen bonds can take many hours, days or longer to exchange with solvent. In order to sample across this wide range, we employed different approaches to detect fast and slowly exchanging NHs.

NMR experiments to detect fast exchanging NH protons are generally based on specific pulse sequences designed to work with samples at equilibrium and are therefore necessarily limited to detecting amide NHs that exchange at a rate comparable to or faster than that of proton longitudinal relaxation. We used the CLEANEX-PM experiment, which largely suppresses unwanted, competing NMR effects (NOE, ROE or TOCSY transfer) that can cause complicating artefacts (57). Given that NHs detected in this experiment must exchange in a fairly narrow rate regime roughly bounded by the random coil exchange rate, here approx. 10 s^-1^ (see above), and the ^1^H longitudinal relaxation rate *R*_*1*_ = 1/ *T*_*1*_ (for the great majority of amide NHs in PARP-1 CAT domain, *R*_*1*_ ≈ 0.4 - 0.7 s^-1^; data not shown), we did not attempt to extract quantitative rate information, but used a single delay time of 150 ms.

The intensities seen in these experiments for the WT protein, mutants and inhibitor complexes are shown by the red bars in Figures 4 and 5. Overall, as expected, there is quite a close correlation between the residues with fast-exchanging NH protons as detected by CLEANEX-PM and those with fast internal motions as revealed by the ^15^N relaxation data. However, in a few regions the CLEANEX-PM data shows fast exchange where the steady-state ^15^N{^1^H} heteronuclear NOE data either shows no reduction or is ambiguous. In the HD subdomain, such cases include the extreme N-terminus of helix D (Lys703 - Arg704; this might in part reflect enhanced base catalysis by the sidechains), while in the ART subdomain they include the linker N-terminal to helix H (Asp805 - Glu809), the N-terminus (Gly843 - Gln846) of helix I, a short surface loop (Ser911 - Gly913) between helix K and the following β-strand, and Asp965 located in a surface loop between two short β-strands; in addition, the surface loop Ser976 - Leu985 is more clearly implicated in the CLEANEX-PM data than in the ^15^N relaxation results. As was also seen for regions with fast motions detected by ^15^N relaxation, several of the regions with fast-exchanging NHs are located at or near contacts that would be present in the recently described complex of PARP CAT domain with HPF1 (24). Very similar patterns are also seen in each of the mutants and inhibitor complexes, again implying that there are no large-scale unfolding events causing substantial portions of the structure to become highly mobile in any of these cases. It is very likely that there may be a number of more localised differences, particularly upon inhibitor binding, but these may be too subtle to detect reliably with these approaches, and in general interpretation at the single residue level is probably not warranted.

However, it is noticeable that at the hinge between helices A and B, residue Val679 shows some degree of enhanced NH exchange in both mutants of Leu765, possibly due to enhanced motion of its neighbour Asp678 (as detected in the ^15^N relaxation data), and also that fast NH exchange in residues Gly843 – Gln846 is at least somewhat reduced in the veliparib, olaparib and talazoparib complexes relative to the WT and mutant proteins, and the EB-47 complex.

To characterise more slowly exchanging NH protons, we recorded a series of TROSY spectra for each of the WT, mutant and inhibitor complex samples, to follow the decay of their NMR signals in real time after re-suspending lyophilised protein in ^2^H_2_O (to which inhibitor was added immediately afterwards in the case of the complexes); each spectrum required approximately three hours to acquire, this timescale being set by sensitivity considerations and the required spectroscopic resolution. The intensities seen in these experiments are indicated by the blue bars in Figures 4 and 5, with darker blues corresponding to intensities at later time points and hence to slower exchange. The slowest exchanging NHs correspond clearly to elements of secondary structure that are protected from solvent exchange, but it is notable that not all such elements are protected in these experiments. In the HD subdomain, the most prominently protected region in the WT free protein is the C-terminal part of helix F, with moderately strong protection also seen for the central part of helix D (where an approximately periodic pattern of protection appears involving residues Ala709, Tyr710, Leu713, Glu715 and Val716, packed against helices E and F) and helix A (where again a periodic pattern appears, involving Gln670, Ile673 and Ile676, all packed against helices G and J); somewhat less pronounced protection is seen for parts of helix B and the C-terminal part of helix E. In the ART subdomain, helix J, which packs across helix F near its mid-point, is strongly protected, while the most prominent protection is seen for the central three β-stands (Asp829 - Glu840, Ile916 - Ala925, Val997 - Phe1007) of the large 5-stranded sheet in the core of the domain. Slightly less pronounced protection is seen for the inner three β-stands (Ile895 - Ala989, Ser947 - Gly950, Glu988 - Val991) of the four-stranded sheet that partly lines the active site cavity, and also for the central part of helix H (Ile814 - Val818).

The patterns of slowly exchanging NHs observed for the free protein are preserved largely unchanged in the complexes with veliparib, olaparib and talazoparib, but in the case of the EB-47 complex there are more substantial changes. Most notably, there is almost complete loss of protection for both helices B and F, at least on the timescale detected in this experiment, as well as a partial loss for helix D, where residue Leu713 seems to be affected specifically (Figure 5). For the mutants, protection is lost to varying degrees for helices D, E and F; L713F shows the most pronounced losses closely followed by L765F, while those for L765A are less pronounced (Figure 5). Notably, it is mainly the C-terminal portion of helix E and the middle portion of helix F that are affected, presumably reflecting proximity to the mutation sites.

To further test the structural significance of these observations, particularly concerning the helices in the HD subdomain, we measured ^15^N HSQC-NOESY spectra (Figures 4 and 5). This revealed that the patterns of NH-NH NOE cross peaks between sequentially neighbouring residues, characteristic of persistently formed helices, are essentially maintained in all of the helices in all of the complexes and mutants, with the notable exceptions of helices F and D in the EB-47 complex. In those latter two cases, the number of identifiable cross peaks is very substantially reduced. Considerable caution is needed in interpreting this result, however, since these NOE cross-peaks might become undetectable for several reasons. In both helices, several of the NH signals involved are substantially weaker in the TROSY data from the EB-47 complex than in the corresponding WT spectrum, to the point where NOESY cross peaks involving those signals in the EB-47 complex would be expected to be well below the detection threshold in our experiments; at the same time others have shifted to positions where there is more signal overlap. Thus, it remains possible that these helices do still persist at least to some degree, and consistent with this the pattern of chemical shift changes seen for the NH protons of both helices do not show any clear trend towards random coil values, nor do they differ qualitatively from the patterns of shift changes seen for the other inhibitor complexes, particularly those for talazoparib (Figure S17). On the other hand, the intensity losses for many of the NHs in these helices is significantly greater than for other regions of the protein, or for HD subdomain helices in the other complexes or mutants, strongly suggesting that there is significant additional signal broadening in these cases. This in turn would be consistent with helices D and F in the EB-47 complex being substantially perturbed, even if not completely losing their coherent structure. Loss of protection in the case of helices such as B in the EB-47 complex, where the pattern of NOESY cross peaks remains largely intact, presumably results from increased breathing motions that increase the exposure of the NHs to solvent, but without substantial perturbation of the overall helical conformation. Consistent with this, the WT CAT domain, mutants and inhibitor complexes all show essentially identical CD spectra (Figure S18).

In summary, the exchange data show that the NH protons located in specific regions of helices in the HD subdomain show considerable differences in the extent to which they are protected against solvent exchange. These differences are significantly modulated both by complexation with the inhibitors, particularly EB-47, and by the point mutations studied here, presumably reflecting changes in the extent of underlying, slow conformational transitions that transiently disrupt helical hydrogen bonds.

### X-ray crystal structures

To gain further insights into the nature of differences in the HD subdomain between the different inhibitor complexes, we analysed X-ray crystal structures of the complexes of PARP-1 CAT domain with veliparib, olaparib, talazoparib and EB-47 (Table 1). In the cases of olaparib and EB-47, no crystal structures of complexes with the CAT domain of human PARP-1 had been published at the time this study was undertaken, so new structures were determined as part of this work (PDB 7AAD and 7AAB respectively); in addition, since the previously existing structure of the veliparib complex (PDB 2RD6) was in a markedly different crystal form from these new structures and that of the apo-protein, a new structure was also determined for the veliparib complex (PDB 7AAC). In the event, another structure of the EB-47 complex (PDB 6VKQ) (58) was determined in parallel while this work was in progress, while in the case of talazoparib two structures (PDB 4UND and 4PJT) have been published previously (59,60), and it is of interest also to include all these in the analysis. In all of these comparisons, the new structure of the human PARP-1 CAT domain apo-protein (PDB 7AAA) was used as reference.

Figures 6A-D show a superposition of all of these structures onto the ART subdomain of the apo-protein (showing only chain A for structures having multiple chains), from which two points are quickly clear: i) the backbone conformations of the ART subdomains in all the complexes are highly similar, and ii) the spatial relationship between the ART and HD subdomains varies substantially amongst the complexes. Rmsd values for these superpositions appear in Table 2; as before, superposition statistics are included in which the rmsd is calculated over a different set of atoms to that used in the fit so as to emphasise differences in the relationships between HD and ART (see also Tables S1-S5 for tables of superposition statistics including all chains as well as for individual helices and combinations of helices, and Figure S19 for views of different superpositions). Some of these variations must reflect the different crystal contacts and packing present in the various different crystal forms, but despite this complication it is very clear that the structural perturbations to the HD subdomain caused by inhibitor binding depend strongly on the extent to which each inhibitor molecule extends towards helix F (Figure 6E). Neither veliparib nor olaparib have steric clashes with residues of helix F, and in those cases the variations in the HD are no larger than those seen within the ART subdomain; indeed, if the comparison is restricted to 7AAA, 7AAC and 7AAD, all of which have the same crystal form, then the structural differences are very small indeed in both HD and ART, and the spatial relationships between ART and HD are essentially identical. The larger differences between 7AAA and the previous veliparib complex structure (2RD6) most likely reflect the very different crystal form in the latter case. In the case of olaparib, although the molecule does approach helix F quite closely, at its nearest point (the cyclopropyl ring) it interdigitates between protein sidechains (Asp766 and Leu769), and again the structure and disposition of the HD subdomain remain almost completely unperturbed relative to 7AAA.

**Figure 6:**
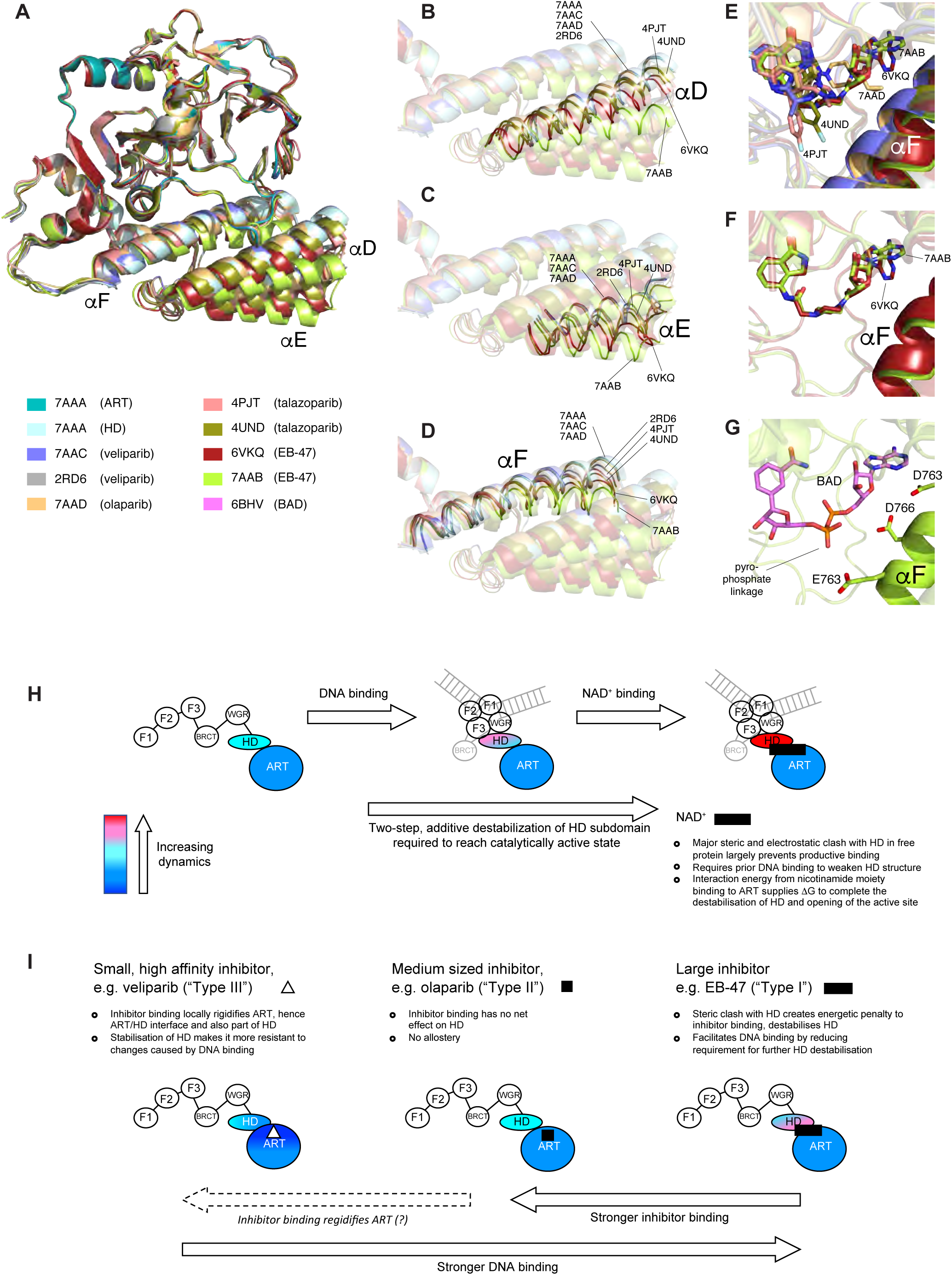
A-D) Superposition of PARP-1 CAT domain in the apo-protein and in inhibitor complexes with veliparib (PDB 7AAC and 2RD6), olaparib (PDB 7AAD), talazoparib (PDB 4PJT and 4UND) and EB-47 (PDB 7AAB and PDBB:6VKQ), in each case superposing using only the ART subdomain (N, Cα, C’ of residues 790-936, 939-1009) of 7AAA. The inhibitor structures themselves are omitted from these views. E) Close up of the same superposition as in A-D), showing the inhibitors bound in the nicotinamide binding pocket; those parts of inhibitors that approach closely to helix F are labelled. The apo-protein is omitted from this view. F) Close up of the same superposition as in E), showing only the two EB-47 complexes (PDB 7AAB and PDB 6VKQ), demonstrating the different binding poses of the adenosine moiety in the two cases. G) Hypothetical binding pose of the non-hydrolysable NAD+ analogue benzamide adenine dinucleotide (BAD) in the presence of the HD subdomain of PARP-1. The model was obtained by superposing the ART domains (N, Cα, C’ of residues 790-936, 939-1009) of the BAD complex with PARP-1 ΔHD-CAT (PDB 6BHV) and the EB-47 complex (PDB 7AAB) with that of the apo-protein (PDB 7AAA), then displaying the structure of BAD (as sticks) with the backbone of the EB-47 complex. Note the prediction that this arrangement would lead to a severe electrostatic clash between the pyrophosphate linker of BAD and the acidic sidechains on helix F. H-I) Schematic model for contributions of dynamics to allostery in H) PARP-1 activation and I) PARP-1 inhibitor binding. H) Shows the proposed two-stage nature of activation by successive interaction with DNA damage and substrate, while I) shows the different contributions to allostery for inhibitors of types I, II and II as defined in Zandrashvili et al. (58).

In contrast, in both the talazoparib and EB-47 complexes the inhibitor clashes with residues of helix F (Glu763 and Gln759 in the case of talazoparib and Asp766 and Asp770 in the case of EB-47), with the result that the whole HD subdomain is displaced relative to ART to varying degrees. For both talazoparib and EB-47, two different structures are available each having different crystal forms, and it is clear that the extent of displacement of the HD varies substantially amongst all four of these structures due to differences in the identity of the inhibitor, its binding mode and the crystal packing. By far the greatest overall displacement of HD relative to ART occurs in one of the EB-47 complex crystal structures, 7AAB (Figure 6 and Table 2), while related, though smaller, displacements occur in the other EB-47 complex (6VKQ) and the two talazoparib complexes. In each case, there is a small hinge motion at Asp678 (the residue linking helices A and B shown to be flexible by ^15^N relaxation), so that helix A remains “attached” to the ART subdomain while helix B moves with the remainder of the HD; at the C-terminal end of the HD subdomain, linkage to ART is via a longer flexible region (Gly780 - Lys787). The relative movement of the HD varies both in extent and direction amongst the structures, but the mutual arrangement of the helices (other than A) within the HD is largely unperturbed in each case. However, one clear difference amongst the structures concerns helix F, which is distorted to differing degrees in each (Figure 6D). The strongest perturbation occurs in the EB-47 complex 6VKQ, where a sharp kink is seen at residue Asp766, near the site of the steric clash with the inhibitor; similar but less pronounced distortions are present in helix F in the other EB-47 complex structure 7AAB and the two talazoparib complex structures.

In addition to the different displacements of the HD, there is also a striking difference between the two structures of the EB-47 complex in the binding mode of the adenosine moiety (Figure 6F); the inhibitors superpose closely throughout the benzamide moiety but differences accumulate through the piperazine and ribose rings, and the *χ* angle of the adenosine-ribose connection differs by over 30° between the structures (−68.1° in 7AAB, -32.4° in 6VKQ, averaged across chains in each case). This difference seems likely to reflect a much weaker contribution to the binding from the adenosine moiety, which has no particularly specific interactions with nearby protein groups in either binding pose. However, comparison with the complex of PARP-1 ΔHD CAT domain with the non-hydrolysable NAD+ analogue BAD (19) clearly shows a close similarity with the pose of the adenosine moiety of EB-47 as seen in 7AAB, but not 6VKQ (Figures 6F,G). For the other two cases where multiple structures are available, veliparib and talazoparib, the inhibitor structures do not include flexible portions and the binding poses observed are more similar when compared between crystal forms.

Although the crystal structures are intrinsically static, comparisons across the structures reveal regions where variations amongst their backbone conformations are concentrated, which correlate well with flexible regions revealed by the ^15^N relaxation and/or NH exchange experiments. Such regions can be seen in Figure 6, while for those structures having multiple copies in the asymmetric unit similar patterns appear in the deviations between chains from the same structure (Figure S20). As commented earlier, several of these flexible regions correspond to sites known to be involved in inter-domain or intermolecular interactions either in the full-length DNA-bound context (Phe744 - Pro749 contacts the WGR domain in PDB 4DQY) or in the complex with the accessory protein HPF1 (Ala823 - Asn827, His934 - Leu941 and Ser977 - Thr981 contact HPF1 in PDB 6TX3). However, particularly in the HD subdomain, some flexible loops do not correspond to known interaction sites, including most notably the loop connecting helices D and E (Gln722 - Ser725). In this loop, which also experiences the greatest overall movements as a result of inhibitor binding, the variations amongst the structures are particularly strong, and even extend into substantial distortions of the nearest turns of the helices themselves, suggesting this region may be a particular “hotspot” for local flexibility (Figure S19E).

Further evidence of flexibility and plasticity in the HD subdomain comes from comparison of the two EB-47 complex structures. The very different displacements of the HD in these structures demonstrates that crystal packing forces must differ substantially in the two cases. This in turn must presumably underlie another big difference between them, which is that for 6VKQ, but not 7AAB, the quality of the observed electron density for the HD was particularly weak, which led to considerable difficulties in modelling this portion of the structure (58). This is also reflected in the much higher B factors for the HD subdomain than those for the ART subdomain for 6VKQ (Figure S21); similar effects were seen in the closely related structure of a complex of PARP-1 CAT domain with the inhibitor UKTT15 (6VKO) that was also reported recently (58), whereas for 7AAB (and for the other structures discussed here) the quality of the electron density was essentially normal for both the HD and ART subdomains. The origin of these differences must lie in the detailed nature of the crystal contacts that presumably prevent the HD in 6VKQ from being displaced to the same extent as it is in 7AAB, but the fact that packing forces are capable of destabilising the HD subdomain so strongly in 6VKQ suggests that the steric clash with EB-47 affects the HD to the point where it may even come close to unfolding.

In summary, the comparison amongst these various crystal structures shows that inhibitor structures that reach beyond the nicotinamide binding pocket can cause large relative displacements of the HD to the extent that they clash against the HD subdomain, and in the case of the largest inhibitor studied here, EB-47, the stability of the folded structure of the HD may become quite marginal as a result. Regions of flexibility identified by the NMR experiments correspond to regions of variability between the different crystal structures, and the region around the loop linking helices D and E appears to be particularly mobile.

## DISCUSSION

Catalytic activation of PARP-1 in response to detection of DNA damage is central to its role in co-ordinating DNA repair, in particular of DNA single-strand breaks that are the most abundant form of DNA damage. The architecture of the complex formed by PARP-1 upon binding to DNA damage sites (14,15) immediately makes clear the importance of the HD subdomain in transmitting the activation signal from the DNA-binding domains to the catalytic centre, since it forms the only structural connection between them. It has previously been proposed that changes in dynamics within the HD play a key role during this process (16), but a detailed description has been elusive. In this study we have used a combination of approaches to probe the nature of dynamic processes involving the HD over a wide range of rates, and to investigate how these are modulated, both by mutations that partially mimic the activating effects of DNA binding and also by binding of a selection of PARP inhibitors that in different ways partially mimic the effects of substrate (NAD^+^) binding.

Our data show that it is principally the slower conformational exchange processes, those detected by the NH solvent exchange experiments, that show differences amongst the WT protein, mutants and inhibitor complexes, whereas for the fastest processes, those detected mainly by the ^15^N relaxation experiments, there is little variation across the different species. The most prominent effects are seen as changes in the stability of particular parts of individual helices, and these parallel differences seen amongst the available crystal structures, including movements of the whole HD subdomain relative to the ART as well as varying degrees of distortion and relative shifts of individual helices. While these perturbations resulting from mutation or inhibitor binding are in some cases substantial, they are not sufficient to destroy the structural integrity of the HD subdomain; in almost all cases where individual helices loose protection against NH solvent exchange, the presence of NOESY cross peaks between sequentially adjacent amide protons shows the helices remain substantially intact in solution, implying that increases in NH exchange on inhibitor binding or mutation are caused by increased “breathing” motions of the helices whereby individual hydrogen bonds are transiently broken while the majority remain intact. That said, there are indications that the HD may be only marginally stable in its folded form at least in the case of the complex with EB-47, the inhibitor of those studied here that most closely resembles substrate. In one of its crystal forms (6VKQ), the HD subdomain (but not the ART) shows very poor electron density and consistently high B factors throughout (58), while in solution, where the absence of crystal packing forces may increase the tendency to disorder, it is unclear to what extent the helix most strongly affected by presence of the inhibitor, helix F, remains truly intact. Overall, however, it seems unlikely the changes seen in any of these cases would be sufficient in themselves to correspond to a completely activated state of PARP-1, in which the active site must accommodate during the reaction cycle not only a single substrate molecule but also the elongating, and occasionally branching, PAR chain.

We propose, therefore, that activation of PARP-1 is in all probability a two-step process, in which first DNA binding and then substrate binding contribute consecutively and additively to successive destabilisation of the folded structure of the HD subdomain, as summarised in Figure 6H. Previous work has shown that DNA binding also causes changes in the HD subdomain, with the strongest effects on NH solvent exchange seen for helices B, D and F (Figure 1D) (16), while similar effects for EB-47 binding as reported here were found in a parallel HXMS study of allostery involving PARP inhibitors and their influence on the HD subdomain (58). Comparison of the crystal structures of the inhibitor complexes with that of the DNA-bound complex (4DQY) also shows differences, most notably that DNA binding causes helix F to become straight (Figure 1J,K), whereas inhibitor binding, particularly of EB-47, has the opposite effect of increasing the bend in helix F, with movements in other helices, particularly E, that also differ between DNA and EB-47 binding (Figures 6A-D). Subjecting the HD to opposing forces in this way, as would happen during successive binding of DNA and substrate, might thus be expected to destabilise the structure, since it makes it harder for a single conformation to accommodate them simultaneously.

It is notable from the results of the HDMX study that addition of EB-47 to DNA-bound full-length PARP causes broadly similar changes in NH solvent exchange within HD to those reported here, but apparently does not result in substantially more extensive local unfolding. However, EB-47 is only a partial mimic of the true substrate NAD^+^, and one of the most prominent differences between them is that in NAD^+^ the nicotinamide and adenosine moieties are linked through, *inter alia*, a pyrophosphate group, whereas in EB-47 the equivalent portion of the linker is a piperazine ring. Comparison with the structure of the BAD ΔHD-CAT complex shows that, were the HD subdomain present, binding of NAD^+^ in a similar pose to that of BAD would bring about a highly unfavourable electrostatic interaction between the pyrophosphate group and a strongly negative patch on helix F formed by acidic sidechains on three adjacent helical turns (Glu763, Asp766 and Asp770; see Figure 6G) (19). It seems likely this could in large part be the reason for the much weaker binding interaction of BAD (and presumably also NAD+) with PARP species that include the HD than that seen with EB-47; it could also be that this interaction is required to destabilise the HD to the point where it reaches the fully activated state. In the case of the activating mutants, the differences in patterns of NH protection from solvent exchange again indicate that particular parts of individual helices are destabilised, and not surprisingly these effects occur mainly in locations close to the mutations. While it is entirely plausible that disruption of local packing interactions, as must presumably occur in these mutants, would destabilise the HD and thus partially mimic the functional outcome of activation, it appears that the local details of this disruption probably differ from those caused by either DNA or inhibitor binding.

No structural model is currently available for the fully activated state of PARP-1, but on the basis of the data presented here it seems reasonable to propose it would involve significantly greater loss of folded structure in the HD than that seen in any of the species for which structural data exists so far. Given the particularly strong indications of flexibility on a fast timescale for the loop between helices D and E, and the fact that this loop apparently has no interaction partners, it could be that this loop, which is located close to the path that a substrate molecule would need to follow on entering the active site, has a role in nucleating a partial local unfolding process during activation.

Given the existence of extensive parallel studies of NH solvent exchange in this system, it is worth briefly comparing a few key properties of the different techniques used, NMR and HXMS. One clear difference is that NMR has the potential to record data resolved at the single residue level, something that, while possible (61), is harder to achieve in HXMS studies of large systems. Not all residues may be observed in the NMR case, however, which among other considerations is due to a second key difference, sensitivity to exchange rate. Unlike HXMS, where NH exchange is effectively frozen by a jump to extreme low pH prior to tryptic digestion and mass-spectrometric analysis, NMR analysis is typically conducted on a sample in which exchange is still “live” during data acquisition. In principle it might be possible to employ a pH jump approach to freeze the exchange during NMR experiments, but this would be severely complicated in practice, particularly if the protein failed to retain its native fold following the jump as this would greatly impact spectral resolution of the signals; to our knowledge results from such an approach have not been reported. NMR can access faster rates than HXMS by using equilibrium pulse-sequence-based methods such as CLEANEX, or other experiments designed for still faster rates (62), but there is a range of rates, very roughly 0.5-500 s^-1^, that are too slow for such equilibrium methods to be applicable (*k*_*ex*_ < ^1^H *T*_*1*_) yet too fast for real-time exchange measurements to be practical. The fastest rate accessible by real-time NMR measurements depends both on the time required prior to measuring the first point, which can be reduced using rapid mixing pulse-labelling (63), and the time required for measurement of each point, which is set by the intrinsic sensitivity of the system (i.e. available sample concentration, the NMR experiment deployed and spectrometer sensitivity). Unfortunately, a proportion of NH signals in the present study fall into this “blind window” of inaccessible exchange rates, which to some degree limits interpretation.

A related question that affects both techniques equally concerns the relationship between the rate of NH solvent exchange and that of the underlying conformational exchange processes responsible for modulating the exchange. For the most flexible protein backbone regions, those revealed by ^15^N relaxation experiments, the rates of conformational exchange must be faster than overall molecular tumbling, *k*_*ex*_ > (*τ*_*c*_)^-1^, meaning they occur on the nanosecond timescale (*τ*_*c*_ ≈ 25 ns for the PARP-1 CAT domain). NH protons in such regions experience essentially no protection from solvent exchange, so they exchange at or near the random coil rate, estimated at 10 s^-1^ under the conditions used here. Thus, in these cases there is roughly a 10^7^-fold difference between the rates of conformational exchange and NH proton solvent exchange. In contrast, in the limit of extreme protection in the most stable structural regions, disruption of hydrogen bonds due to conformational exchange can become rate limiting (the so-called EX1 limit) and then rates of conformational and NH exchange for a given NH become equal. This condition is rarely reached in practice. Between these two extremes intermediate behaviour is expected (see Figure S16), but generally little quantitative detail is available. However, in the particular case of the protein re-modelling required to accommodate binding of EB-47 to PARP-1 CAT domain, further information is available from the values of *k*_*on*_ and *k*_*off*_ as measured by SPR. The fact that the on-rate for EB-47 binding is very similar to that of the other inhibitors tested (except veliparib) whereas the off-rate is 3 orders of magnitude faster than for the others indicates that the additional re-modelling of the HD subdomain required in the EB-47 case occurs as a *consequence* of EB-47 binding, not prior to binding; in other words this is apparently a case of “induced fit”, rather than “conformational selection”. This in turn means that the reversal of the re-modelling process must occur *during* dissociation rather than afterwards, acting to accelerate expulsion of the ligand. The conformational fluctuations responsible for breaking individual hydrogen bonds in helix F and allowing NH solvent exchange to occur while EB-47 is bound are presumably closely related to those that occur during this overall process of protein re-modelling coupled with dissociation, and likely occur at a similar rate characterised by *k*_*off*_ = 0.16 s^-1^, or faster. Overall, therefore, it seems safe to conclude that these conformational fluctuations are significantly faster than NH solvent exchange in this case.

The SPR data is also relevant to another outstanding question, namely how it is that binding of veliparib causes mild *stabilisation* of the HD subdomain, and thus leads to allosteric effects in the opposite sense to those caused by EB-47 (prior inhibitor binding causes weaker binding of PARP-1 to DNA damage sites in the case of veliparib, but stronger binding in the case of EB-47) (58). The most noticeable characteristic of veliparib relative to the other inhibitors studied here is its small size; when bound, no part of the veliparib molecule comes close to any part of the HD. Despite this, veliparib has a high affinity, not much below those of talazoparib and olaparib, implying that the more limited number of interactions it makes with protein are individually strong. This in turn suggests it is likely to cause stabilisation of nearby protein structure by quenching local flexibility, and this effect may transmit through intervening parts of the protein structure to reach helix B in the HD, thereby slowing its NH exchange rates as was seen in HXMS experiments with the veliparib complex (58). Such an explanation raises the question of why little or no evidence was seen in either the HXMS or NMR studies for additional protection in the intervening structural regions of the ART (such as residues Arg865 – Thr867), however this could easily be because NH protons of these residues have exchange rates too fast to detect using these experiments, even if they are slowed to an extent by additional protection gained upon inhibitor binding. Indeed, even the slowing of the exchange in helix B involved rates too fast to have been detected in our real-time NMR experiments.

A consequence of the model proposed here, where substrate participates in the final part of the activation process, is that some key structural requirements of substrate and inhibitors work in opposition; the interactions of substrate with the HD that help unlock the active site have an energetic cost that reduces binding affinity to a very low level. This clearly aids in achieving the very tight regulation of PARP activity that is crucial to prevent indiscriminate and inappropriate PAR-ylation, by blocking substrate access to the active site in un-activated PARPs. For inhibitors, however, the more they mimic those aspects of substrate structure involved in interactions with the HD, the more weakly they will bind to free PARP, making them less likely to emerge from conventional drug screening strategies. Their effectiveness in the clinic depends on many factors, but important amongst these is the balance between inhibitor binding and trapping efficiency, which in turn depends critically on the degree of interaction between the inhibitor and the HD (58).

While much recent work, including that described here, has cast light on the processes involved in activation and their effects on the HD subdomain, it remains true that a detailed description of the fully activated state of PARP-1 has yet to emerge. It is likely that in this state at least some parts of the HD subdomain may become substantially disordered, and that the detailed nature of this disorder may be key to further understanding its function. The study of intrinsically disordered proteins (IDPs) is of course a topic of intensive and active research worldwide, but in the case of the PARP HD subdomain these disorder-related properties remain largely hidden until the molecule is actively engaged in the catalytic cycle. As such, one could consider the HD subdomain to be a “cryptic” IDP; the study of such systems poses a particular challenge to structural biology for the future.

## DATA AVAILABILITY

Crystal structure co-ordinates and structure factors have been deposited with the PDB database under the following accession codes: WT free human PARP-1 CAT domain, 7AAA; PARP-1 CAT domain complex with veliparib, 7AAC; PARP-1 CAT domain complex with olaparib, 7AAD; PARP-1 CAT domain complex with EB-47, 7AAB. NMR chemical shift assignments have been deposited at the BMRB under the following accession codes: WT free human PARP-1 CAT domain (NH, N, CA and CB assignments), 50454; PARP-1 CAT domain complex with veliparib (NH and N assignments), 50455; PARP-1 CAT domain complex with olaparib (NH and N assignments), 50456; PARP-1 CAT domain complex with talazoparib (NH and N assignments), 50457; PARP-1 CAT domain complex with EB-47 (NH and N assignments), 50458; PARP-1 CAT domain L765F mutant (NH and N assignments), 50459; PARP-1 CAT domain L765A mutant (NH and N assignments), 50460; PARP-1 CAT domain L713F mutant, 50461 (NH and N assignments).

## SUPPLEMENTARY DATA

Supplementary Data comprising Figures S1-S21 and Tables S1-S5 are available.

## ACKNOWLEDGEMENT

We thank the Francis Crick Institute for access to the MRC Biomedical NMR Centre, and assistance from G. Kelly (the Francis Crick Institute receives core funding from Cancer Research UK (FC001029), the UK Medical Research Council (FC001029) and the Wellcome Trust (FC001029)). We thank Christopher Johnson and Stephen McLaughlin for assistance with running and analysing the ITC and CD experiments. We thank Peter Neuhaus for the scripts used to calculate the RMSD values in Table 2, and Leo Kiss and Katherine Stott for careful reading of the manuscript.

