## Supplementary Figures S1-S21 and Tables S1-S5 for "Dynamics of the HD regulatory subdomain of PARP-1; substrate access and allostery in PARP activation and inhibition"

### Supplementary Material

**Figure S1.** Overlay of [$^{15}$N, $^{1}$H] TROSY spectra of PARP-1 CAT domain (residues 657-1014) before and after the lyophylisation procedure described in Materials and Methods in the main paper.

**Figure S2.** Correlation times for overall tumbling ($\tau_c$) for PARP-1 CAT domain and its complexes with veliparib, olaparib, talazoparib and EB-47.

**Figure S3.** Correlation times for overall tumbling ($\tau_c$) for WT PARP-1 CAT domain and its point mutants L713F, L765F and L765A.

**Figure S4.** Steady-state {$^{1}$H} $^{15}$N NOE data for PARP-1 CAT domain and its complexes with veliparib, olaparib, talazoparib and EB-47, showing residues of HD subdomain only.

**Figure S5.** Steady-state {$^{1}$H} $^{15}$N NOE data for WT PARP-1 CAT domain and its point mutants L713F, L765F and L765A, showing residues of HD subdomain only.

**Figure S6.** Steady-state {$^{1}$H} $^{15}$N NOE data for PARP-1 CAT domain and its complexes with veliparib, olaparib, talazoparib and EB-47, showing residues of ART subdomain only.

**Figure S7.** Steady-state {$^{1}$H} $^{15}$N NOE data for WT PARP-1 CAT domain and its point mutants L713F, L765F and L765A, showing residues of ART subdomain only.

**Figure S8.** $^{15}$N longitudinal relaxation time ($T_1$) data for PARP-1 CAT domain and its complexes with veliparib, olaparib, talazoparib and EB-47, showing residues of HD subdomain only.

**Figure S9.** $^{15}$N longitudinal relaxation time ($T_1$) data for WT PARP-1 CAT domain and its point mutants L713F, L765F and L765A, showing residues of HD subdomain only.

**Figure S10.** $^{15}$N longitudinal relaxation time ($T_1$) data for PARP-1 CAT domain and its complexes with veliparib, olaparib, talazoparib and EB-47, showing residues of ART subdomain only.

**Figure S11.** $^{15}$N longitudinal relaxation time ($T_1$) data for WT PARP-1 CAT domain and its point mutants L713F, L765F and L765A, showing residues of ART subdomain only.

**Figure S12.** $^{15}$N spin-locked relaxation time ($T_{1\rho}$) data for PARP-1 CAT domain and its complexes with veliparib, olaparib, talazoparib and EB-47, showing residues of HD subdomain only.

**Figure S13.** $^{15}$N spin-locked relaxation time ($T_{1\rho}$) data for WT PARP-1 CAT domain and its point mutants L713F, L765F and L765A, showing residues of HD subdomain only.

**Figure S14.** $^{15}$N spin-locked relaxation time ($T_{1\rho}$) data for PARP-1 CAT domain and its complexes with veliparib, olaparib, talazoparib and EB-47, showing residues of ART subdomain only.

**Figure S15.** $^{15}$N spin-locked relaxation time ($T_{1\rho}$) data for A) WT PARP-1 CAT domain and its point mutants B) L713F, C) L765F and D) L765A, showing residues of ART subdomain only.

**Figure S16.** Relationship between NH solvent exchange rates and conformational fluctuation rates.

**Figure S17.** Schematic [$^{15}$N,$^{1}$H] correlation plots showing chemical shift differences for the amide group $^{1}$H and $^{15}$N signals of residues in helix F.

**Figure S18.** CD spectra of PARP-1 CAT domain, its complexes with veliparib, olaparib, talazoparib and EB-47, and the point mutants L713F, L765F and L765A.

**Figure S19.** Superpositions of PARP-1 CAT domain in the apo-protein and in inhibitor complexes with veliparib, olaparib, talazoparib and EB-47, in each case superposing each structure onto atoms of 7AAA.

**Figure S20.** Co-ordinate deviations between the different protein chains in the asymmetric unit for the structures of free PARP-1 CAT domain and its complexes with talazoparib, veliparib, olaparib, and EB-47, as well as for chains C and F of the complex of PARP-1 F1, F3 and WGR-CAT with a DNA duplex.

**Figure S21.** B factors for each chain in each of the structures of free PARP-1 CAT domain and its complexes with veliparib, olaparib, talazoparib, and EB-47, as well as for chains C and F of the complex of PARP-1 F1, F3 and WGR-CAT with a DNA duplex.

**Table S1.** Superposition statistics for individual helices of PARP-1 CAT domain (all chains fitted onto 7AAA).

**Table S2.** Superposition statistics for groups of helices in PARP-1 HD subdomain, demonstrating differences in helical packing (all chains fitted onto 7AAA).

**Table S3.** Superposition statistics for individual helices in PARP-1 HD subdomain, comparing different chains in the same structure.

**Table S4.** Superposition statistics for the HD and ART subdomains of PARP-1 CAT domain (all chains onto 7AAA).

**Table S5.** Superposition statistics for the HD and ART subdomains of PARP-1 CAT domain between different complexes with the same inhibitor (all chains).

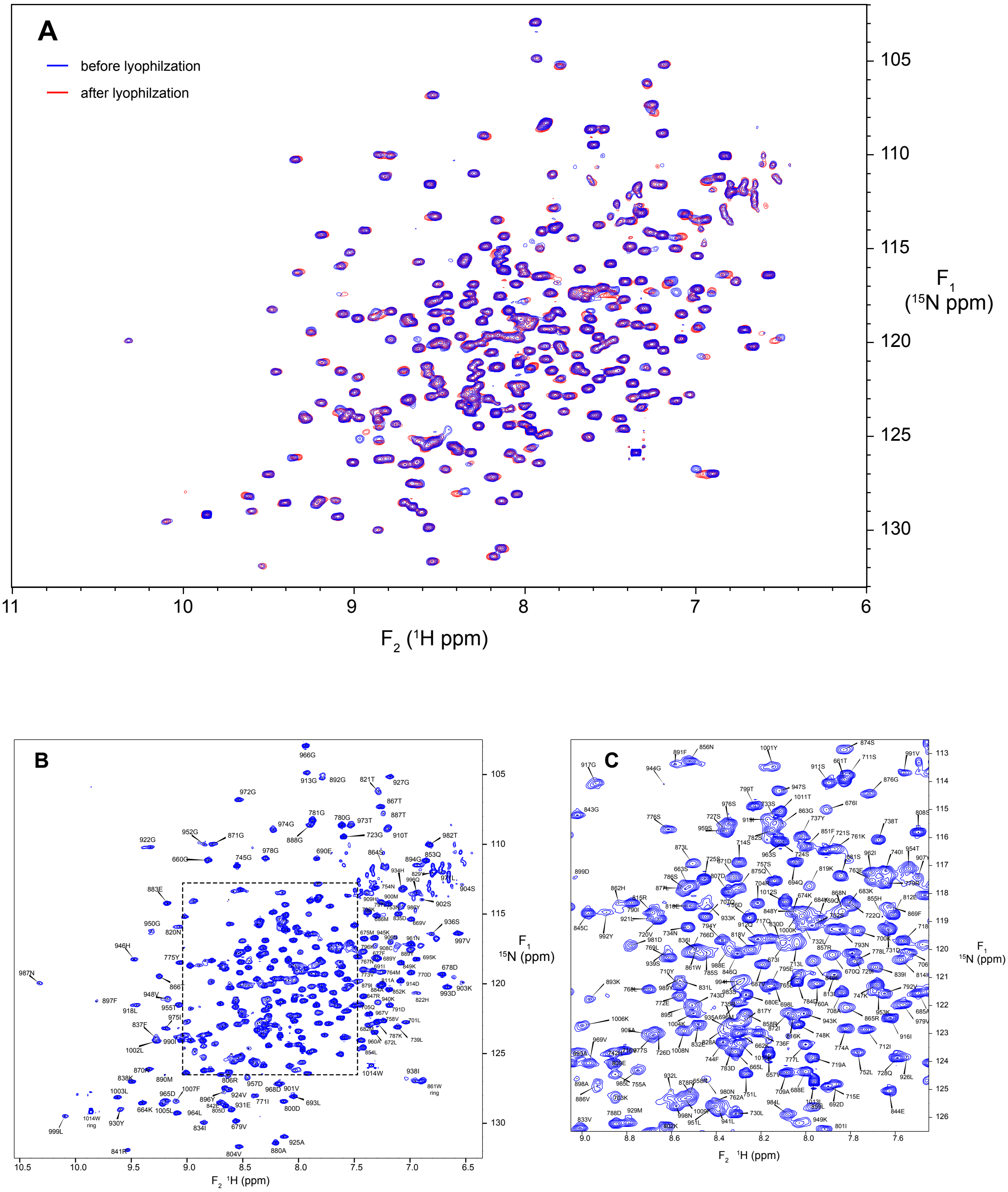


**Figure S1.** A) Overlay of [15N, 1H] TROSY spectra of $^{15}$N-labelled PARP-1 CAT domain (residues 656-1014) before (blue) and after (red) the lyophylisation procedure described in Materials and Methods in the main paper. For this control the protein was resuspended in $H_2O$, but in the real-time $^2H_2O$ exchange experiments described in the main paper, protein was resuspended in $^2H_2O$ and a time series of TROSY spectra recorded. A small number of small differences in chemical shift are likely caused by partial sublimation of buffer components during lyophilsation and did not interfere with interpretation of the data. B) [$^{15}$N, $^1$H]-TROSY spectrum of $^{15}$N-labelled human PARP-1 CAT domain acquired at 800 MHz and 25 °C, showing backbone amide NH signal assignments. Protein concentration was 400 μM in 50 mM [$^2H_{11}$]Tris, pH 7.0, 50 mM NaCl and 2 mM [$^2H_{10}$]DTT in 95:5 $H_2O$/$^2H_2O$. C) Expansion of the most heavily overlapped region of the spectrum shown in B).

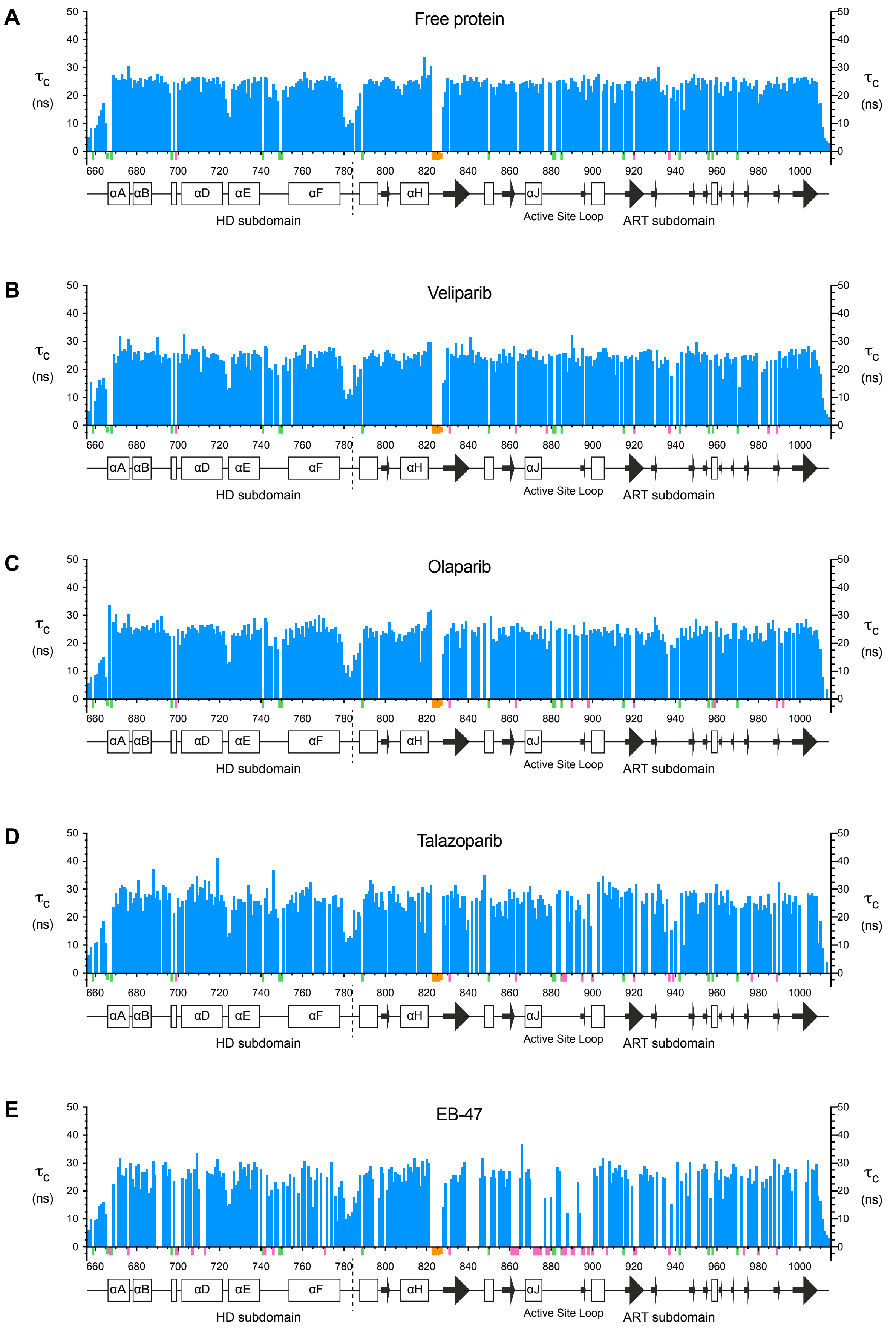


**Figure S2.** Correlation times for overall tumbling ($\tau_c$) for A) PARP-1 CAT domain and its complexes with B) veliparib, C) olaparib, D) talazoparib and E) EB-47. Values were calculated for each residue from $T_1$ and $T_{1\rho}$ data as described in Materials and Methods. Small coloured bars beneath the sequence scale are used to indicate the positions of prolines (pale green), overlapped or unassigned signals (pink) and the Ala823-Asn827 loop for which no signals were seen in any spectrum (orange).

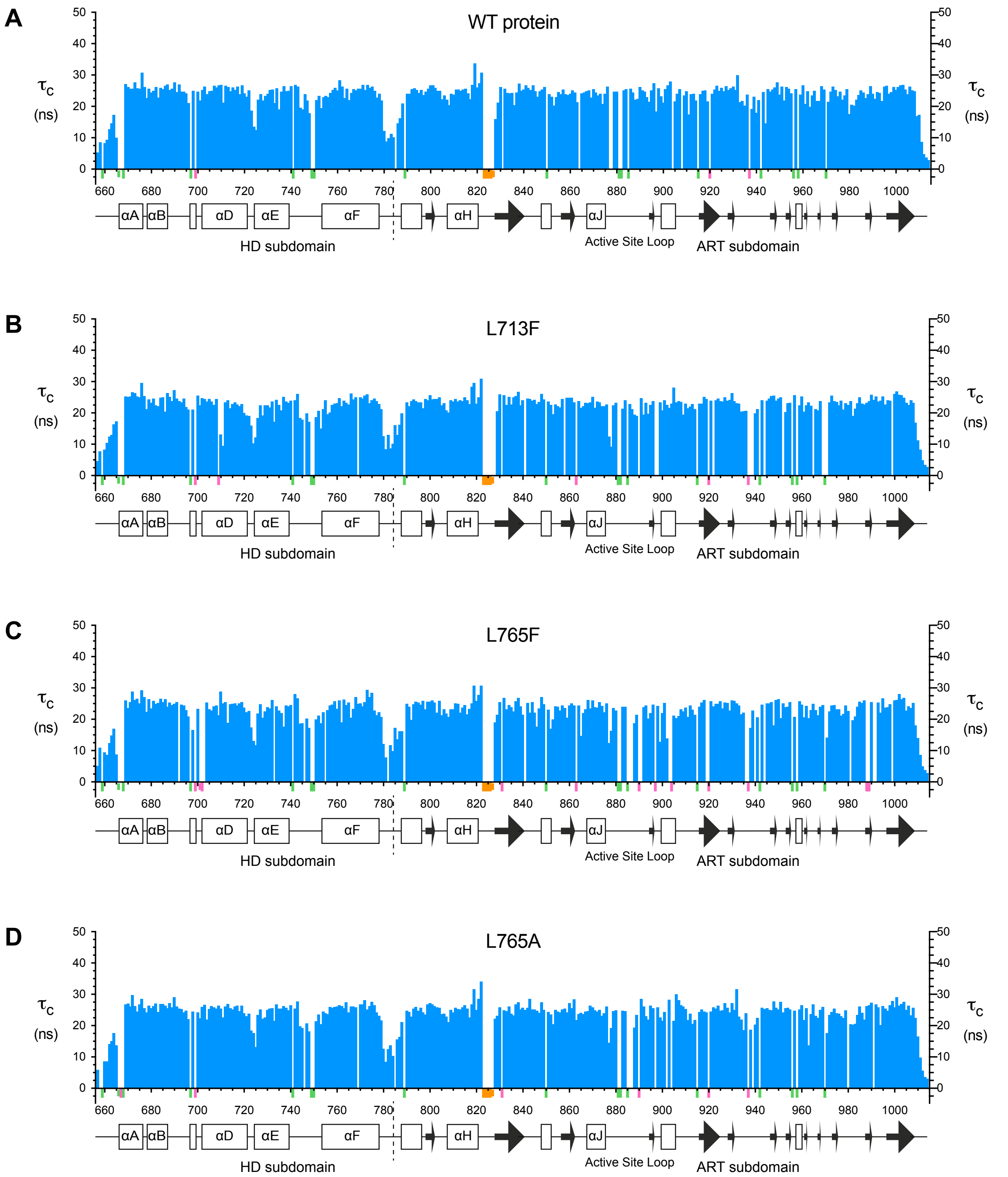


**Figure S3.** Correlation times for overall tumbling ($\tau_c$) for A) WT PARP-1 CAT domain and its point mutants B) L713F, C) L765F and D) L765A. Values were calculated for each residue from $T_1$ and $T_{1\rho}$ data as described in Materials and Methods. Small coloured bars beneath the sequence scale are used to indicate the positions of prolines (pale green), overlapped or unassigned signals (pink) and the Ala823-Asn827 loop for which no signals were seen in any spectrum (orange).

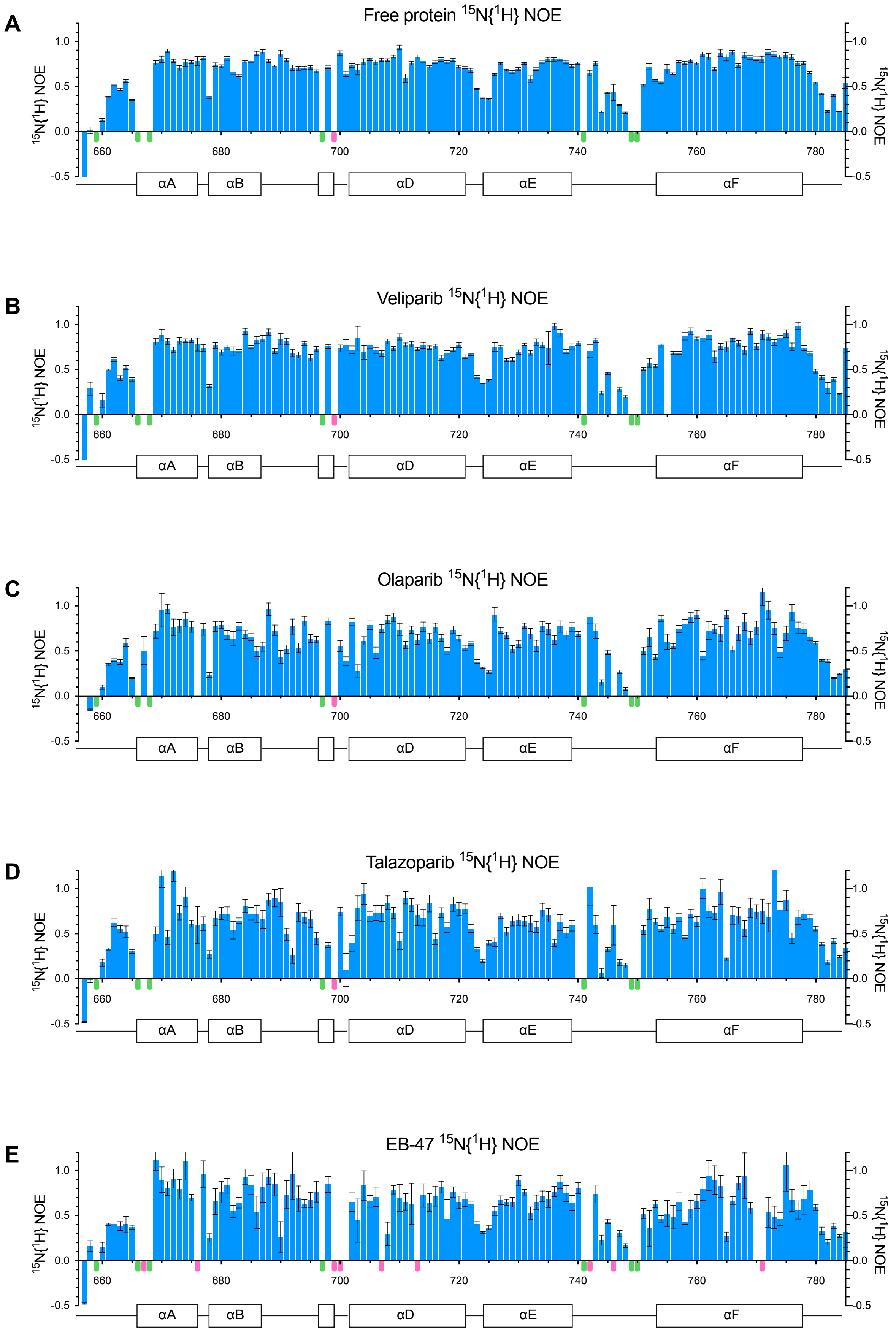


**Figure S4.** Steady-state {$^1$H} $^{15}$N NOE data for A) PARP-1 CAT domain and its complexes with B) veliparib, C) olaparib, D) talazoparib and E) EB-47, showing residues of HD subdomain only. Small coloured bars beneath the sequence scale are used to indicate the positions of prolines (pale green) and overlapped or unassigned signals (pink). Error bars were derived as described in Materials and Methods.

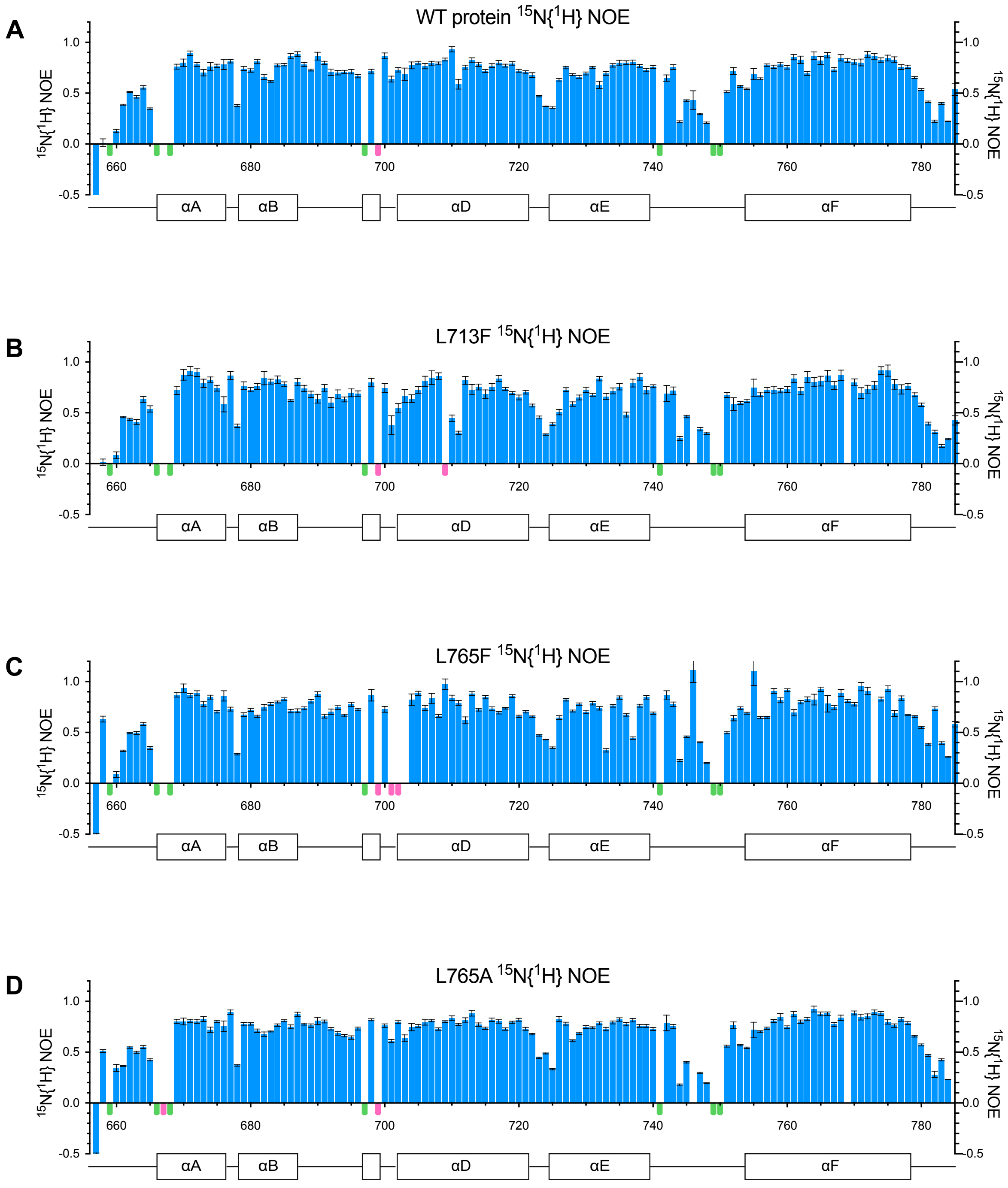


**Figure S5.** Steady-state {$^1$H} $^{15}$N NOE data for A) WT PARP-1 CAT domain and its point mutants B) L713F, C) L765F and D) L765A, showing residues of HD subdomain only. Small coloured bars beneath the sequence scale are used to indicate the positions of prolines (pale green) and overlapped or unassigned signals (pink). Error bars were derived as described in Materials and Methods.

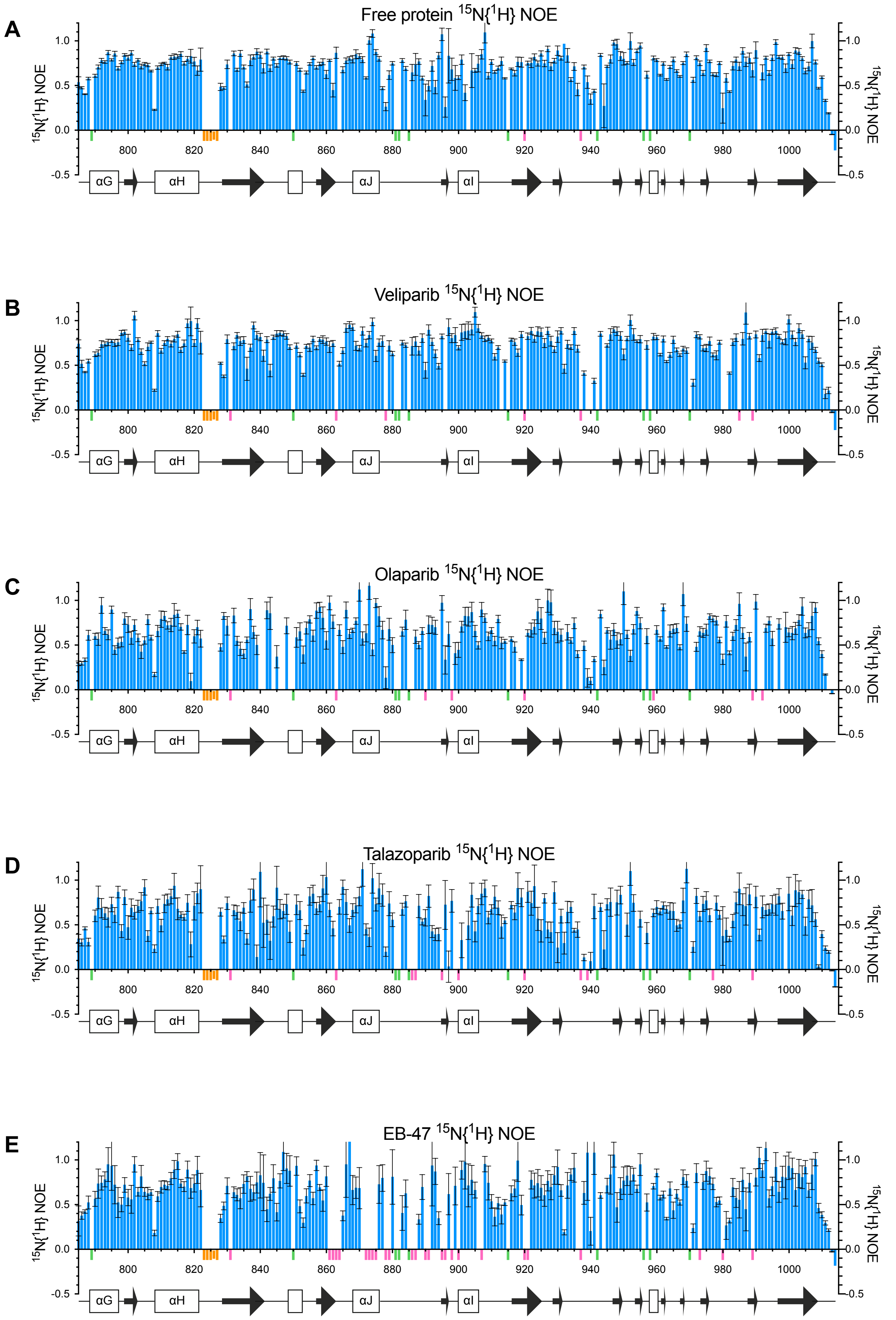


**Figure S6.** Steady-state {$^1$H} $^{15}$N NOE data for A) PARP-1 CAT domain and its complexes with B) veliparib, C) olaparib, D) talazoparib and E) EB-47, showing residues of ART subdomain only. Small coloured bars beneath the sequence scale are used to indicate the positions of prolines (pale green), overlapped or unassigned signals (pink) and the Ala823-Asn827 loop for which no signals were seen in any spectrum (orange). Error bars were derived as described in Materials and Methods.

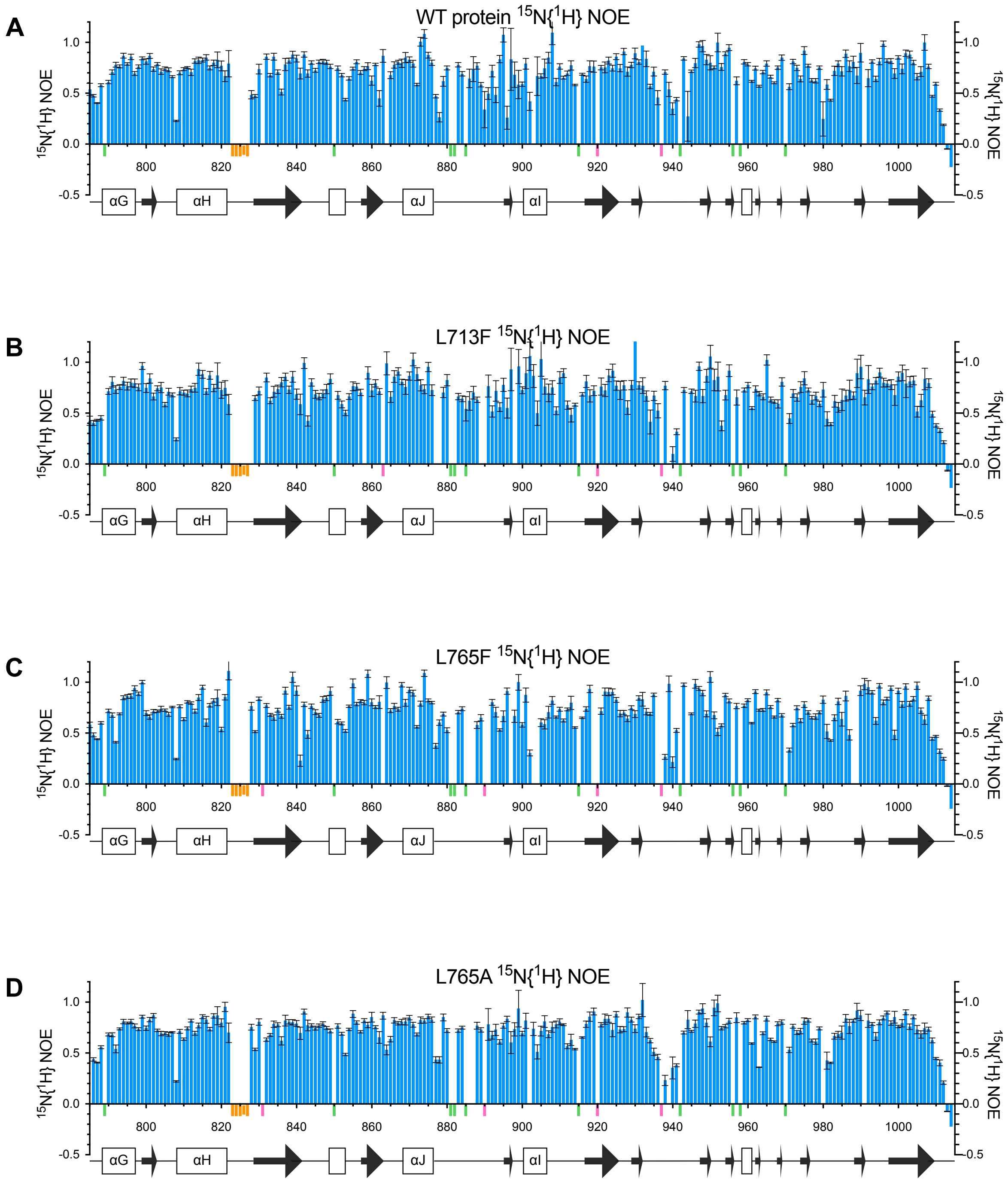


**Figure S7.** Steady-state {$^{1}$H} $^{15}$N NOE data for A) WT PARP-1 CAT domain and its point mutants B) L713F, C) L765F and D) L765A, showing residues of ART subdomain only. Small coloured bars beneath the sequence scale are used to indicate the positions of prolines (pale green), overlapped or unassigned signals (pink) and the Ala823-Asn827 loop for which no signals were seen in any spectrum (orange). Error bars were derived as described in Materials and Methods.

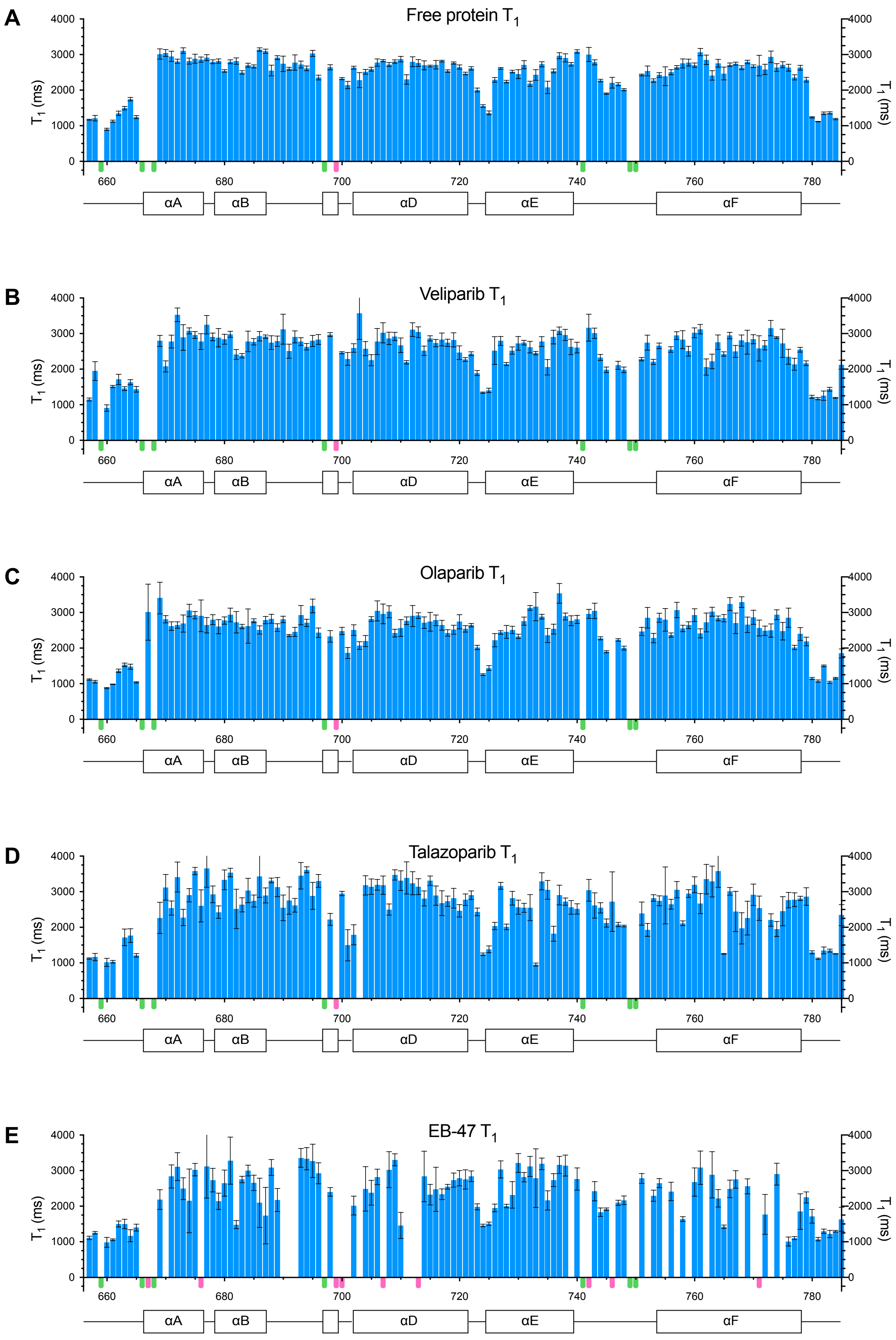


**Figure S8.** [15]N longitudinal relaxation time ($T_1$) data for A) PARP-1 CAT domain and its complexes with B) veliparib, C) olaparib, D) talazoparib and E) EB-47, showing residues of HD subdomain only. Small coloured bars beneath the sequence scale are used to indicate the positions of prolines (pale green) and overlapped or unassigned signals (pink). Error bars were derived as described in Materials and Methods.

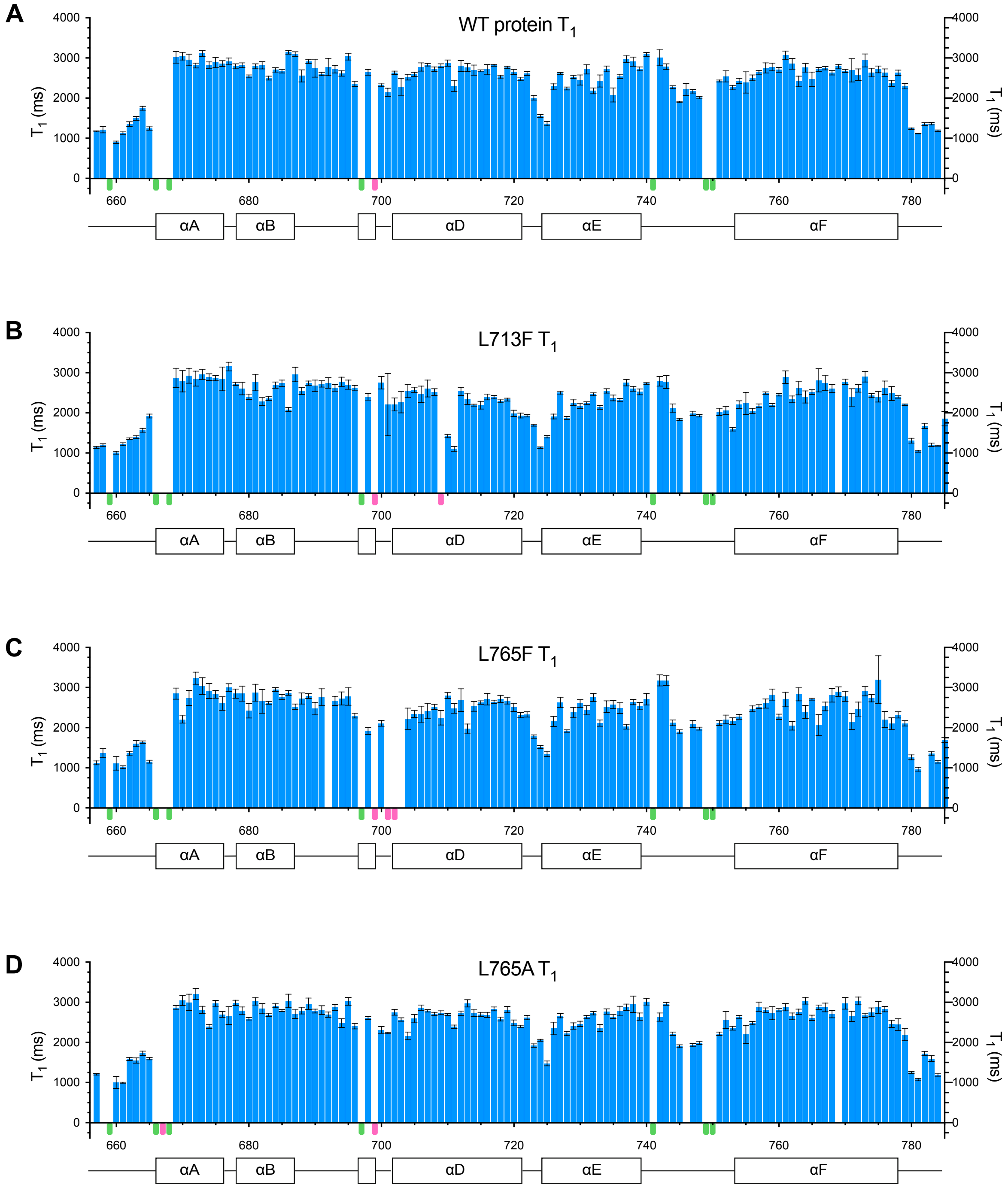


**Figure S9.** $^{15}N$ longitudinal relaxation time ($T_1$) data for A) WT PARP-1 CAT domain and its point mutants B) L713F, C) L765F and D) L765A, showing residues of HD subdomain only. Small coloured bars beneath the sequence scale are used to indicate the positions of prolines (pale green) and overlapped or unassigned signals (pink). Error bars were derived as described in Materials and Methods.

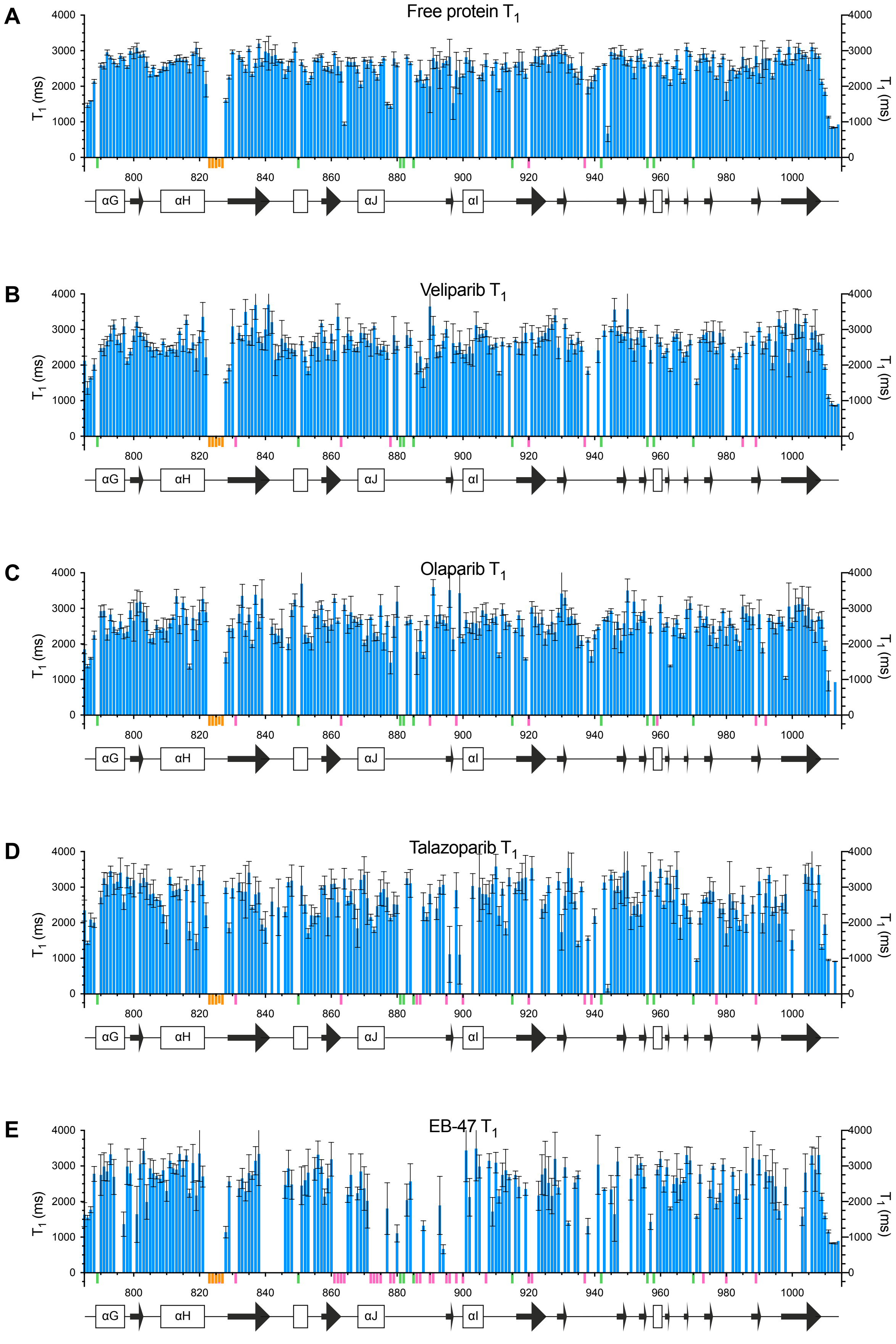


**Figure S10.** $^{15}$N longitudinal relaxation time ($T_1$) data for A) PARP-1 CAT domain and its complexes with B) veliparib, C) olaparib, D) talazoparib and E) EB-47, showing residues of ART subdomain only. Small coloured bars beneath the sequence scale are used to indicate the positions of prolines (pale green), overlapped or unassigned signals (pink) and the Ala823-Asn827 loop for which no signals were seen in any spectrum (orange). Error bars were derived as described in Materials and Methods.

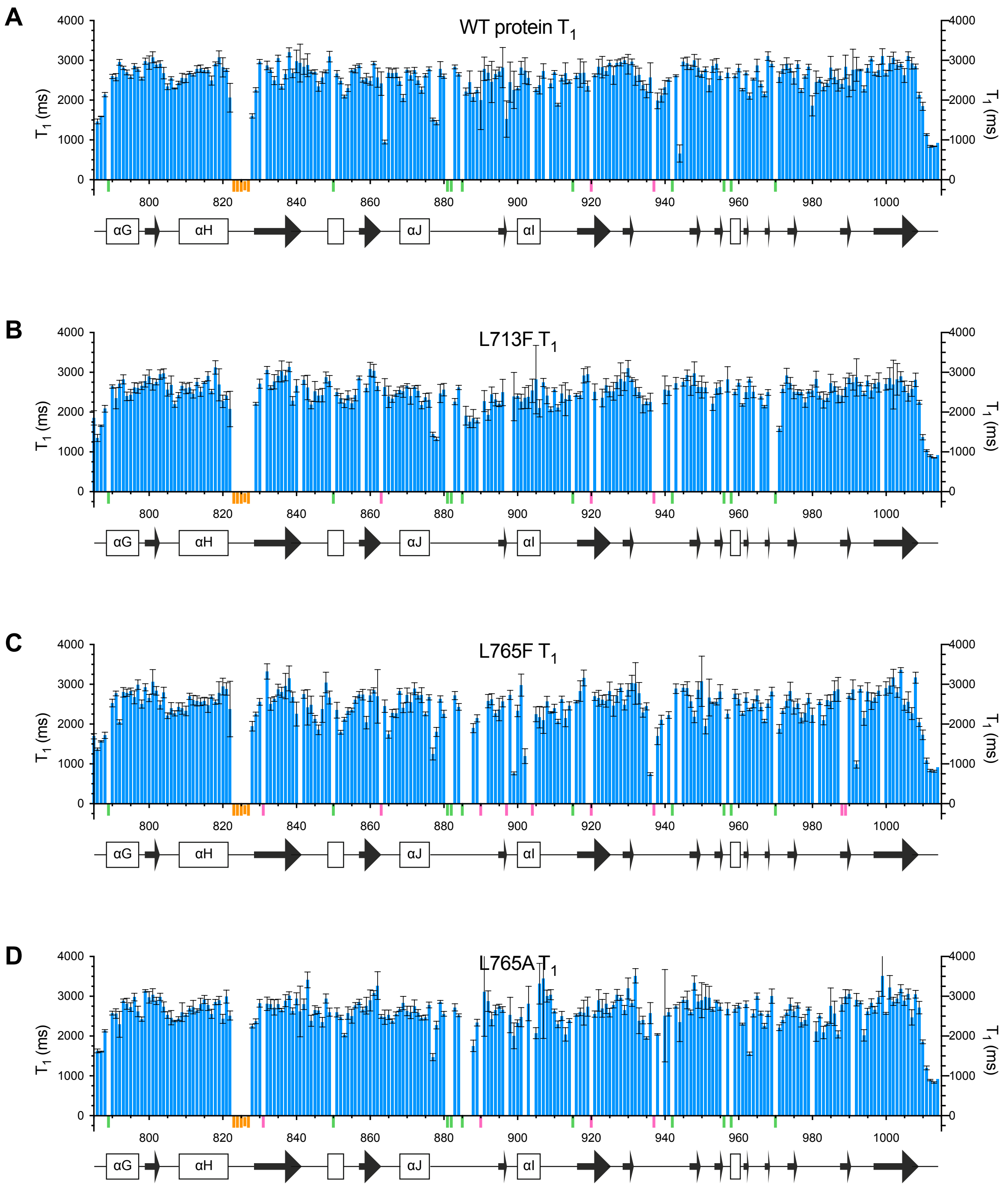


**Figure S11.** $^{15}$N longitudinal relaxation time ($T_1$) data for A) WT PARP-1 CAT domain and its point mutants B) L713F, C) L765F and D) L765A, showing residues of ART subdomain only. Small coloured bars beneath the sequence scale are used to indicate the positions of prolines (pale green), overlapped or unassigned signals (pink) and the Ala823-Asn827 loop for which no signals were seen in any spectrum (orange). Error bars were derived as described in Materials and Methods.

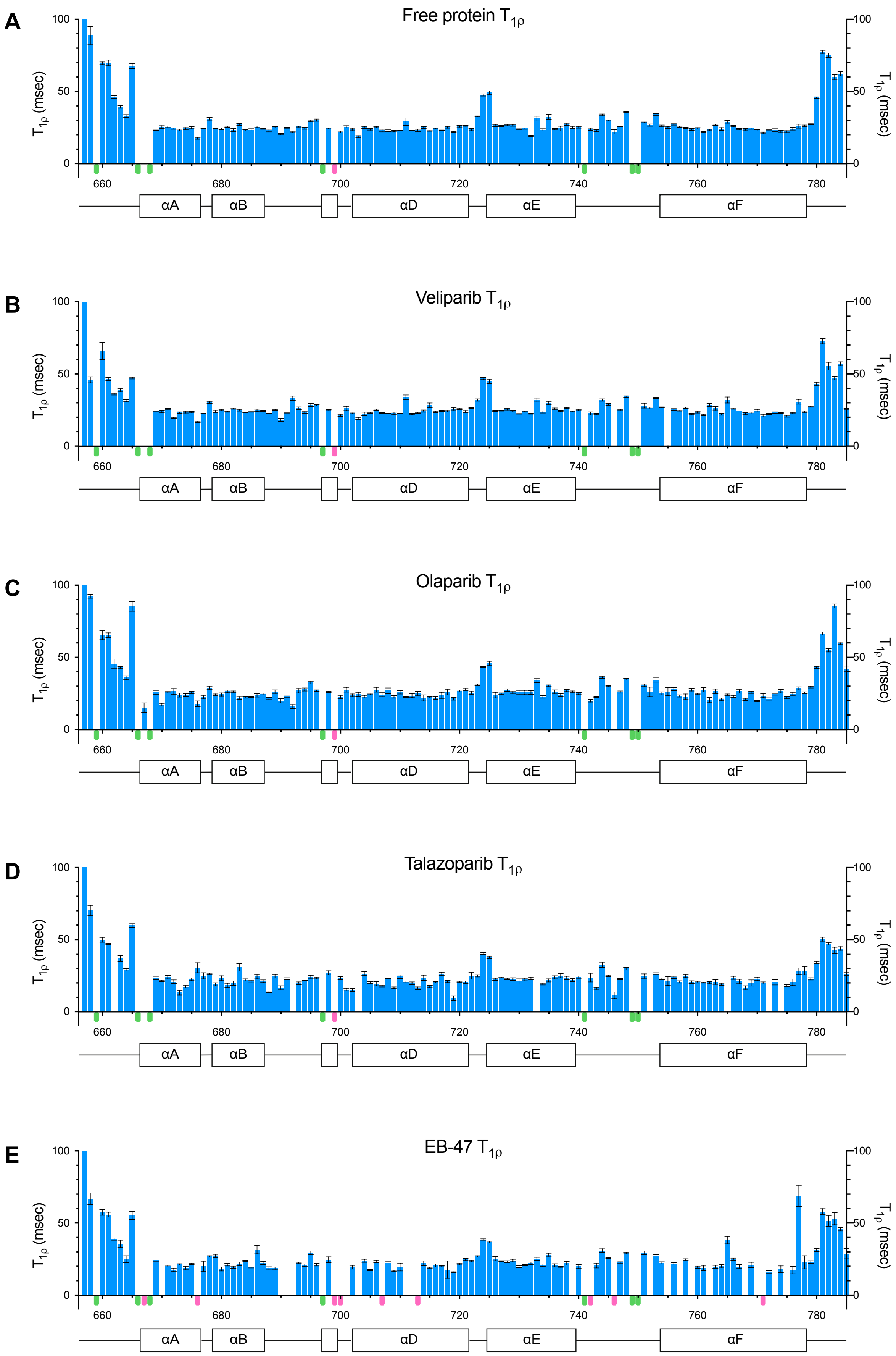


**Figure S12.** $^{15}$N spin-locked relaxation time ($T_{1\rho}$) data for A) PARP-1 CAT domain and its complexes with B) veliparib, C) olaparib, D) talazoparib and E) EB-47, showing residues of HD subdomain only. Small coloured bars beneath the sequence scale are used to indicate the positions of prolines (pale green) and overlapped or unassigned signals (pink). Error bars were derived as described in Materials and Methods.

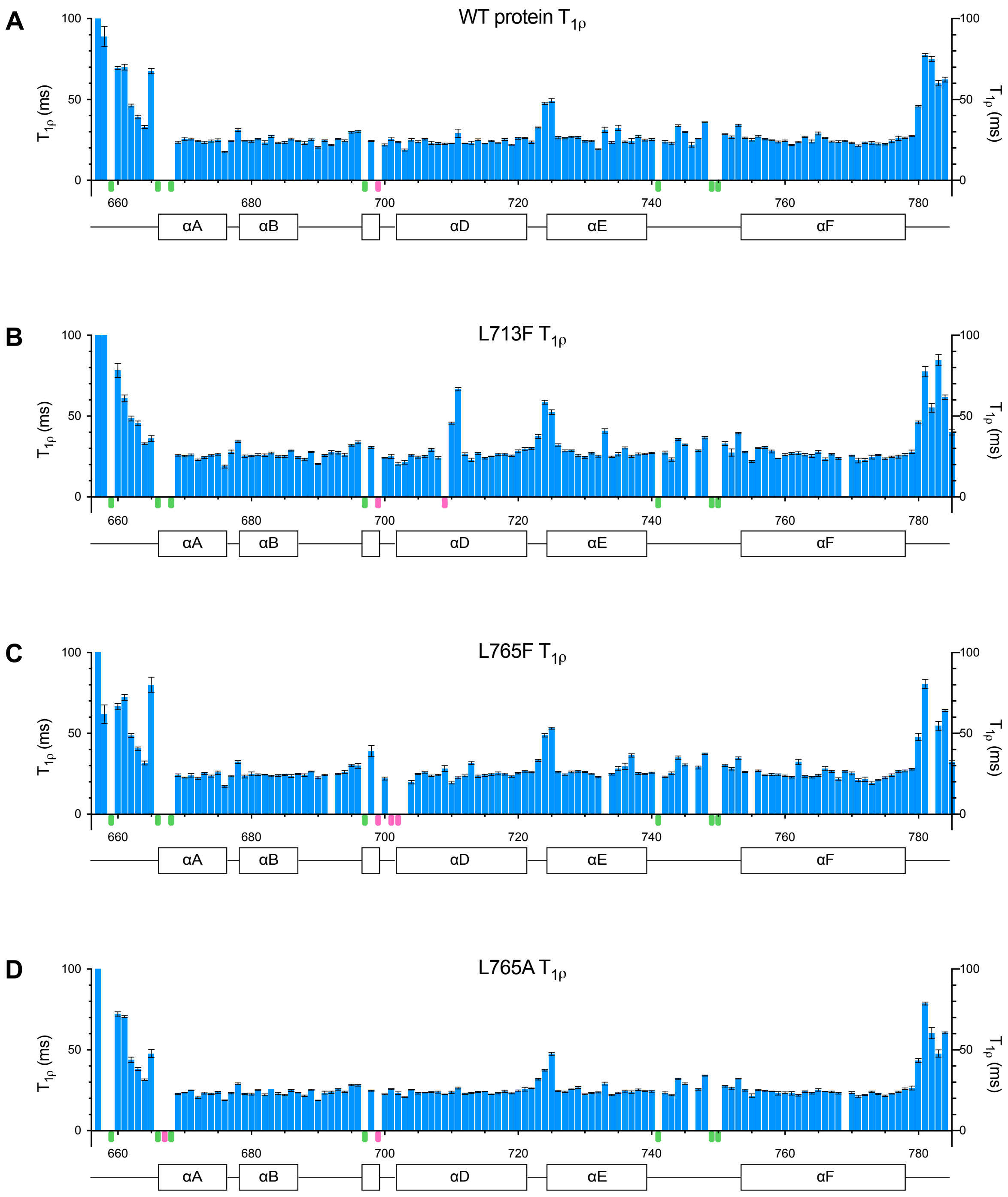


**Figure S13.** [15]N spin-locked relaxation time ($T_{1\rho}$) data for A) WT PARP-1 CAT domain and its point mutants B) L713F, C) L765F and D) L765A, showing residues of HD subdomain only. Small coloured bars beneath the sequence scale are used to indicate the positions of prolines (pale green) and overlapped or unassigned signals (pink). Error bars were derived as described in Materials and Methods.

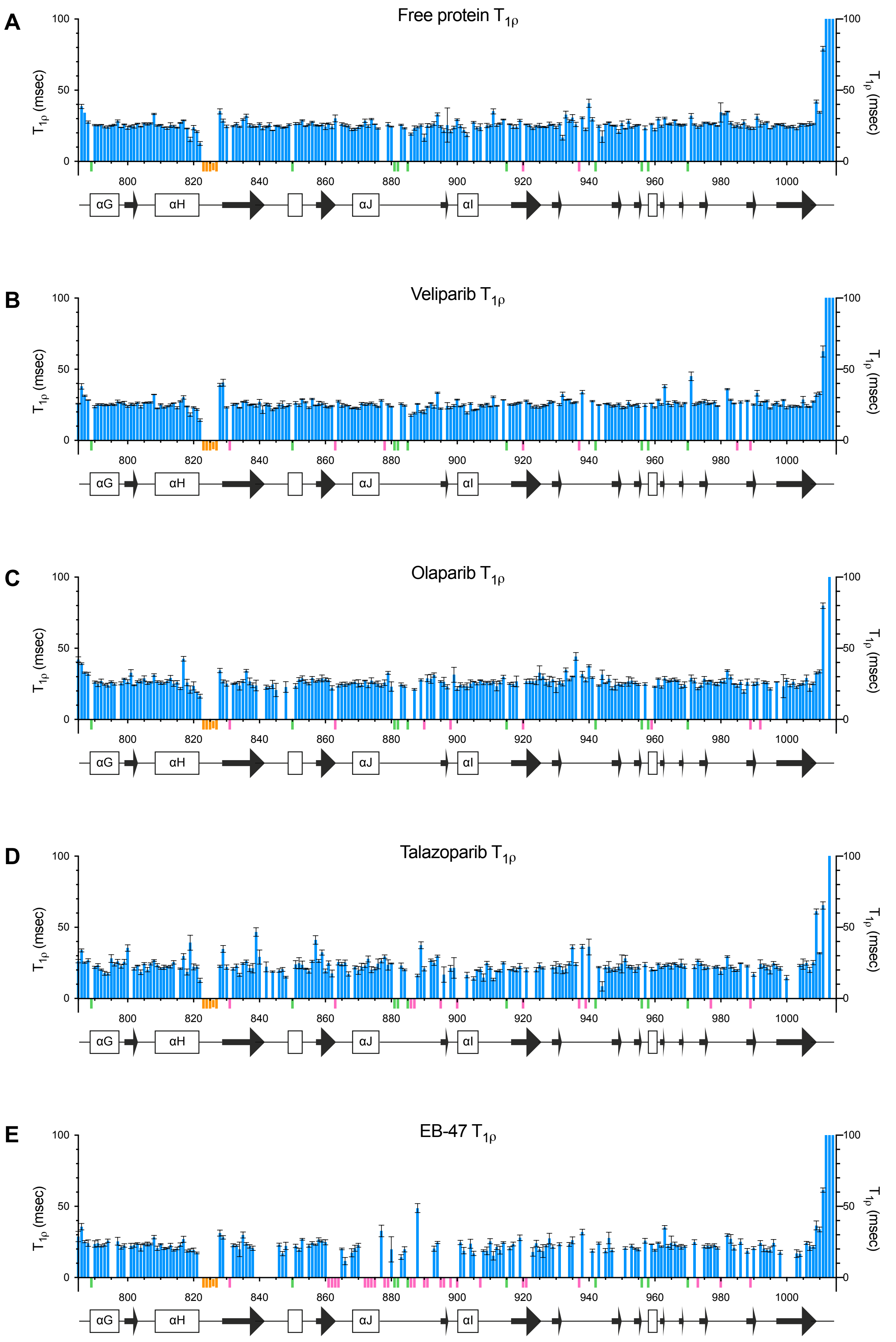


**Figure S14.** $^{15}$N spin-locked relaxation time ($T_{1\rho}$) data for A) PARP-1 CAT domain and its complexes with B) veliparib, C) olaparib, D) talazoparib and E) EB-47, showing residues of ART subdomain only. Small coloured bars beneath the sequence scale are used to indicate the positions of prolines (pale green), overlapped or unassigned signals (pink) and the Ala823-Asn827 loop for which no signals were seen in any spectrum (orange). Error bars were derived as described in Materials and Methods.

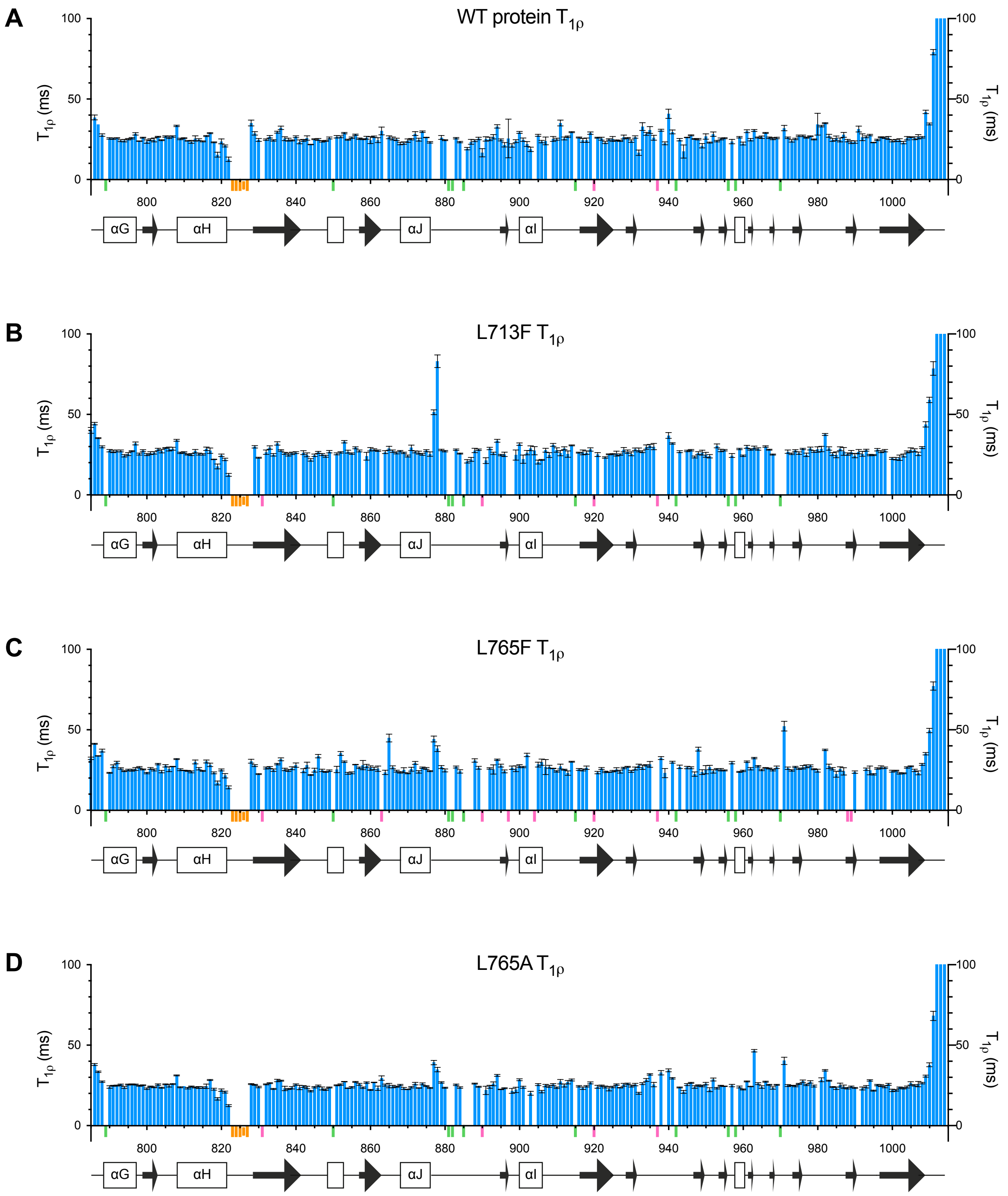


**Figure S15.** [15]N spin-locked relaxation time ($T_{1\rho}$) data for A) WT PARP-1 CAT domain and its point mutants B) L713F, C) L765F and D) L765A, showing residues of ART subdomain only. Small coloured bars beneath the sequence scale are used to indicate the positions of prolines (pale green), overlapped or unassigned signals (pink) and the Ala823-Asn827 loop for which no signals were seen in any spectrum (orange). Error bars were derived as described in Materials and Methods.

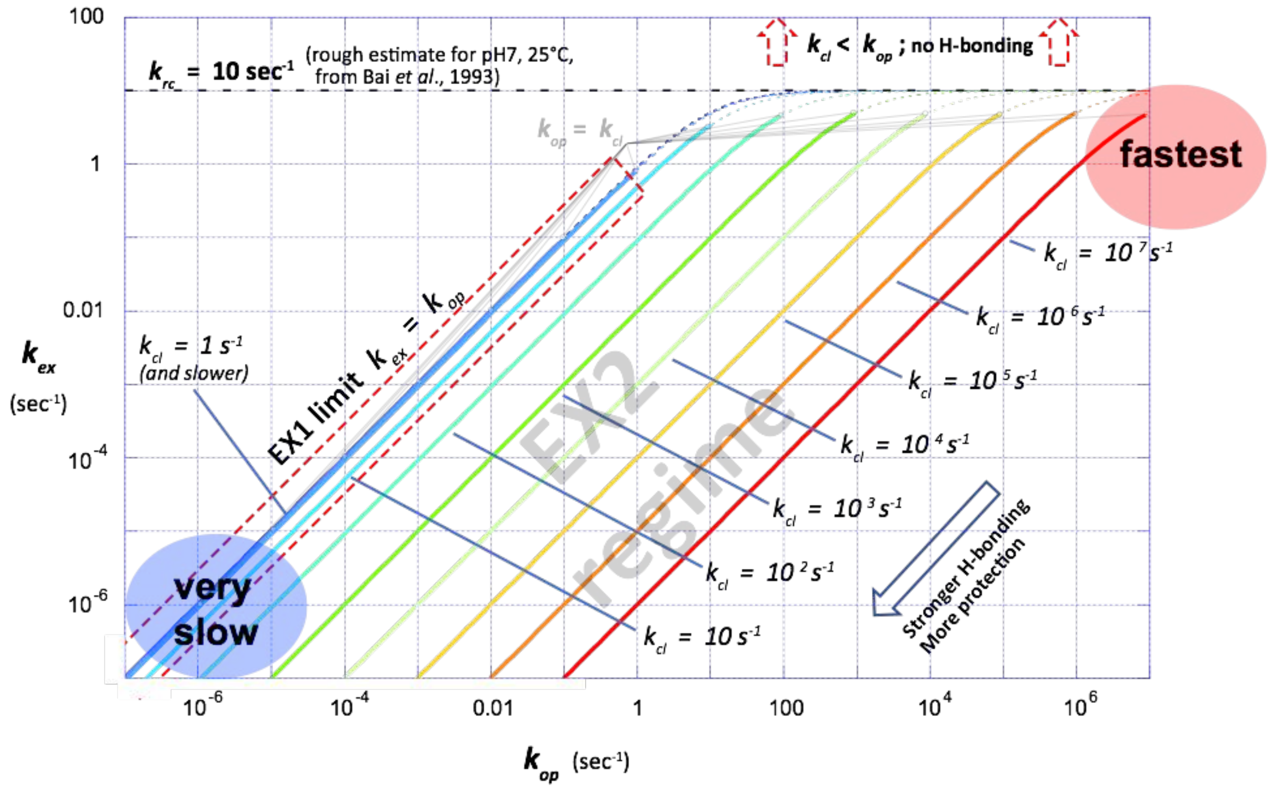


**Figure S16.** Relationship between NH solvent exchange rates and conformational fluctuation rates. The figure shows plots of NH solvent exchange rate constant $k_{ex}$ against the conformational opening rate constant $k_{op}$ calculated using Eq (1) (defined below) for different values of the conformational closing rate constant $k_{cl}$, with all variables in the range $10^{-7}$ to $10^{7}$.

#### *i) Theory:*

The fastest that an NH group can exchange with solvent corresponds to the "random coil" rate, which is the rate for an NH that experiences no protection through intramolecular hydrogen bonding; this is characterised by the rate constant $k_{rc}$ (see point v) below). Participation in intramolecular hydrogen bonds within a folded structure retards this rate to an extent that depends on the strength of the hydrogen bond. This behaviour is usually represented using the simplified reaction scheme:

$$\mathrm{NH_{cl}} \underset{k_{cl}}{\overset{k_{op}}{\rightleftharpoons}} \mathrm{NH_{op}} \xrightarrow{k_{rc}} \mathrm{ND}$$

where $k_{op}$ and $k_{cl}$ are respectively the rate constants for opening and closing of the hydrogen-bonded state and it is assumed that exchange can only take place in the open state; each of these rate constants, and indeed the nature of the open state, is specific for a particular NH (also, each open state could itself comprise an ensemble of states, the properties of which are effectively averaged in this formalism). The solvent exchange rate constant $k_{ex}$ is then given by (1):

$$k_{ex} = \frac{k_{op}k_{rc}}{k_{cl}+k_{op}+k_{rc}} \quad (1)$$

Often this equation appears without the term $k_{op}$ in the denominator, e.g. in (2), corresponding to the assumption that all hydrogen bonds are strong, $k_{cl} \gg k_{op}$; however, here it is of interest to plot the behaviour for both strong and weak hydrogen bonds so the $k_{op}$ term on the lower line is retained. In the figure, those parts of the curves for which $k_{cl} > k_{op}$ (implying that the closed form containing the hydrogen bond predominates) are shown using thick solid lines, while those parts for which $k_{cl} < k_{op}$ (implying that the open form where there is no hydrogen bond predominates) are shown using thin dotted lines.

Because solvent exchange is a chemical reaction, it shows strong dependence on solution conditions, particularly pH and temperature, and in addition is sensitive to the chemical nature of nearby sidechains. For an amide group in a random coil conformation, i.e. one that is "unprotected" as it participates only in transient hydrogen bonds with solvent, the random coil rate solvent exchange constant $k_{rc}$ can be roughly predicted from model compound studies (3); for a protein at pH 7.0 and 25 °C, $k_{rc}$ would be expected to be very roughly 10 $s^{-1}$, with variations of up to about a factor of ten in either direction depending on sequence context. This is the value that was used in the simulations shown in the figure; for more details see point vi) below.

#### *ii) EX2 and EX1 limits:*

Two limiting cases are usually considered, which differ according to whether it is the first step (structural opening) or second step (exchange) that is rate-limiting; the first limit is referred to as EX1 and the second as EX2 (4). By far the commonest situation is for exchange to be in or near the EX2 limit, where structural fluctuations are much faster than intrinsic exchange, $k_{cl} + k_{op} \gg k_{rc}$ , so that Eq. 1 becomes:

$$k_{ex} = \frac{k_{op}}{k_{cl}+k_{op}} k_{rc} = f_{op} k_{rc} \qquad \text{(EX2)} \qquad (2)$$

In effect, this implies that an equilibrium is rapidly established between the closed and open states, and the observed exchange rate then reflects both the fractional abundance of the opened state ($f_{op}$) and the rate of exchange events it undergoes once formed. For hydrogen bonds strong enough that $k_{cl} \gg k_{op}$ , the EX2 condition is often written as

$$k_{ex} = \frac{k_{op}}{k_{cl}} k_{rc} = K_{op} k_{rc} \qquad \text{(EX2)} \qquad (3)$$

where $K_{op}$ is the equilibrium constant for the pre-equilibrium between open and closed states.

For extremely slow rates, exchange can reach the EX1 limit, $k_{cl} + k_{op} \ll k_{rc}$ , and then Eq. 1 becomes simply:

$$k_{ex} = k_{op} \qquad \text{(EX1)} \qquad (4)$$

On the figure, the EX1 limit manifests as the coalescence into a single line of all of the curves for which $k_{cl} < 1\ s^{-1}$, in the region marked with a dashed red box. The EX2 limit is less obviously visualisable on the figure, but corresponds to those curves for which $k_{cl} > 100\ s^{-1}$ (i.e. the bulk of the remainder of the plot), where increasing $k_{cl}$ leads to a proportionate movement of the $k_{ex}$ vs $k_{op}$ line. It may be noted that, formally, Eq. 1 is an approximation, since between the EX1 and EX2 limits (here corresponding approximately to the range $1\ s^{-1} < k_{cl} < 100\ s^{-1}$) exchange follows a bi-exponential rate that cannot be represented analytically using a single rate constant (5); however, this complication is often ignored.

#### *iii) Fastest events:*

The NH protons that undergo the fastest solvent exchange are those situated in particularly mobile loops, where they participate only very weakly or not at all in intra-molecular hydrogen bonding; they are largely or completely unprotected from solvent exchange and consequently exchange at or near the random-coil rate. Under the conditions of the present study (25 °C and pH7), $k_{rc}$ is roughly 10 $s^{-1}$ (see vi below). However, the rate of underlying conformational motions for such protons is very much faster. For such motions to be detected in $^{15}$N relaxation experiments, they must be faster than overall molecular tumbling, which for PARP-1 CAT domain at 25 °C in the present study has a correlation time of $\tau_c \approx 25$ ns (see main text).

In other words, in such cases there are *at least seven orders of magnitude* difference between the rates of NH proton solvent exchange and the underlying rates of conformational exchange events involving those protons. This corresponds roughly to the part of the plot highlighted in red labelled "fastest". In this regime it would be expected that the conformational motions of individual peptide groups are largely un-coordinated of with those their neighbours, and any transient making or breaking of intramolecular hydrogen bonds involving these peptides would probably be largely or completely non-cooperative in nature.

##### *iv) Slowest events:*

In contrast, NH protons that exchange with solvent the most slowly can be at the EX1 limit, where the rate of solvent NH proton exchange is *equal to* the rate of the underlying conformational processes responsible for transiently breaking hydrogen bonds. Such cases could lie within the part of the plot highlighted in blue labelled "very slow" (though still slower cases of exchange corresponding to still stronger hydrogen bonds, and which would lie outside the limits of this plot, are also entirely possible). Hydrogen bonds involved in these processes are very strong indeed, and the processes that break them are likely to be highly co-operative in nature, involving the disruption of large sections of the structure (2,6,7); in the limit, for NH protons involved in the very strongest hydrogen bonds in the most stable of protein structures, solvent exchange can require global unfolding of the entire structure (8). Cases of EX1 exchange are relatively rare, although working at high pH makes EX1 more likely as it increases greatly the random coil exchange rate (2).

##### *v) Between the extremes:*

The great majority of hydrogen-bonded NH protons characterised in exchange studies fall between the limiting behaviours just described, and are mostly within the EX2 regime. Generalisations are difficult, but it seems reasonable to expect that the trend from fastest to slowest solvent exchange would correspond very approximately to a trend from least to most co-operativity in the conformational processes underlying exchange; for the great majority of hydrogen bonds, the conformational fluctuations required for transient opening will involve some other portions of nearby structure that are interconnected in the hydrogen bonding network, and the stronger the hydrogen bond, the larger the "structural reach" of such co-operative events is likely to have to be (8). Implications for the relationship between solvent exchange rates and rates of the underlying conformational fluctuations causing exchange are also hard to predict, but one may expect that the slower the exchange, the smaller is likely to be the extent by which conformational fluctuation rates are faster than solvent exchange. Even so, the EX2 regime covers a very broad range of NH solvent exchange rates, and in the majority of cases it is likely that the underlying conformational fluctuations that drive exchange are significantly faster than solvent exchange itself, potentially by orders of magnitude.

Without measured data concerning $k_{op}$ and $k_{cl}$, quantitative statements are impossible. However, the purpose of the analysis presented here is to extend the caution given elsewhere in the literature, e.g. in (2,7), against assuming that measured solvent exchange rates are similar to the rates of underlying conformational transitions responsible for that exchange; this is highly unlikely to be the case except for the very slowest exchanging NH protons that are at or near the EX1 limit.

##### *vi) Random coil exchange rate at pH7 and 25 °C:*

The prediction for $k_{rc}$ used in the simulations was obtained from the data and analysis in Bai *et al*. (3) as follows: For a random coil amide group, the solvent NH exchange rate constant uncorrected for neighbouring sidechain effects is given by

$$k_{rc} = k_{ex} = k_A\ 10^{-pD} + k_B\ 10^{[pD-pK_D]} + k_W \qquad (4)$$

where $k_A$ is the rate constant for acid-catalysed exchange, $k_B$ is that for base catalysed exchange and $k_W$ is that for water catalysed exchange. At 20 °C, the temperature for which Bai *et al*. give tabulated data, the value of $K_D$ (the self-ionisation constant for $D_2O$) is 15.049 (9). At pH 7, catalysis by acid and water can be neglected, so after conversion to log units Eq. (4) becomes:

$$\log(k_{rc}) = \log(k_B) + pD - pK_D$$
$$= 10.36 + 7.4 - 15.05 = 2.71$$

where $\log(k_B)$ = 10.36 comes from Bai *et al*. Table 3 (note that rates in Bai *et al*. are all expressed in $min^{-1}$) and 7.4 is the value of pD corresponding to an uncorrected pH meter reading of 7.0 made with a conventional glass electrode (10). From this,

$$k_{rc} = 10^{2.62} = 513\ min^{-1} = 8.55\ s^{-1}.$$

Corrections for local sequence (see Bai *et al*. Table 2) can cause a maximal negative variation in $k_B$ of -0.97 log units (for the NH of Ile in the sequence Ile-Pro) and maximal positive variation in $k_B$ of +1.45 log units (for the NH of Cys in the sequence Cys-$His^+$). These corrections would correspond to a range in the rates between 0.91 $s^{-1}$ and 241 $s^{-1}$. The average over all sidechain corrections (ignoring COOH forms, as these would be absent at pH 7) would give -0.02 to +0.14 log units, which corresponds to a range in the rates between 8.16 $s^{-1}$ and 11.8 $s^{-1}$.

Using the Arrhenius equation to correct these values from a temperature of 20 °C to 25 °C gives:

$$k_{rc}(T) = k_{rc}(293)\ exp^{(-E_a[1/T-1/293]/R)}$$

where $E_a$ is the activation energy for base-catalysed exchange, given as 17 kcal.$mol^{-1}$ by Bai *et al.*, and $R$ is the gas constant, 1.985 × $10^{-3}$ kcal.$K^{-1}$.$mol^{-1}$, giving a correction factor of:

$$exp^{\left(\frac{-17[1/298-1/293]}{0.001985}\right)} = exp^{0.49} = 1.63$$

Thus, the random coil exchange rate constant at 25°C, uncorrected for sequence variations, is approximately 13.9 $s^{-1}$. This value was rounded to 10 $s^{-1}$ for use in the simulations.

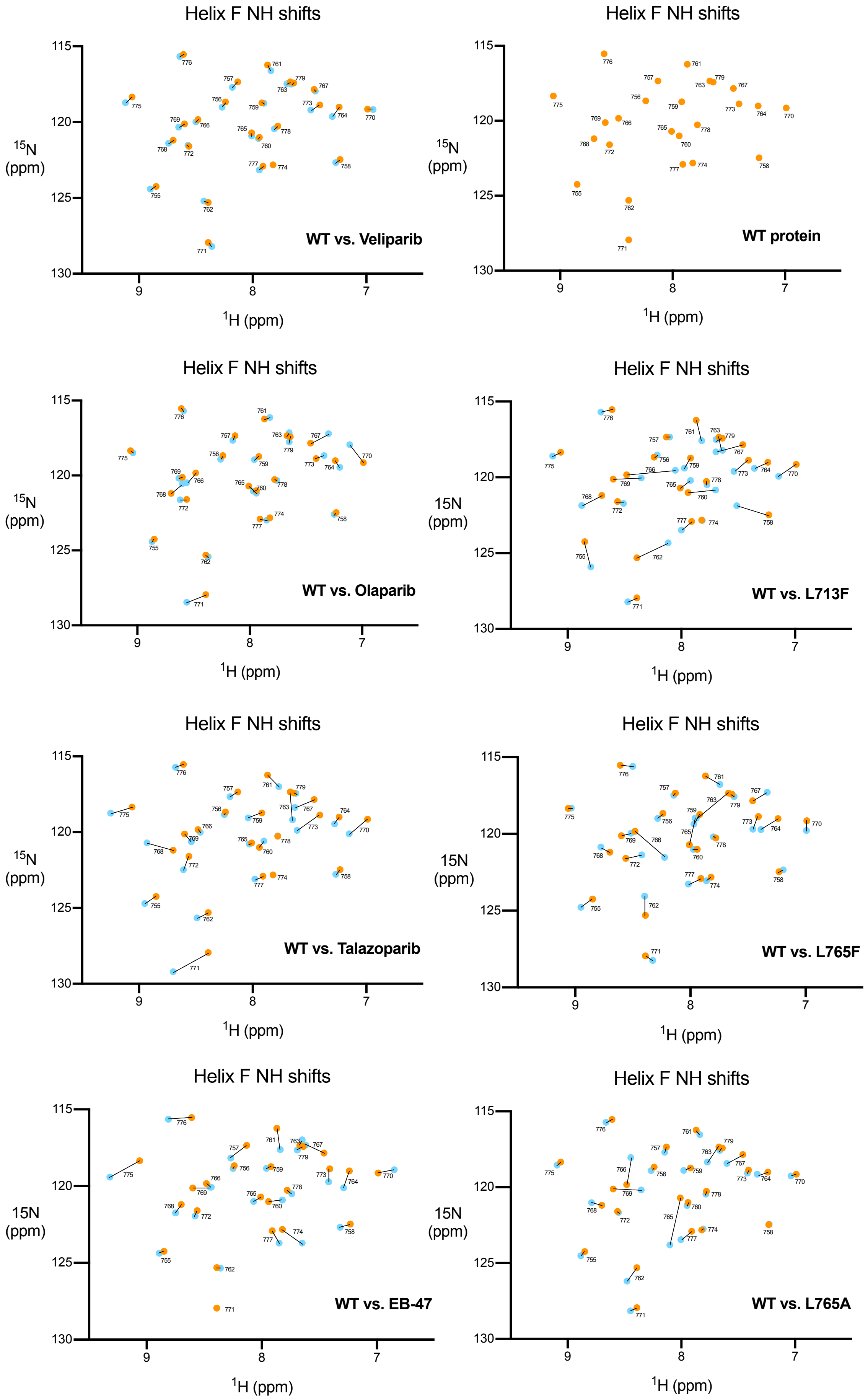


**Figure S17.** Schematic [$^{15}$N,$^{1}$H] correlation plots showing chemical shift differences for the amide group $^{1}$H and $^{15}$N signals of residues in helix F for the comparisons indicated on the individual panels; in each case the WT free protein signal is indicated in orange and the mutant or inhibitor complex signal in light blue. The corresponding schematic for helix F of WT protein alone is also shown.

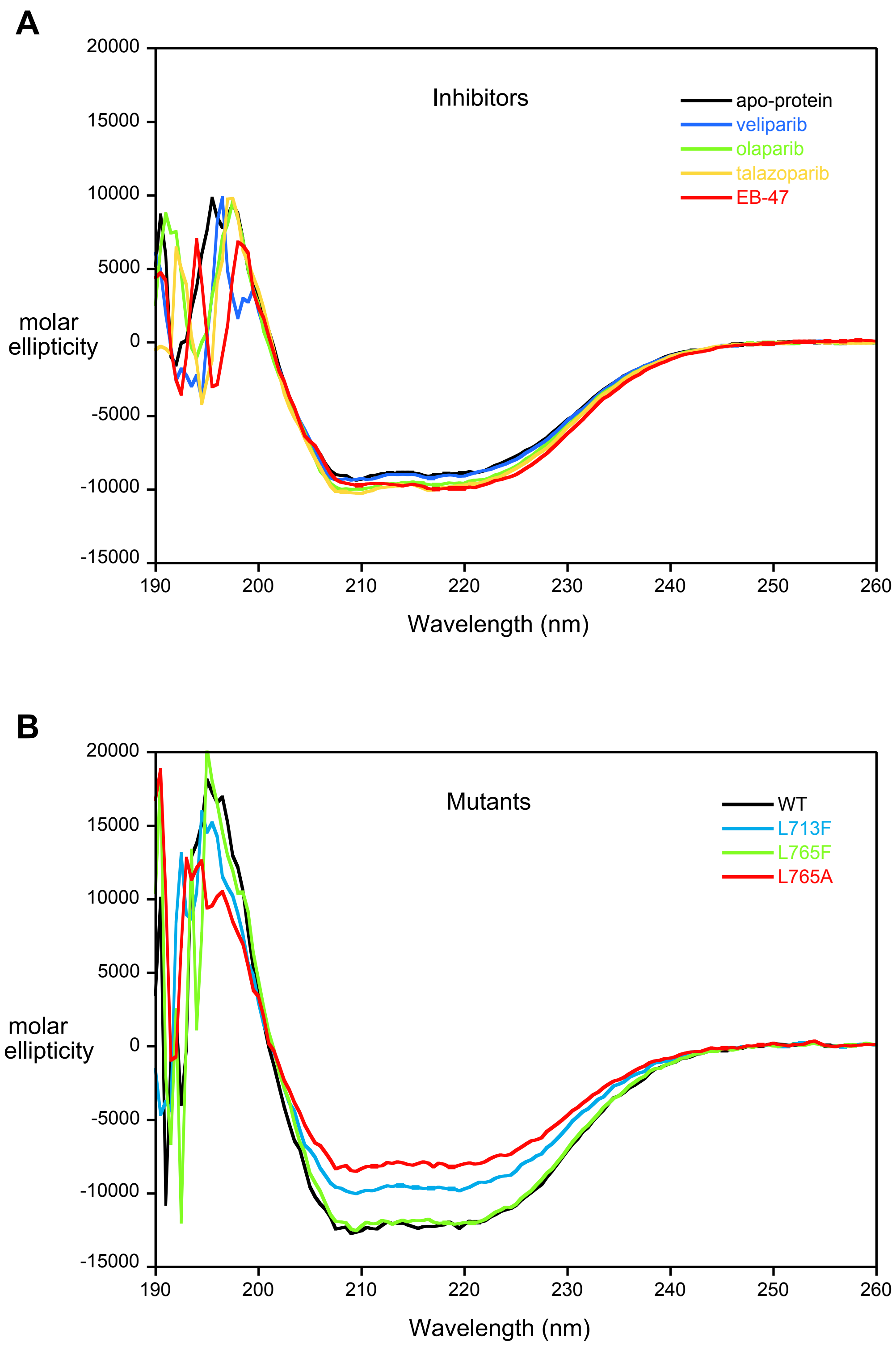


**Figure S18.** CD spectra of PARP-1 CAT domain, and A) its complexes with veliparib, olaparib, talazoparib and EB-47, and B) the point mutants L713F, L765F and L765A, showing that no large-scale perturbation of secondary structure results from binding of these inhibitors or from these mutations. Apparent scaling differences amongst the spectra of the mutants probably reflect errors in estimating concentrations between these different protein samples, whereas the curves for inhibitor complexes all rely on a single protein concentration estimate.

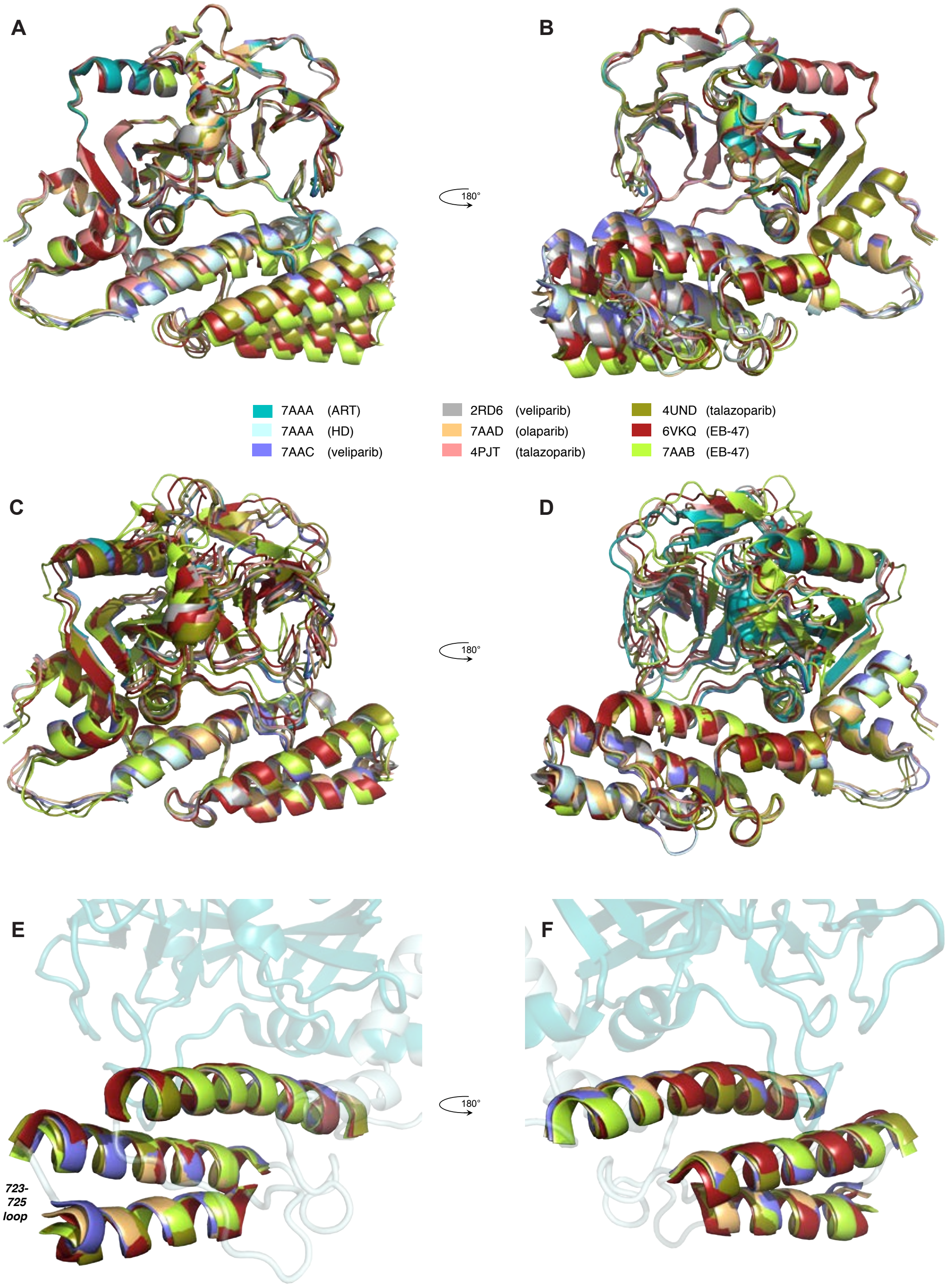


**Figure S19.** Superpositions of PARP-1 CAT domain in the apo-protein and in inhibitor complexes with veliparib (PDB:7AAC and 2RD6), olaparib (PDB:7AAD), talazoparib (PDB:4PJT and 4UND) and EB-47 (PDB:7AAB and PDBB:6VKQ), in each case superposing each structure onto atoms of 7AAA. Superpositions were carried out using backbone atoms of the ART subdomain (N, Cα, C' of residues 790-936, 939-1009) (panels A and B), backbone atoms of the HD subdomain (N, Cα, C' of residues 667-721,730-743,750-779) (panels C and D), or backbone atoms of helices D, E and F (N, Cα, C' of residues 702-721,730-740,754-779) (panels E and F). N.B. The viewpoints in panels E and F are changed relative to those in panels A-D so as to facilitate viewing the helical distortions.

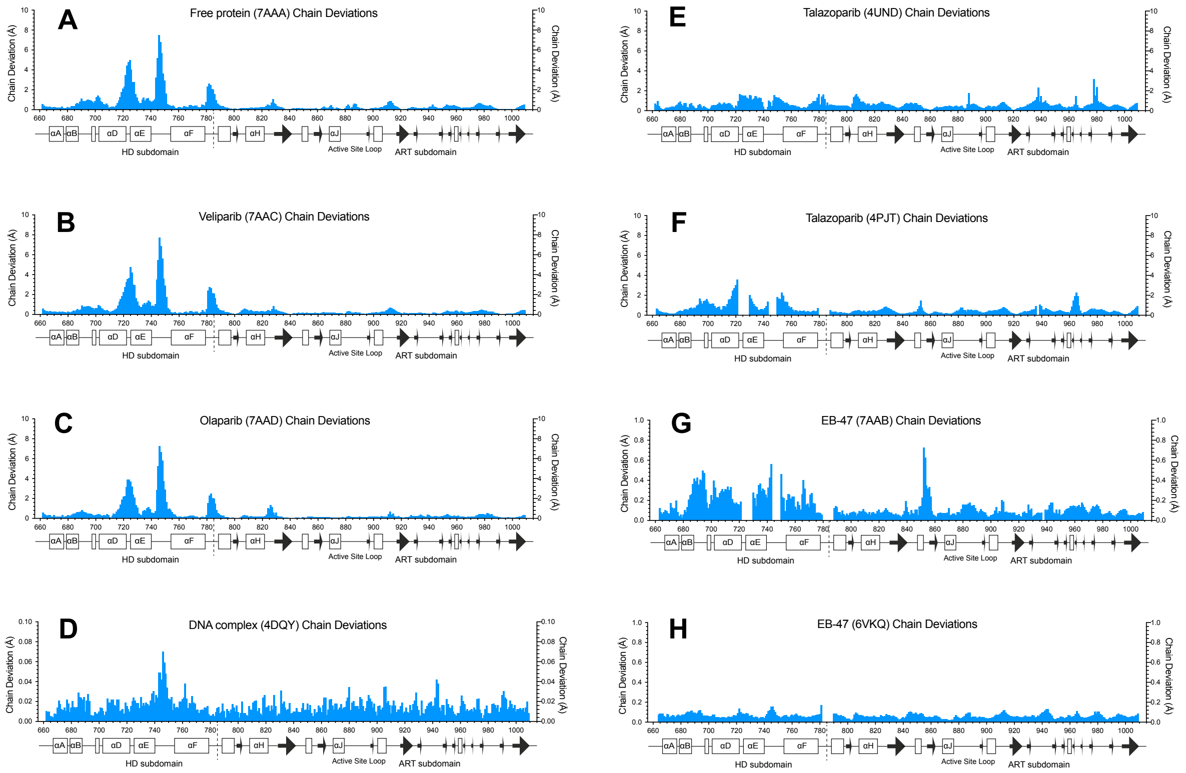


**Figure S20.** Co-ordinate deviations between the different protein chains in the asymmetric unit for the structures of free PARP-1 CAT domain (7AAA, panel A) and its complexes with talazoparib (4UND and 4PJT, panels B and D), veliparib (7AAC, panel C; the other structure of veliparib, 2RD6, has only one protein chain in the asymmetric unit), olaparib (7AAD, panel E), and EB-47 (7AAB and 6BVQ, panels F and H), as well as for chains C and F of the complex of PARP-1 F1, F3 and WGR-CAT with a DNA duplex (4DQY, panel G). In each case, the values shown were calculated by first determining the best ensemble superposition of all of the protein chains in the asymmetric unit for the relevant structure using the program CLUSTERPOSE ["Co-ordinate-based cluster analysis", R. Diamond (1995), Acta Cryst D., 51, 125-135], and then calculating separately for each residue in the sequence the average deviation across all occurrences of the backbone atoms N, $C^{\alpha}$ and C'. (The relatively uniform lower values seen for structures 4UND (panel E) and 6VKQ (panel G) likely reflect the use of strongly weighted non-crystallographic symmetry terms during refinement.)

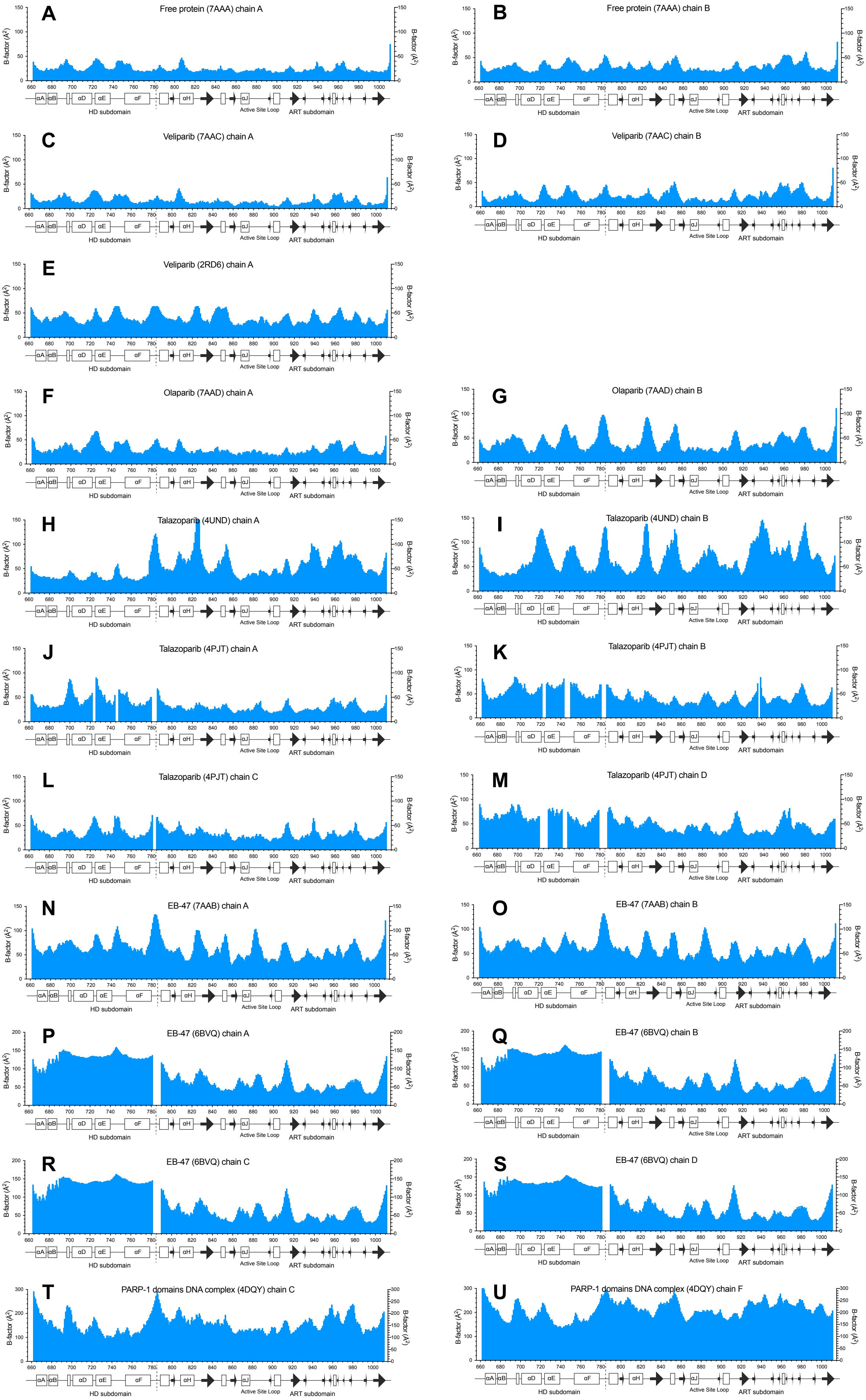
A Free protein (7AAA) chain A
B Free protein (7AAA) chain B
C Veliparib (7AAC) chain A
D Veliparib (7AAC) chain B
E Veliparib (2RD6) chain A
F Olaparib (7AAD) chain A
G Olaparib (7AAD) chain B
H Talazoparib (4UND) chain A
I Talazoparib (4UND) chain B
J Talazoparib (4PJT) chain A
K Talazoparib (4PJT) chain B
L Talazoparib (4PJT) chain C
M Talazoparib (4PJT) chain D
N EB-47 (7AAB) chain A
O EB-47 (7AAB) chain B
P EB-47 (6BVQ) chain A
Q EB-47 (6BVQ) chain B
R EB-47 (6BVQ) chain C
S EB-47 (6BVQ) chain D
T PARP-1 domains DNA complex (4DQY) chain C
U PARP-1 domains DNA complex (4DQY) chain F

**Figure S21.** B factors for each chain in each of the structures of free PARP-1 CAT domain (7AAA, panels A and B) and its complexes with veliparib (7AAC, panels C and, and 2RD6, panel E), olaparib (7AAD, panels F and G), talazoparib (4UND, panels H and I, and 4PJT, panels J - M), and EB-47 (7AAB, panels N and O, and 6BVQ, panels P - S), as well as for chains C and F of the complex of PARP-1 F1, F3 and WGR-CAT with a DNA duplex (4DQY, panels T and U). Note the different vertical scales: panels A-O run to 150 $Å^2$, panels P-S run to 200 $Å^2$, and panels T and U run to 300 $Å^2$.

**Table S1.** Superposition statistics for individual helices of PARP-1 CAT domain (all chains fitted onto 7AAA).

| **Fitted onto 7AAA chain A** | **Helix A** | **Helix B** | **Helix D** | **Helix E** | **Helix F** | **Helix G** | **Helix H** | **Helix I** | **Helix J** | **Helix K** |
|---|---|---|---|---|---|---|---|---|---|---|
| *sequence* | *666-677* | *679-688* | *702-722* | *725-740* | *754-779* | *788-797* | *808-821* | *848-853* | *868-876* | *900-906* |
| *# residues* | *12* | *10* | *21* | *16* | *26* | *10* | *14* | *6* | *9* | *7* |
| | (Å) | (Å) | (Å) | (Å) | (Å) | (Å) | (Å) | (Å) | (Å) | (Å) |
| 2PAW A | 0.129 | 0.106 | 0.359 | 1.050 | 0.453 | 0.227 | 0.165 | 0.667 | 0.143 | 0.211 |
| 1A26 A | 0.147 | 0.104 | 0.383 | 1.085 | 0.494 | 0.200 | 0.161 | 0.663 | 0.126 | 0.115 |
| 4DQY C | 0.235 | 0.561 | 0.477 | 1.024 | 0.834 | 0.401 | 0.348 | 0.566 | 0.267 | 0.222 |
| 4DQY F | 0.241 | 0.564 | 0.476 | 1.021 | 0.832 | 0.403 | 0.348 | 0.565 | 0.267 | 0.217 |
| 7AAC A | 0.047 | 0.038 | 0.122 | 0.101 | 0.117 | 0.054 | 0.069 | 0.045 | 0.044 | 0.052 |
| 7AAC B | 0.073 | 0.038 | 0.502 | 1.131 | 0.246 | 0.083 | 0.088 | 0.037 | 0.047 | 0.058 |
| 2RD6 | 0.139 | 0.143 | 0.294 | 1.038 | 0.416 | 0.227 | 0.188 | 0.657 | 0.092 | 0.253 |
| 7AAD A | 0.086 | 0.068 | 0.161 | 0.487 | 0.177 | 0.104 | 0.106 | 0.067 | 0.070 | 0.082 |
| 7AAD B | 0.106 | 0.065 | 0.488 | 1.113 | 0.260 | 0.124 | 0.124 | 0.111 | 0.068 | 0.109 |
| 4PJT A | 0.112 | 0.127 | 0.320[a] | 0.198[b] | 0.506 | 0.133 | 0.180 | 0.160 | 0.090 | 0.091 |
| 4PJT B | 0.100 | 0.077 | 0.436[a] | 0.231[b] | 0.301 | 0.165 | 0.153 | 0.471 | 0.087 | 0.101 |
| 4PJT C | 0.093 | 0.089 | 0.428[a] | 0.194[b] | 0.297 | 0.172 | 0.150 | 0.444 | 0.080 | 0.093 |
| 4PJT D | 0.098 | 0.117 | 0.352[a] | 0.288[b] | 0.476 | 0.139 | 0.144 | 0.156 | 0.081 | 0.097 |
| 4UND A | 0.151 | 0.110 | 0.334 | 0.980 | 0.315 | 0.188 | 0.164 | 0.533 | 0.111 | 0.232 |
| 4UND B | 0.129 | 0.118 | 0.409 | 1.053 | 0.395 | 0.173 | 0.194 | 0.540 | 0.111 | 0.232 |
| 7AAB A | 0.211 | 0.140 | 0.466 | 0.999 | 0.362 | 0.223 | 0.187 | 0.181 | 0.156 | 0.151 |
| 7AAB B | 0.230 | 0.128 | 0.463 | 0.893 | 0.366 | 0.218 | 0.185 | 0.278 | 0.157 | 0.147 |
| 6VKQ A | 0.237 | 0.259 | 0.989 | 0.818 | 0.548 | 0.215[c] | 0.400 | 0.379 | 0.168 | 0.177 |
| 6VKQ B | 0.236 | 0.258 | 0.997 | 0.818 | 0.548 | 0.214[c] | 0.400 | 0.378 | 0.168 | 0.176 |
| 6VKQ C | 0.237 | 0.262 | 0.994 | 0.817 | 0.550 | 0.215[c] | 0.398 | 0.376 | 0.167 | 0.175 |
| 6VKQ D | 0.235 | 0.262 | 0.988 | 0.816 | 0.547 | 0.211[c] | 0.400 | 0.377 | 0.169 | 0.176 |

| **Fitted onto 7AAA chain B** | **Helix A** | **Helix B** | **Helix D** | **Helix E** | **Helix F** | **Helix G** | **Helix H** | **Helix I** | **Helix J** | **Helix K** |
|---|---|---|---|---|---|---|---|---|---|---|
| *sequence* | *666-677* | *679-688* | *702-722* | *725-740* | *754-779* | *788-797* | *808-821* | *848-853* | *868-876* | *900-906* |
| *# residues* | *12* | *10* | *21* | *16* | *26* | *10* | *14* | *6* | *9* | *7* |
| | (Å) | (Å) | (Å) | (Å) | (Å) | (Å) | (Å) | (Å) | (Å) | (Å) |
| 2PAW A | 0.132 | 0.100 | 0.532 | 0.452 | 0.652 | 0.202 | 0.165 | 0.662 | 0.124 | 0.209 |
| 1A26 A | 0.137 | 0.107 | 0.570 | 0.455 | 0.683 | 0.182 | 0.163 | 0.660 | 0.119 | 0.133 |
| 4DQY C | 0.233 | 0.562 | 0.364 | 0.608 | 0.758 | 0.378 | 0.348 | 0.559 | 0.320 | 0.204 |
| 4DQY F | 0.238 | 0.565 | 0.362 | 0.605 | 0.757 | 0.381 | 0.348 | 0.558 | 0.320 | 0.200 |
| 7AAC A | 0.064 | 0.042 | 0.569 | 1.136 | 0.278 | 0.051 | 0.072 | 0.053 | 0.093 | 0.059 |
| 7AAC B | 0.045 | 0.033 | 0.145 | 0.372 | 0.141 | 0.053 | 0.090 | 0.039 | 0.076 | 0.040 |
| 2RD6 | 0.147 | 0.138 | 0.406 | 0.633 | 0.301 | 0.228 | 0.194 | 0.650 | 0.098 | 0.253 |
| 7AAD A | 0.107 | 0.066 | 0.578 | 0.767 | 0.348 | 0.097 | 0.103 | 0.069 | 0.078 | 0.100 |
| 7AAD B | 0.105 | 0.068 | 0.193 | 0.186 | 0.225 | 0.098 | 0.124 | 0.108 | 0.080 | 0.113 |
| 4PJT A | 0.092 | 0.127 | 0.492[a] | 0.301[b] | 0.627 | 0.116 | 0.185 | 0.155 | 0.129 | 0.116 |
| 4PJT B | 0.090 | 0.079 | 0.209[a] | 0.234[b] | 0.306 | 0.139 | 0.156 | 0.463 | 0.127 | 0.128 |
| 4PJT C | 0.085 | 0.092 | 0.287[a] | 0.235[b] | 0.327 | 0.154 | 0.152 | 0.438 | 0.121 | 0.120 |
| 4PJT D | 0.089 | 0.117 | 0.459[a] | 0.276[b] | 0.526 | 0.122 | 0.146 | 0.150 | 0.122 | 0.124 |
| 4UND A | 0.144 | 0.108 | 0.377 | 0.578 | 0.446 | 0.161 | 0.165 | 0.525 | 0.150 | 0.216 |
| 4UND B | 0.127 | 0.113 | 0.317 | 0.563 | 0.343 | 0.147 | 0.197 | 0.533 | 0.137 | 0.217 |
| 7AAB A | 0.186 | 0.144 | 0.416 | 0.521 | 0.264 | 0.203 | 0.190 | 0.174 | 0.204 | 0.151 |
| 7AAB B | 0.207 | 0.136 | 0.408 | 0.549 | 0.265 | 0.197 | 0.188 | 0.271 | 0.211 | 0.149 |
| 6VKQ A | 0.237 | 0.270 | 1.259 | 1.272 | 0.643 | 0.221[c] | 0.405 | 0.371 | 0.143 | 0.194 |
| 6VKQ B | 0.236 | 0.269 | 1.273 | 1.275 | 0.644 | 0.220[c] | 0.406 | 0.370 | 0.144 | 0.193 |
| 6VKQ C | 0.238 | 0.273 | 1.268 | 1.276 | 0.647 | 0.221[c] | 0.403 | 0.368 | 0.143 | 0.191 |
| 6VKQ D | 0.235 | 0.274 | 1.257 | 1.272 | 0.642 | 0.217[c] | 0.406 | 0.369 | 0.143 | 0.192 |

Only helices with 6 or more residues are shown.
[a] 4PJT uses residues 702-721 for helix D
[b] 4PJT uses residues 730-740 for helix E
[c] 6VKQ uses residues 790-797 for helix G

**Table S2.** Superposition statistics for groups of helices in PARP-1 HD subdomain, demonstrating differences in helical packing (all chains fitted onto 7AAA).

| **Fitted onto 7AAA chain A** | **Helices A and B** | **Helices D and E** | **Helices D and F** | **Helices E and F** | **Helices D, E and F** |
|---|---|---|---|---|---|
| | (Å) | (Å) | (Å) | (Å) | (Å) |
| 2PAW A | 0.278 | 0.563 | 0.549 | 0.599 | 0.599 |
| 1A26 A | 0.302 | 0.586 | 0.594 | 0.630 | 0.635 |
| 4DQY C | 0.621 | 0.696 | 1.294 | 0.936 | 1.197 |
| 4DQY F | 0.625 | 0.696 | 1.292 | 0.934 | 1.196 |
| 7AAC A | 0.043 | 0.115 | 0.131 | 0.123 | 0.131 |
| 7AAC B | 0.105 | 0.653 | 0.638 | 0.440 | 0.683 |
| 2RD6 | 0.372 | 0.496 | 0.419 | 0.415 | 0.476 |
| 7AAD A | 0.107 | 0.180 | 0.216 | 0.200 | 0.209 |
| 7AAD B | 0.107 | 0.577 | 0.525 | 0.375 | 0.564 |
| 4PJT A | 0.460 | 0.367 | 0.867 | 0.619 | 0.802 |
| 4PJT B | 0.308 | 0.488 | 0.477 | 0.361 | 0.477 |
| 4PJT C | 0.307 | 0.466 | 0.496 | 0.333 | 0.471 |
| 4PJT D | 0.332 | 0.413 | 0.619 | 0.542 | 0.588 |
| 4UND A | 0.671 | 0.679 | 0.446 | 0.507 | 0.603 |
| 4UND B | 0.605 | 0.649 | 0.482 | 0.453 | 0.572 |
| 7AAB A | 0.507 | 0.595 | 0.564 | 0.540 | 0.625 |
| 7AAB B | 0.448 | 0.651 | 0.568 | 0.558 | 0.649 |
| 6VKQ A | 0.511 | 0.900 | 0.831 | 0.729 | 0.855 |
| 6VKQ B | 0.514 | 0.897 | 0.830 | 0.727 | 0.853 |
| 6VKQ C | 0.528 | 0.897 | 0.833 | 0.734 | 0.856 |
| 6VKQ D | 0.538 | 0.895 | 0.831 | 0.732 | 0.854 |

| **Fitted onto 7AAA chain B** | **Helices A and B** | **Helices D and E** | **Helices D and F** | **Helices E and F** | **Helices D, E and F** |
|---|---|---|---|---|---|
| | (Å) | (Å) | (Å) | (Å) | (Å) |
| 2PAW A | 0.289 | 0.593 | 0.862 | 0.753 | 0.850 |
| 1A26 A | 0.319 | 0.612 | 0.889 | 0.787 | 0.878 |
| 4DQY C | 0.614 | 0.629 | 1.414 | 0.873 | 1.296 |
| 4DQY F | 0.618 | 0.627 | 1.412 | 0.871 | 1.294 |
| 7AAC A | 0.127 | 0.652 | 0.621 | 0.441 | 0.660 |
| 7AAC B | 0.079 | 0.136 | 0.154 | 0.151 | 0.159 |
| 2RD6 | 0.301 | 0.495 | 0.602 | 0.413 | 0.620 |
| 7AAD A | 0.198 | 0.589 | 0.654 | 0.479 | 0.668 |
| 7AAD B | 0.130 | 0.186 | 0.271 | 0.217 | 0.257 |
| 4PJT A | 0.383 | 0.558 | 1.222 | 0.801 | 1.146 |
| 4PJT B | 0.246 | 0.345 | 0.411 | 0.411 | 0.439 |
| 4PJT C | 0.246 | 0.422 | 0.549 | 0.434 | 0.555 |
| 4PJT D | 0.254 | 0.518 | 0.901 | 0.612 | 0.849 |
| 4UND A | 0.607 | 0.632 | 0.678 | 0.682 | 0.776 |
| 4UND B | 0.526 | 0.412 | 0.429 | 0.451 | 0.496 |
| 7AAB A | 0.434 | 0.477 | 0.380 | 0.567 | 0.511 |
| 7AAB B | 0.380 | 0.504 | 0.348 | 0.569 | 0.511 |
| 6VKQ A | 0.450 | 1.036 | 1.032 | 0.873 | 1.048 |
| 6VKQ B | 0.451 | 1.034 | 1.034 | 0.872 | 1.048 |
| 6VKQ C | 0.465 | 1.035 | 1.036 | 0.879 | 1.052 |
| 6VKQ D | 0.473 | 1.030 | 1.029 | 0.875 | 1.045 |

4PJT uses residues 702-721 for helix D;
4PJT uses residues 730-740 for helix E;
6VKQ uses residues 790-797 for helix G.

**Table S3.** Superposition statistics for individual helices in PARP-1 HD subdomain, comparing different chains in the same structure.

| **7AAA** | **Helix A** | **Helix B** | **Helix D** | **Helix E** | **Helix F** | **Helix G** | **Helix H** | **Helix I** | **Helix J** | **Helix K** |
|---|---|---|---|---|---|---|---|---|---|---|
| *sequence* | *666-677* | *679-688* | *702-722* | *725-740* | *754-779* | *788-797* | *808-821* | *848-853* | *868-876* | *900-906* |
| *# residues* | *12* | *10* | *21* | *16* | *26* | *10* | *14* | *6* | *9* | *7* |
| | (Å) | (Å) | (Å) | (Å) | (Å) | (Å) | (Å) | (Å) | (Å) | (Å) |
| A vs B | 0.052 | 0.029 | 0.550 | 1.197 | 0.289 | 0.044 | 0.016 | 0.016 | 0.101 | 0.045 |

| **4DQY** | **Helix A** | **Helix B** | **Helix D** | **Helix E** | **Helix F** | **Helix G** | **Helix H** | **Helix I** | **Helix J** | **Helix K** |
|---|---|---|---|---|---|---|---|---|---|---|
| *sequence* | *666-677* | *679-688* | *702-722* | *725-740* | *754-779* | *788-797* | *808-821* | *848-853* | *868-876* | *900-906* |
| *# residues* | *12* | *10* | *21* | *16* | *26* | *10* | *14* | *6* | *9* | *7* |
| | (Å) | (Å) | (Å) | (Å) | (Å) | (Å) | (Å) | (Å) | (Å) | (Å) |
| A vs B | 0.013 | 0.014 | 0.015 | 0.021 | 0.018 | 0.010 | 0.016 | 0.010 | 0.014 | 0.018 |

| **7AAC** | **Helix A** | **Helix B** | **Helix D** | **Helix E** | **Helix F** | **Helix G** | **Helix H** | **Helix I** | **Helix J** | **Helix K** |
|---|---|---|---|---|---|---|---|---|---|---|
| *sequence* | *666-677* | *679-688* | *702-722* | *725-740* | *754-779* | *788-797* | *808-821* | *848-853* | *868-876* | *900-906* |
| *# residues* | *12* | *10* | *21* | *16* | *26* | *10* | *14* | *6* | *9* | *7* |
| | (Å) | (Å) | (Å) | (Å) | (Å) | (Å) | (Å) | (Å) | (Å) | (Å) |
| A vs B | 0.057 | 0.029 | 0.532 | 1.074 | 0.195 | 0.044 | 0.037 | 0.025 | 0.031 | 0.038 |

| **7AAD** | **Helix A** | **Helix B** | **Helix D** | **Helix E** | **Helix F** | **Helix G** | **Helix H** | **Helix I** | **Helix J** | **Helix K** |
|---|---|---|---|---|---|---|---|---|---|---|
| *sequence* | *666-677* | *679-688* | *702-722* | *725-740* | *754-779* | *788-797* | *808-821* | *848-853* | *868-876* | *900-906* |
| *# residues* | *12* | *10* | *21* | *16* | *26* | *10* | *14* | *6* | *9* | *7* |
| | (Å) | (Å) | (Å) | (Å) | (Å) | (Å) | (Å) | (Å) | (Å) | (Å) |
| A vs B | 0.092 | 0.059 | 0.504 | 0.706 | 0.223 | 0.057 | 0.036 | 0.077 | 0.015 | 0.066 |

| **4PJT** | **Helix A** | **Helix B** | **Helix D** | **Helix E** | **Helix F** | **Helix G** | **Helix H** | **Helix I** | **Helix J** | **Helix K** |
|---|---|---|---|---|---|---|---|---|---|---|
| *sequence* | *666-677* | *679-688* | *702-721* | *730-740* | *754-779* | *788-797* | *808-821* | *848-853* | *868-876* | *900-906* |
| *# residues* | *12* | *10* | *21* | *16* | *26* | *10* | *14* | *6* | *9* | *7* |
| | (Å) | (Å) | (Å) | (Å) | (Å) | (Å) | (Å) | (Å) | (Å) | (Å) |
| A vs B | 0.042 | 0.076 | 0.387 | 0.337 | 0.483 | 0.051 | 0.056 | 0.359 | 0.040 | 0.026 |
| A vs C | 0.044 | 0.101 | 0.303 | 0.092 | 0.395 | 0.064 | 0.066 | 0.332 | 0.036 | 0.040 |
| A vs D | 0.037 | 0.089 | 0.126 | 0.164 | 0.247 | 0.036 | 0.071 | 0.030 | 0.036 | 0.026 |
| B vs C | 0.036 | 0.069 | 0.126 | 0.058 | 0.113 | 0.058 | 0.049 | 0.083 | 0.036 | 0.036 |
| B vs D | 0.028 | 0.087 | 0.350 | 0.162 | 0.414 | 0.056 | 0.054 | 0.359 | 0.054 | 0.023 |
| C vs D | 0.029 | 0.051 | 0.260 | 0.161 | 0.328 | 0.061 | 0.036 | 0.335 | 0.038 | 0.027 |

4PJT uses residues 702-721 for helix D;
4PJT uses residues 730-740 for helix E;

| **4UND** | **Helix A** | **Helix B** | **Helix D** | **Helix E** | **Helix F** | **Helix G** | **Helix H** | **Helix I** | **Helix J** | **Helix K** |
|---|---|---|---|---|---|---|---|---|---|---|
| *sequence* | *666-677* | *679-688* | *702-722* | *725-740* | *754-779* | *788-797* | *808-821* | *848-853* | *868-876* | *900-906* |
| *# residues* | *12* | *10* | *21* | *16* | *26* | *10* | *14* | *6* | *9* | *7* |
| | (Å) | (Å) | (Å) | (Å) | (Å) | (Å) | (Å) | (Å) | (Å) | (Å) |
| A vs B | 0.056 | 0.086 | 0.305 | 0.245 | 0.368 | 0.046 | 0.100 | 0.019 | 0.058 | 0.036 |

| **7AAB** | **Helix A** | **Helix B** | **Helix D** | **Helix E** | **Helix F** | **Helix G** | **Helix H** | **Helix I** | **Helix J** | **Helix K** |
|---|---|---|---|---|---|---|---|---|---|---|
| *sequence* | *666-677* | *679-688* | *702-722* | *725-740* | *754-779* | *788-797* | *808-821* | *848-853* | *868-876* | *900-906* |
| *# residues* | *12* | *10* | *21* | *16* | *26* | *10* | *14* | *6* | *9* | *7* |
| | (Å) | (Å) | (Å) | (Å) | (Å) | (Å) | (Å) | (Å) | (Å) | (Å) |
| A vs B | 0.100 | 0.028 | 0.120 | 0.194 | 0.137 | 0.037 | 0.020 | 0.288 | 0.046 | 0.029 |

**Table S3 (continued).** Superposition statistics for individual helices in PARP-1 HD subdomain, comparing different chains in the same structure.

| **6VKQ** | **Helix A** | **Helix B** | **Helix D** | **Helix E** | **Helix F** | **Helix G** | **Helix H** | **Helix I** | **Helix J** | **Helix K** |
|---|---|---|---|---|---|---|---|---|---|---|
| *sequence* | *666-677* | *679-688* | *702-722* | *725-740* | *754-779* | *790-797* | *808-821* | *848-853* | *868-876* | *900-906* |
| *# residues* | *12* | *10* | *21* | *16* | *26* | *10* | *14* | *6* | *9* | *7* |
| | (Å) | (Å) | (Å) | (Å) | (Å) | (Å) | (Å) | (Å) | (Å) | (Å) |
| A vs B | 0.007 | 0.006 | 0.025 | 0.006 | 0.010 | 0.005 | 0.009 | 0.005 | 0.005 | 0.003 |
| A vs C | 0.008 | 0.012 | 0.018 | 0.007 | 0.013 | 0.004 | 0.013 | 0.011 | 0.003 | 0.008 |
| A vs D | 0.010 | 0.016 | 0.011 | 0.008 | 0.015 | 0.009 | 0.012 | 0.005 | 0.007 | 0.008 |
| B vs C | 0.011 | 0.011 | 0.012 | 0.008 | 0.015 | 0.007 | 0.012 | 0.008 | 0.005 | 0.007 |
| B vs D | 0.011 | 0.014 | 0.022 | 0.009 | 0.018 | 0.011 | 0.013 | 0.007 | 0.008 | 0.006 |
| C vs D | 0.005 | 0.006 | 0.015 | 0.009 | 0.009 | 0.010 | 0.007 | 0.009 | 0.007 | 0.004 |

6VKQ uses residues 790-797 for helix G.

**Table S4.** Superposition statistics for the HD and ART subdomains of PARP-1 CAT domain (all chains onto 7AAA).

| Fitted atoms (set 1) | Resolution | Space group | CAT | ART | HD | ART | HD |
|---|---|---|---|---|---|---|---|
| Measured atoms (set 2) | | | CAT | ART | HD | HD | ART |
| (all fitted onto 7AAA chain A) | (Å) | | (Å) | (Å) | (Å) | (Å) | (Å) |
| 2PAW A (apo-protein) | 2.30 | $P\ 2_12_12_1$ | 0.679 | 0.568 | 0.630 | 1.041 | 0.976 |
| 1A26 A (apo-protein) | 2.25 | $P\ 2_12_12_1$ | 0.681 | 0.496 | 0.646 | 1.189 | 1.033 |
| 4DQY C (apo-protein) | 3.25 | $P\ 2_12_12_1$ | 1.293 | 0.804 | 1.566 | 2.233 | 2.328 |
| 4DQY F (apo-protein) | " | " | 1.294 | 0.806 | 1.565 | 2.234 | 2.340 |
| 7AAC A (veliparib complex) | 1.59 | $P\ 2_12_12_1$ | 0.145 | 0.133 | 0.120 | 0.181 | 0.389 |
| 7AAC B (veliparib complex) | " | " | 0.494 | 0.222 | 0.697 | 0.952 | 0.885 |
| 2RD6 A (veliparib complex) | 2.30 | P 3 2 1 | 0.980 | 0.855 | 0.541 | 1.720 | 1.550 |
| 7AAD A (olaparib complex) | 2.21 | $P\ 2_12_12_1$ | 0.241 | 0.231 | 0.220 | 0.272 | 0.456 |
| 7AAD B (olaparib complex) | " | " | 0.457 | 0.309 | 0.625 | 0.701 | 0.679 |
| 4UND A (talazoparib complex) | 2.20 | $P\ 2_12_12_1$ | 1.174 | 0.760 | 0.850 | 2.429 | 2.855 |
| 4UND B (talazoparib complex) | " | " | 1.152 | 0.855 | 0.823 | 2.140 | 2.877 |
| 4PJT A (talazoparib complex) | 2.35 | $P\ 2_12_12_1$ | 1.185 | 0.690 | 0.847 | 2.574 | 3.042 |
| 4PJT B (talazoparib complex) | " | " | 0.932 | 0.667 | 0.546 | 1.942 | 1.575 |
| 4PJT C (talazoparib complex) | " | " | 0.952 | 0.716 | 0.528 | 1.930 | 1.783 |
| 4PJT D (talazoparib complex) | " | " | 1.044 | 0.664 | 0.653 | 2.311 | 2.177 |
| 7AAB A (EB-47 complex) | 2.8 | $P\ 6_1$ | 1.789 | 0.506 | 0.969 | 4.175 | 4.362 |
| 7AAB B (EB-47 complex) | " | " | 1.825 | 0.509 | 0.992 | 4.196 | 4.747 |
| 6VKQ A (EB-47 complex) | 2.9 | $P\ 4_12_12$ | 1.421 | 0.771 | 0.997 | 3.166 | 2.763 |
| 6VKQ B (EB-47 complex) | " | " | 1.421 | 0.770 | 0.990 | 3.171 | 2.782 |
| 6VKQ C (EB-47 complex) | " | " | 1.432 | 0.780 | 1.003 | 3.181 | 2.829 |
| 6VKQ D (EB-47 complex) | " | " | 1.448 | 0.780 | 1.003 | 3.217 | 2.870 |

| Fitted atoms (set 1) | Resolution | Space group | CAT | ART | HD | ART | HD |
|---|---|---|---|---|---|---|---|
| Measured atoms (set 2) | | | CAT | ART | HD | HD | ART |
| (all fitted onto 7AAA chain B) | (Å) | | (Å) | (Å) | (Å) | (Å) | (Å) |
| 2PAW A (apo-protein) | 2.30 | $P\ 2_12_12_1$ | 0.848 | 0.599 | 0.962 | 1.431 | 1.543 |
| 1A26 A (apo-protein) | 2.25 | $P\ 2_12_12_1$ | 0.855 | 0.530 | 0.960 | 1.581 | 1.683 |
| 4DQY C (apo-protein) | 3.25 | $P\ 2_12_12_1$ | 1.465 | 0.854 | 1.741 | 2.584 | 3.368 |
| 4DQY F (apo-protein) | " | " | 1.466 | 0.855 | 1.741 | 2.585 | 3.380 |
| 7AAC A (veliparib complex) | 1.59 | $P\ 2_12_12_1$ | 0.529 | 0.199 | 0.710 | 1.021 | 1.459 |
| 7AAC B (veliparib complex) | " | " | 0.189 | 0.180 | 0.154 | 0.228 | 0.346 |
| 2RD6 A (veliparib complex) | 2.30 | P 3 2 1 | 1.034 | 0.863 | 0.635 | 1.770 | 2.238 |
| 7AAD A (olaparib complex) | 2.21 | $P\ 2_12_12_1$ | 0.590 | 0.286 | 0.765 | 1.116 | 1.504 |
| 7AAD B (olaparib complex) | " | " | 0.355 | 0.327 | 0.270 | 0.556 | 0.635 |
| 4UND A (talazoparib complex) | 2.20 | $P\ 2_12_12_1$ | 1.298 | 0.745 | 0.913 | 2.789 | 3.841 |
| 4UND B (talazoparib complex) | " | " | 1.193 | 0.842 | 0.658 | 2.210 | 3.863 |
| 4PJT A (talazoparib complex) | 2.35 | $P\ 2_12_12_1$ | 1.366 | 0.702 | 1.165 | 2.868 | 4.070 |
| 4PJT B (talazoparib complex) | " | " | 1.029 | 0.675 | 0.627 | 2.227 | 2.253 |
| 4PJT C (talazoparib complex) | " | " | 1.055 | 0.727 | 0.606 | 2.194 | 2.614 |
| 4PJT D (talazoparib complex) | " | " | 1.146 | 0.680 | 0.845 | 2.400 | 3.077 |
| 7AAB A (EB-47 complex) | 2.8 | $P\ 6_1$ | 1.802 | 0.551 | 0.910 | 4.263 | 4.095 |
| 7AAB B (EB-47 complex) | " | " | 1.823 | 0.554 | 0.933 | 4.243 | 4.373 |
| 6VKQ A (EB-47 complex) | 2.9 | $P\ 4_12_12$ | 1.538 | 0.778 | 1.192 | 3.425 | 3.445 |
| 6VKQ B (EB-47 complex) | " | " | 1.540 | 0.778 | 1.185 | 3.437 | 3.479 |
| 6VKQ C (EB-47 complex) | " | " | 1.550 | 0.787 | 1.198 | 3.440 | 3.542 |
| 6VKQ D (EB-47 complex) | " | " | 1.562 | 0.786 | 1.194 | 3.466 | 3.569 |

The atoms used for superposition were as follows:
HD = 666-721, 730-743, 750-779 (N,$C^{\alpha}$,C');
ART = 790-936, 939-1009 (N,$C^{\alpha}$,C');
Set 1 atoms are used to calculate transformed co-ordinates resulting from fitting;
Set 2 atoms are used to calculate an rms difference without further changing the co-ordinates.

**Table S5.** Superposition statistics for the HD and ART subdomains of PARP-1 CAT domain between different complexes with the same inhibitor (all chains).

| Fitted atoms (set 1) | CAT | ART | HD | ART | HD |
|---|---|---|---|---|---|
| **Measured atoms (set 2)** | **CAT** | **ART** | **HD** | **HD** | **ART** |
| | (Å) | (Å) | (Å) | (Å) | (Å) |
| **Comparing veliparib complexes** | | | | | |
| 7AAC A vs 2RD6 A | 0.983 | 0.874 | 0.518 | 1.710 | 1.442 |
| 7AAC B vs 2RD6 A | 1.040 | 0.869 | 0.648 | 1.758 | 2.091 |
| **Comparing talazoparib complexes** | | | | | |
| 4PJT A vs 4UND A | 0.955 | 0.712 | 0.867 | 1.640 | 1.429 |
| 4PJT A vs 4UND B | 0.653 | 0.542 | 0.612 | 0.908 | 1.957 |
| 4PJT A vs 4UND C | 0.653 | 0.574 | 0.588 | 0.921 | 1.495 |
| 4PJT A vs 4UND D | 0.961 | 0.680 | 0.782 | 1.855 | 1.616 |
| 4PJT B vs 4UND A | 0.822 | 0.695 | 0.944 | 1.174 | 0.867 |
| 4PJT B vs 4UND B | 0.787 | 0.620 | 0.584 | 1.310 | 2.204 |
| 4PJT B vs 4UND C | 0.685 | 0.608 | 0.531 | 1.016 | 1.641 |
| 4PJT B vs 4UND D | 0.725 | 0.671 | 0.699 | 0.913 | 1.184 |
| **Comparing EB-47 complexes** | | | | | |
| 7AAB A vs 6VKQ A | 1.087 | 0.645 | 0.898 | 1.839 | 3.840 |
| 7AAB A vs 6VKQ B | 1.097 | 0.645 | 0.898 | 1.859 | 3.897 |
| 7AAB A vs 6VKQ C | 1.104 | 0.649 | 0.909 | 1.867 | 3.992 |
| 7AAB A vs 6VKQ D | 1.088 | 0.651 | 0.902 | 1.826 | 3.943 |
| 7AAB B vs 6VKQ A | 1.160 | 0.645 | 0.935 | 1.976 | 4.430 |
| 7AAB B vs 6VKQ B | 1.170 | 0.645 | 0.935 | 2.000 | 4.487 |
| 7AAB B vs 6VKQ C | 1.178 | 0.648 | 0.949 | 2.006 | 4.584 |
| 7AAB B vs 6VKQ D | 1.161 | 0.650 | 0.942 | 1.963 | 4.536 |

The atoms used for superposition were as follows:
HD = 666-721, 730-743, 750-779 (N,$C^{\alpha}$,C');
ART = 790-936, 939-1009 (N,$C^{\alpha}$,C');
Set 1 atoms are used to calculate transformed co-ordinates resulting from fitting;
Set 2 atoms are used to calculate an rms difference without further changing the co-ordinates.
